# The 3-Dimensional Genome Drives the Evolution of Asymmetric Gene Duplicates via Enhancer Capture-Divergence

**DOI:** 10.1101/2022.11.30.518413

**Authors:** UnJin Lee, Deanna Arsala, Shengqian Xia, Cong Li, Mujahid Ali, Nicolas Svetec, Christopher B Langer, Débora R. Sobreira, Ittai Eres, Dylan Sosa, Jianhai Chen, Li Zhang, Patrick Reilly, Alexander Guzzetta, J.J. Emerson, Peter Andolfatto, Qi Zhou, Li Zhao, Manyuan Long

**Author notes:** These authors contributed equally to this work (UnJin Lee, Deanna Arsala, Shengqian Xia).

## Abstract

Previous evolutionary models of duplicate gene evolution have overlooked the pivotal role of genome architecture. Here, we show that proximity-based regulatory recruitment of distally duplicated genes (enhancer capture) is an efficient mechanism for modulating tissue-specific production of pre-existing proteins. By leveraging genomic asymmetries in synteny and function that distinguish new genes evolving under enhancer capture-divergence (ECD) from those evolving under previous models, we performed a co-expression analysis on *Drosophila melanogaster* tissue data to show the generality of ECD as a significant evolutionary driver of asymmetric, distally duplicated genes. We use the recently evolved gene *HP6*/*Umbrea*, which duplicated <15 million years ago (mya), as an example of the ECD process. By assaying genome-wide chromosomal conformations in multiple *Drosophila* species, we show that *HP6/Umbrea* was inserted into a pre-existing, evolutionarily stable 3D genomic structure spanning over 125kb. We then utilize this data to identify a newly discovered enhancer (FLEE1), buried within the coding region of the highly conserved, essential gene *MFS18*, that likely neo-functionalized *HP6/Umbrea*, thereby driving the new duplicate gene copy to fixation. Finally, we demonstrate ancestral transcriptional co-regulation of *HP6/Umbrea*’s future insertion site using single-cell transcriptomics, illustrating how enhancer capture provides a highly evolvable, one-step solution to Ohno’s Dilemma. The intuitive molecular mechanism underpinning the ECD model unveils a novel and robust framework to understand the fixation and neofunctionalization of distally duplicated genes.

## Main Text

Newly duplicated genes are at risk of loss in a population through genetic drift or negative selection (*1*) before rare, advantageous neo-functionalizing mutations may occur. The probability of fixation for a slightly deleterious duplicate is more than one to two orders of magnitude lower than a neutral mutant (P_fix_ = 0.085∼0.003 x 1/(2N_e_), Ne ≈ 10^6^ as the effective population) for complete and exon duplicates in *Drosophila melanogaster* (*1, 2*). Consequently, the vast majority of non-fixed duplicate gene copies are likely to be lost in approximately just 2.32 generations or less (**METHODS AND MATERIALS**). Given the low mutation rates in this species (*2, 3*), it is hardly impossible for a newly fixed duplicate gene to fix or even acquire new mutations, let alone an advantageous one, in such a short time. This problem has also been previously referred to as “Ohno’s dilemma” (*4*). Various models have been proposed to resolve this problem including: the duplication, divergence, complementation (DDC)/sub-functionalization model (*5*), the escape from adaptive conflict (EAC) model (*6*), the innovation, amplification, and divergence (IAD) model (*4, 7*).

The DDC model, also known as subfunctionalization, represents a neutral evolutionary process where symmetric (identical) gene duplicates lose different aspects of their original function due to genetic drift. This random divergence results in the preservation of the duplicated genes, each retaining distinct, yet complementary, functions (Figure 1). Conversely, genes evolving under the EAC model, are under selection for enhanced optimization of specific functions originally held by the parental gene that are partitioned to paralogous copies. The EAC model posits that a single parental gene has intrinsic genetic conflict due to its inability to optimize multiple functions simultaneously, and gene duplication can resolve this evolutionary constraint. While the DDC and EAC models explain how ancestral functions are partitioned among gene duplicates, they fall short in explaining the immediate development of novel expression patterns following gene duplication. These novel expression profiles, often resulting from the gene’s new genomic context, can be instrumental in driving the evolution of new functions—processes not fully captured by the DDC and EAC models. In contrast, the IAD model describes how shifts in selection pressures can promote the expression of genes with auxiliary functions by increasing gene copy number (Figure 1). Following the initial increase of auxiliary function through gene amplification, subsequent relaxation of selection pressure will allow for changes to accumulate on the various copies, allowing the new copies to diverge and potentially gain a new function (*4*).

**Figure 1.**
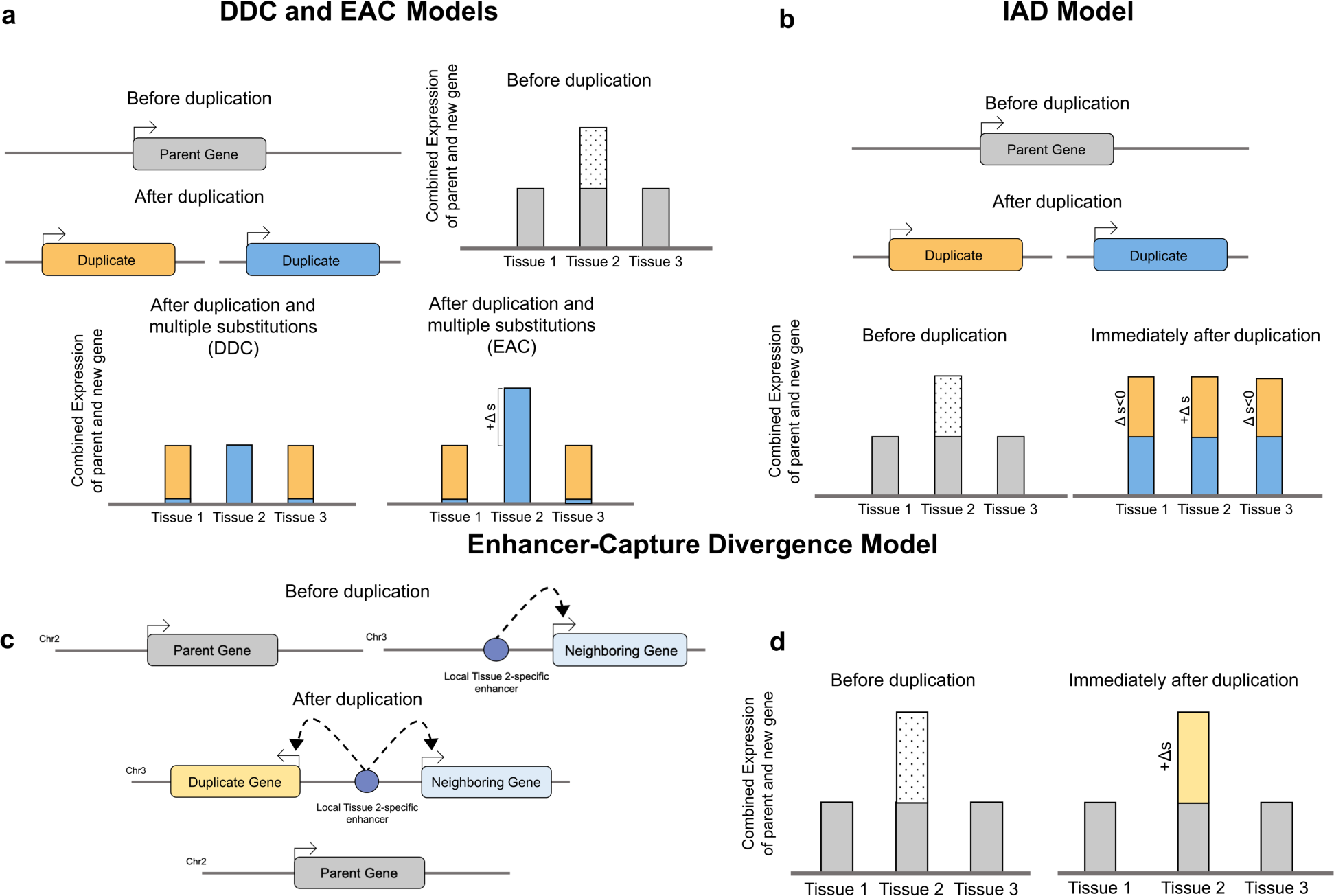
Comparison of extant models. Presented are illustrations for the (a) Duplication-Divergence-Complementation (DDC)/Escape-from-Adaptive-Conflict (EAC), the (b) Innovation-Amplification-Divergence (IAD), and the (c, d) Enhancer Capture-Divergence (ECD) models of duplicate gene evolution, where the gene regulation of three tissue types are considered. In this scenario, we assume no protein substitutions, so all duplicate gene copies produce identical proteins. Dotted box represents selection for increased expression, and Δs indicates the change in selection coefficient. Under the (a) DDC and EAC models, a parental gene duplicates and causes redundancy in the genome. In the DDC model, redundancy allows for compensation of any single loss-of-function event, eventually causing the expression pattern of the ancestral gene to be segregated between both new gene copies in a complementary fashion. Assuming no protein-coding changes, the total output of duplicate gene copies is identical to the original gene – therefore the DDC model is a neutrally evolving process. In the EAC model, increased production in a specific tissue cannot occur within a single gene copy due to internal conflict. This conflict is resolved via the act of duplication, where functions are segregated between duplicate gene copies, allowing the output of these two genes to increase fitness. Note that the identity of duplicate gene copies may not be distinguished under the EAC and DDC models (symmetric), resulting in a random segregation of function. Under the (b) IAD model, an ancestral gene duplicates to increase production of the original protein in a single step. This increased dosage can potentially cause deleterious effects via misexpression or over-activity in multi-cellular organisms. Note that the identity of duplicate gene copies also cannot be distinguished in the IAD model (symmetric), resulting in a redundant segregation of function. Under the (c) ECD model, a parental gene fully duplicates into a distant region of the genome that is under the control of a pre-existing enhancer. By capturing this new interaction (d), this duplication increases tissue-specific production of the original protein in a single step. Notice that the clearly identifiable parental gene copy remains unaltered and thus all original function is retained, while the duplicate copy acts only to increase protein expression in a single tissue.

While the IAD model provides a reasonable explanation for gene family expansions in microbial organisms while encountering environmental changes (*7*), the model faces serious problems when applied to metazoans, as a general and broad increase in gene dosage may be advantageous in some cell or tissue types but potentially deleterious in others. Similar to the EAC and DDC models, the IAD model does not directly explain how novel expression patterns arise immediately following gene duplication, leaving a gap in our understanding of duplicate gene evolution.

We propose the enhancer capture-divergence (ECD) model, which is an evolutionary model produced by asymmetric RNA or DNA-based gene duplication processes that allow for distinct parental and new gene identities and functions (Figure 1). The ECD model first proposes that selective pressures change for the increased expression of a pre-existing (parental) gene within a specific tissue or set of tissues. While the evolution of a new enhancer in the parental gene’s locus is plausible, it would require multiple neutral *de novo* substitutions or insertions to generate one or more necessary transcription factor binding sites that fix within a population and modulates the expression of the new gene duplicate without disrupting parent gene’s expression pattern. Under the ECD model, duplication of the parental gene into another regulatory environment under the control of a pre-existing, tissue-specific enhancer is a solution that requires far fewer genomic changes and can occur in a single step. As the new selection pressures recur, the duplicate copy that is under new regulatory control will increase in frequency in the population, allowing it to fix. If the selection pressures change such that the increased tissue-specific expression of the new gene is no longer advantageous or compensatory mutations appear in the original parent locus, selective pressures will relax on the new gene copy allowing for divergence. While loss of the new gene copy by drift or negative selection is one possible fate, if the duplicate gene copy is at high enough frequency within a population, substitutions may accumulate and result in the gain of new, tissue-specific protein function.

The previous models addressing Ohno’s Dilemma (DDC, EAC, IAD) are symmetric models of duplication-based evolution which assume that the original parental gene function is randomly partitioned or entirely retained between identical duplicate copies, making parent and new gene copies indistinguishable from one another. They offer plausible mechanisms for the retention of certain types of gene duplicates in various processes of subfunctionalization, conflict resolution and amplification of ancestral gene functions. The ECD offers a 3D-facilitated efficient route of neofunctionalization than the development of new enhancers from scratch. It also provides a coherent and testable framework that describes the evolution of asymmetric or distal gene duplicates that is currently unexplained by previous models. The asymmetry is a key feature of the ECD model that distinguishes it from the DDC, EAC and IAC models in different consequences of functional evolution, and allows for clear identification of genes that evolved under enhancer capture. (An extended discussion of prior models and genomic symmetry is available in **SUPPLEMENTARY INFORMATION**).

## RESULTS

### Analysis of Tissue Co-Expression Reveals New Genes Evolve by Enhancer Capture

Central to the IAD model is the observation that gene duplication via unequal crossing over is more likely to occur than a point mutation (*4, 7*). As previously described, one issue with this model is that there is an implicit assumption that during the environmental shift, the increase in fitness gained by over-activity of the auxiliary function must be greater than the decrease in fitness imparted by over-activity of the original gene’s function(s). In the case of the enzymatic activity of single-celled organisms where environments are encountered sequentially, it is reasonable to assume that selection might tolerate over-activity of the gene’s original function during the transient environment in which the auxiliary function is favored. However, the decrease in fitness for improper expression or activity is larger in multicellular organisms than in single-celled organisms, where a multi-cellular organism’s overall phenotype is the cumulative (development) and simultaneous (organ systems) product of many different gene functions.

In the case of multicellular organisms, selection may increase the expression of a gene within a single tissue type (Figure 1). Under the IAD model, a full duplication of the parent gene function and expression pattern drives the duplicate copies to fixation as it provides the most evolvable solution to new conditions. In contrast, under the enhancer capture-divergence model, a copy of the parent gene duplicates into a region of the genome containing an active enhancer(s) that modulates the new gene copy’s expression in a tissue-specific manner. Alternatively, the new gene may duplicate into an inactive region of the genome containing unbound transcription factor binding sites, thus activating a previously inert non-coding sequence into a *de novo* enhancer.

Compared to the tissue-specific nature of genes evolving under the ECD model, genes evolving under the IAD model are over-expressed in all tissues, as they are assumed to take on the parent gene expression pattern. We therefore predict that enhancer capture will be more dominant than the IAD model for asymmetrically duplicated genes within multicellular organisms, as it avoids the potentially deleterious effects of increased dosage in multiple tissues resulting from full duplication. However, we stress that the IAD model is likely to drive the evolution of a large number of tandem duplicates as well as a subset of asymmetrically duplicates where the recruitment of pre-existing regulatory elements is unlikely. This increase in fitness caused by the combined output of the new and parental genes thus drives the new gene copy to fixation, providing an alternate resolution to Ohno’s Dilemma than the IAD model. Once the tissue-specific selection for the new gene is relaxed, the new gene may then begin to diverge, accumulating substitutions.

Some classes of new genes will continue to evolve under the IAD, DDC, and EAC models. However, the relationships of new genes with their parent genes and neighboring genes differ in expression between those evolving under those previous models and our ECD model, allowing for direct testing of the ECD mechanism as a driver of newly evolved genes. Under the DDC or EAC models, the tissue expression patterns of parental and new genes are complementary, resulting in low co-expression between parental and new gene copies (“parental co-expression”). Since new gene evolution under the DDC and EAC models is assumed to occur in a regulatory-independent context, the tissue expression patterns of the new gene and its neighboring genes should have no relationship, resulting in random co-expression between the new gene and its neighboring gene (“neighboring co-expression”). Under the IAD model, genes and their upstream regulatory sequences are fully duplicated, which predicts a high co-expression between the parent and new gene copies, while the new gene copy and its neighboring genes should have low co-expression. In the enhancer capture-divergence model, the parent gene is predicted to be more broadly expressed, while the new gene which resides in a distant region of the genome is under the control of one or more tissue-specific enhancers. Here, parental genes are expected to have broad tissue expression patterns, while new genes have expression patterns with high tissue specificity, resulting in low parental co-expression. On the other hand, since the new gene becomes regulated by a locally captured enhancer that is already influencing other genes, neighboring co-expression is high, particularly in gene-dense genomes.

To determine whether the enhancer capture-divergence process is a significant driver of new gene evolution in *D. melanogaster*, we obtained tissue expression data from FlyBase (*8, 9*)(c.f. Methods and Materials) and calculated co-expression between new/parental and new/neighboring gene pairs (Spearman correlation coefficient) for a random subset of newly evolved genes that 1) underwent a duplication into a different topologically associating domain (TAD) than its parental gene (as defined in (*10*)) and 2) whose essentiality has been validated experimentally ((N=87, Supp. Table S1, Methods and Materials). We focused on experimentally validated genes that were in a different TAD, using distal duplications as a proxy, as their asymmetry allowed us to definitively identify the parent and new gene copies via synteny. To calculate co-expression, we used expression data that contained tissue types extracted from both L3 larvae, pre-pupae, and adult flies, including gut, salivary glands, and imaginal discs from wandering L3 larvae, as well as the head, ovaries, gut, and reproductive organs from adults (c.f. Methods and Materials). For tissues that were represented with multiple experimental runs, data from those tissue types were averaged prior to further analyses to avoid representation bias.

To determine whether a significant number of distally duplicated (non-tandem) genes evolved by enhancer capture, we used two concurrent features of the new genes in our dataset (parent/neighbor tissue co-expression (“PNC”) and essentiality) that together determine whether the ECD process is a significant driver of new gene evolution alongside other established models: (Figure 2b). We define “low” and “high” co-expression as being below or above the median co-expression value across new genes in our data set. Under the symmetrical DDC, EAC, and IAD models, the parent and new gene copies are indistinguishable in that all segregable and essential functions of the original gene partition randomly between both parent and duplicate gene copies. In contrast, the ECD model predicts that all original functions, including essential function, are expected to remain with the parental gene while the new copy retains an auxiliary non-essential function. Thus, genes which evolved under enhancer capture are expected to be disproportionately enriched for non-essential functions. Furthermore, as genes under ECD are duplicated into a different regulatory environment from that of their parents, they are expected to appear in the lower right quadrant (quadrant IV) in the PNC plots, with high neighboring co-expression and low parental co-expression (Figure 2b, 2c). In contrast, genes with that have evolved symmetrically via the DDC or EAC models are expected to have a random partitioning of all functions (including essential functions) and should appear in the bottom half of the PNC plots (quadrants III and IV). Specifically, genes evolving under these processes are expected to have low parental co-expression resulting from divergent and complementary expression patterns, while the absence of regulatory context in the DDC and EAC models result in a prediction of random neighboring co-expression, as there is no expected relationship between the new gene and its neighboring genes (Figure 2b, 2c). Similarly, genes that have evolved via the IAD model should also have a random partitioning of essential functions while also appearing in the upper half of the PNC plots (quadrants I and II), with high parental co-expression resulting from full duplication (Figure 2b, 2c).

**Figure 2.**
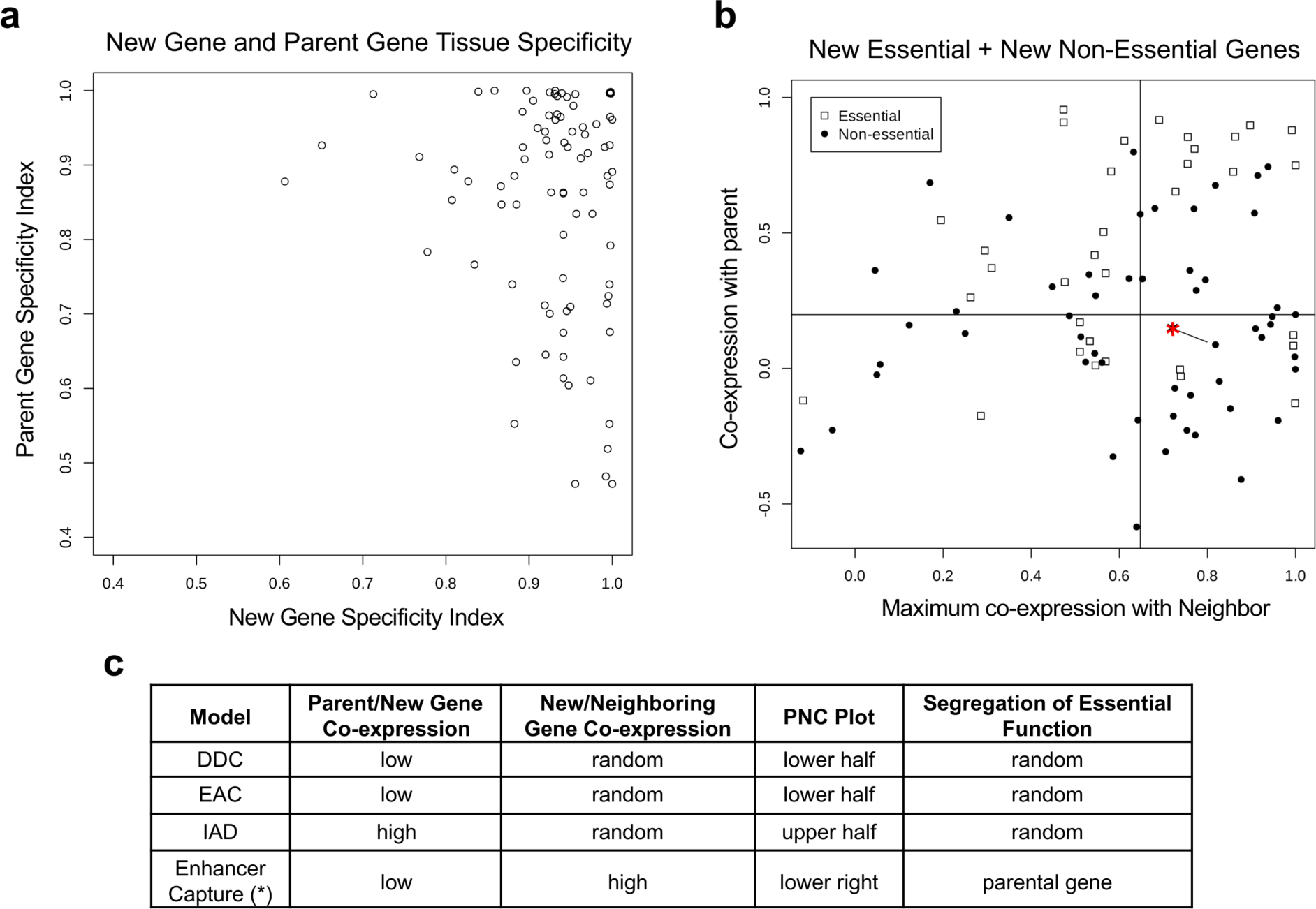
Asymmetrically duplicated new genes evolve via enhancer capture. (a) Using new-gene/parent-gene pairs for genes evolving via distal duplication in *D. melanogaster*, the tissue specificity index *τ* is calculated and plotted above, demonstrating that new genes evolving via distal duplication have higher tissue specificity than parental genes. (b) Shown are parent/neighbor tissue co-expression patterns for new genes in *D. melanogaster* which have duplicated either more than 500kb away or between chromosomes. Tissue co-expression (Spearman correlation coefficient) between new gene/parental gene pairs is plotted on the vertical axis while maximal tissue co-expression between new gene/neighboring genes pairs is plotted on the horizontal axis. Vertical and horizontal lines indicate median co-expression value of all distally duplicated new genes presented here. Genes that evolved via enhancer capture are expected to have low parental co-expression and high neighboring co-expression and should thus be present in the lower right quadrant. (c) While a new gene’s essential function is equally likely to be partitioned between either parent or new gene under prior models, new genes evolving via enhancer capture are unlikely to have essential functions, as the expression of the new gene will only augment existing expression of the parental gene, leaving the original essential function intact. Comparing the ratio of new essential to new non-essential genes (29:36, 44.6%) in quadrants I-III to the ratio of new essential to new non-essential genes in quadrant IV showing high neighboring/low parental co-expression (5:17, 22.7%) shows that new genes likely evolve via regulatory capture (2.18 fold enrichment, p=0.0055, 2D K-S test). (* denotes *HP6/Umbrea*)

Alternatively, the enhancer capture-divergence model predicts that most function, including essential gene function, will remain with the parental gene copy, while the tissue-specific expression pattern of the duplicate gene copy serves only to augment the function of the parental gene – a pattern frequently seen in new genes evolving via distal duplication (Figure 2a). Specifically, selection for increased tissue-specific expression of the parental gene predicts the appearance of a distal duplicate of the parental gene copy both with non-essential function and high neighboring co-expression. Meanwhile, the expression pattern and gene function – including all essential function – of the parental copy remains unaltered and is retained. Together, the enhancer capture-divergence model predicts a combination of high neighboring co-expression, low parental co-expression, and non-essentiality. This prediction may be tested by looking for a statistical enrichment of non-essential genes in the lower right quadrant of the PNC plot relative to background (Figure 2b, Supp. Table S1). A distortion in the segregation of essential function is readily identified using the parent/neighbor co-expression plots for distally-duplicated genes in *D. melanogaster*, where the ratio of new essential:new non-essential genes in the lower right quadrant (5:17, 22.7%) was found to be significantly lower than the ratio of remaining new essential:new non-essential genes (29:36, 44.6%) (2.18 fold enrichment, p=0.0294 binomial, p=0.0055, 2D K-S Test based on co-expression data without median thresholding (*11, 12*)), showing that enhancer capture is a significant driver of new gene evolution alongside previously established processes (Figure 2).

### Generality of Enhancer-Capture Divergence

To further address the generality of the enhancer-capture model, we utilized an orthogonally defined, manually curated data set of newly evolved duplicate genes/parental gene pairs in *D. melanogaster* (N=156) which contained information regarding the duplication method of these genes (i.e., tandem, distal, or retro-transposition)(*13*). We also used a separate, publicly available FlyBase data set (*9*) containing 30 classes of developmental tissues produced by the modENCODE consortium (*14*) spanning 0-2 hour embryos to 30-day adults. Using these two data sources, we then calculated the Spearman correlation coefficient for the gene expression each new gene/parent gene pair across all 30 developmental conditions (“developmental co-expression”). Comparison of the developmental co-expression for tandem duplicates vs non-tandem duplicates (i.e., both distal duplicates and retro-transpositions) revealed significantly lower developmental co-expression in non-tandem duplicates than tandem duplicates (p=3.45 x 10^-10^, Supp. Figure S2). When these comparisons were done with tandem duplicates vs either distal duplicates or retro-transpositions alone, non-tandem duplicates continued to show significantly lower developmental co-expression (distal: p=8.99 x 10^-9^, retro-transposition: p=5.41 x 10^-3^, Supp. Figure S2). In support of the generality of the Enhancer Capture-Divergence model, these results demonstrate that regulatory neo-functionalization is a strong driver for non-tandem duplicates, as predicted by the asymmetric ECD model but not the symmetric DDC, EAC, or IAD models. Indeed, similar results have been observed in a wide range of studies, including but not limited to studies of retro-transposons and transposable element domestication as reviewed in (*15*) and (*16*).

### HP6/Umbrea as an Illustration of Enhancer Capture-Divergence

Evolution of the *HP6/Umbrea* locus is well-suited for demonstrating the enhancer capture-divergence model, as *HP6/Umbrea* is one of few recently-evolved genes in *D. melanogaster* whose protein evolution has been previously described in the literature (Figure 2b, denoted as (*)) (*17*). *HP1b*, a gene located on the X chromosome, duplicated approximately 12-15 million years ago (mya) into a gene-poor, intronic region of *dumpy*, located on chromosome 2L (Figure 3). The new gene, *HP6/Umbrea*, was the result of a full duplication, including its three known domains: the chromo domain, the chromo-shadow domain, and the hinge domain connecting the two.

**Figure 3.**
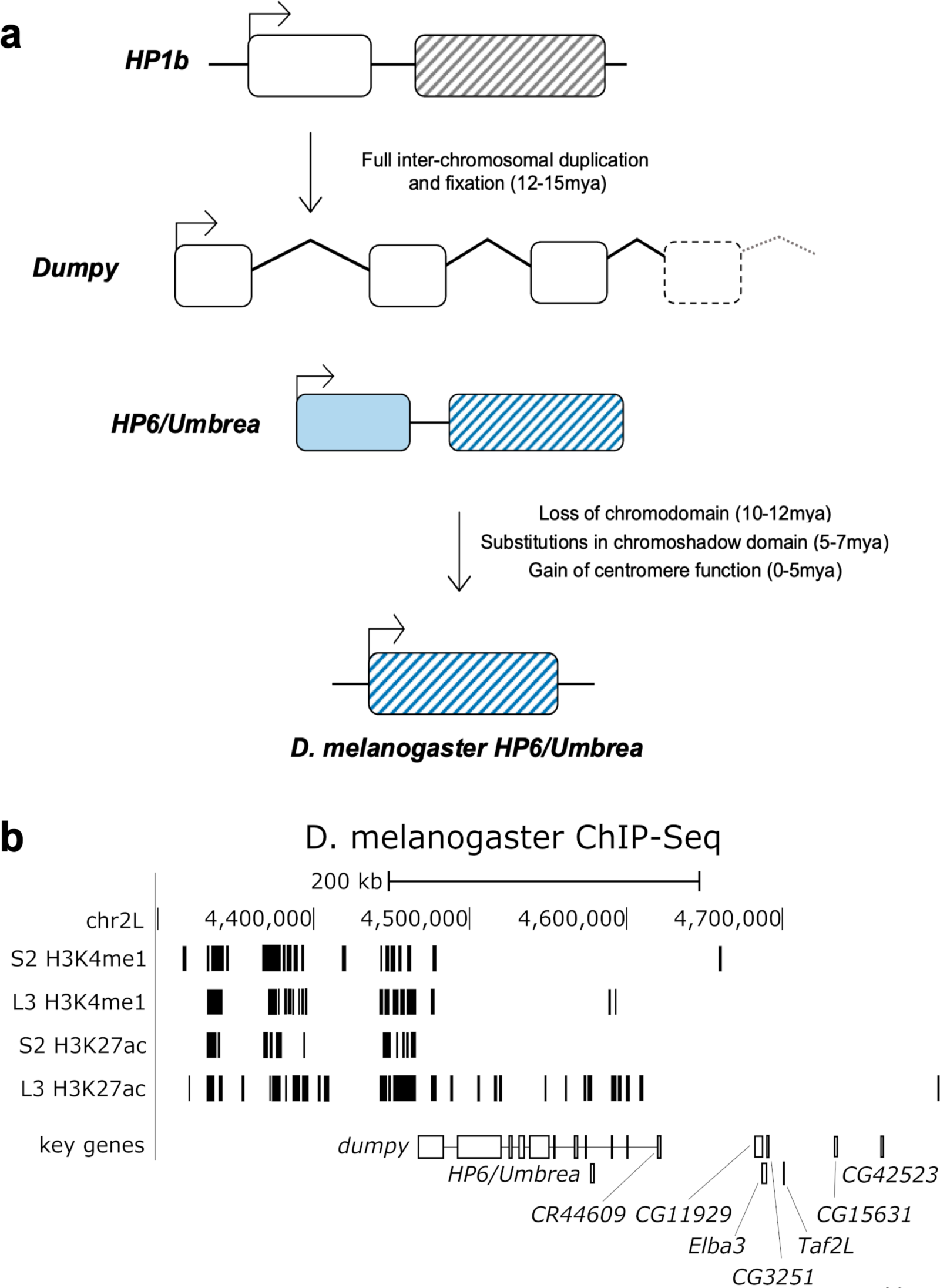
HP6/Umbrea likely evolved via enhancer capture. (a) *HP6/Umbrea* is a new gene in *D. melanogaster* that arose from a full duplication of *HP1b* into an intronic region of *dumpy*, migrating from chromosome X to 2L. *HP6/Umbrea*’s well characterized, step-wise protein evolution suggests that amino-acid substitutions were unlikely to have driven the duplicate gene copy to fixation. (b) A comparison of ChIP-Seq/ChIP-Chip markers for primed (H3K4me1) and active (H3K27ac) enhancers between embryonic S2 (no/low *HP6/Umbrea* expression) and whole L3 larvae (high *HP6/Umbrea* expression) tracks shows strong activation of a larval enhancer in a 100kb intronic region of *dumpy* that is, aside from *HP6/Umbrea*, devoid of protein coding genes.

Though *HP6/Umbrea* was lost ancestrally to multiple speciation events, suggesting that it was originally non-essential (*17*), *HP6/Umbrea* continued to evolve in a step-wise manner in the *melanogaster* lineage, gaining its semi-lethal phenotype (*18–20*) and rapidly diverging from its parental gene, *HP1b*. Subsequent to fixation, *HP6/Umbrea* lost its chromo domain approximately 10-12 mya; this was followed by sequence divergence and an accumulation of key substitutions 0-7 mya, resulting in *HP6/Umbrea’s* known centromeric protein function in *D. melanogaster* (*17, 18, 20–22*). The stepwise protein evolution from these results thus eliminates protein neo-functionalization as the driving force behind the fixation of *HP6*/*Umbrea*. Sub-functionalization and/or subsequent optimization of protein function may also be eliminated for similar reasons.

To determine whether enhancer capture is the primary driving force underlying *HP6/Umbrea*’s fixation, we examined the tissue expression of both *HP6/Umbrea* and *HP1b*. A simple comparison of *HP6/Umbrea*’s tissue-specific expression pattern to the parental gene *HP1b*’s very broad expression pattern suggests that *HP1b* is under constitutive regulation (Supp. Figure S1). Conversely, *HP6/Umbrea* is found only in a subset of tissues which express *HP1b*, suggesting that the new duplicate is under the control of one or more tissue-specific enhancers. *HP6/Umbrea*’s expression pattern is not similar to its first neighboring gene, *dumpy*, but its second neighboring gene, *CR44609*, which expresses in the imaginal discs, larval salivary glands, and male reproductive organs, which suggests that these genes are likely co-regulated. The non-complementary nature of the tissue expression patterns of *HP1b* and *HP6/Umbrea* provide further evidence ruling out sub-functionalization and/or subsequent optimization of regulatory function.

Consistent with the hypothesis that enhancer capture is the driving force being the evolution of *HP6/Umbrea*, publicly available modENCODE ChIP-Seq/ChIP-Chip data (*23*) provides positive evidence that enhancer capture likely drove its early evolution. Using the embryonic S2 cell line as a negative control where there is little to no *HP6/Umbrea* expression, primed (H3K4me1) and active (H3K27ac) enhancer marks in whole L3 larvae show strong enhancer activity in an intronic, gene-poor region of *dumpy*, coinciding with the onset of *HP6/Umbrea* transcription and its co-regulated, neighboring gene, *CR44609* (Figure 3b). Given the absence of other genes in the region (Figures 3b, 4a), *HP6/Umbrea* remains the likeliest target of the putative enhancer based on proximity and expression.

As *HP6/Umbrea* duplicated into a region that appears to be under the control of a pre-existing enhancer, we tested for further co-regulation in the region by using tissue expression data (c.f. the section of ***Analysis of Tissue Co-Expression Shows New Genes Evolve by Enhancer Capture***). We then applied a correlational analysis on this tissue expression data set to determine whether *HP6/Umbrea* is co-regulated with other neighboring genes. We took a 500kb region of the genome centered on the insertion site of *HP6/Umbrea* and calculated the tissue co-expression of each gene within this region in relation to *HP6/Umbrea*. As enhancers function in a proximity-based manner, we would expect a distance-dependent effect on the co-expression of neighboring genes across the genome. To generate a baseline estimate of this distance dependent co-expression distribution, we sampled 1000 random genic loci within the *D. melanogaster* genome, calculating the degree of co-regulation expected on proximity alone. Notably, we find that using this distribution, the region of influence of any given regulatory region of the genome appears to be on the order of 25kb, suggesting that this is a characteristic distance (1/e reduction) for enhancer interaction in *D. melanogaster* (Figure 4a). Outside of this region of influence, the likelihood of co-expression relaxes to the genomic average. Therefore, genes found within this region of influence with high tissue co-expression with neighboring genes are potentially the result of co-regulation with the focal gene. As expected, we find that the neighboring gene, *CR44609*, possesses a similar expression pattern as *HP6/Umbrea*. We also find that a locus of 6 neighboring genes (*CG11929, Elba3, CG3251, Taf12L, CG15631, CG42523*) located approximately 100kb away from *HP6/Umbrea* also expresses in the same tissues as *HP6/Umbrea*, expressing primarily in the imaginal discs, larval salivary glands, and adult male reproductive organs (Figure 4a).

**Figure 4.**
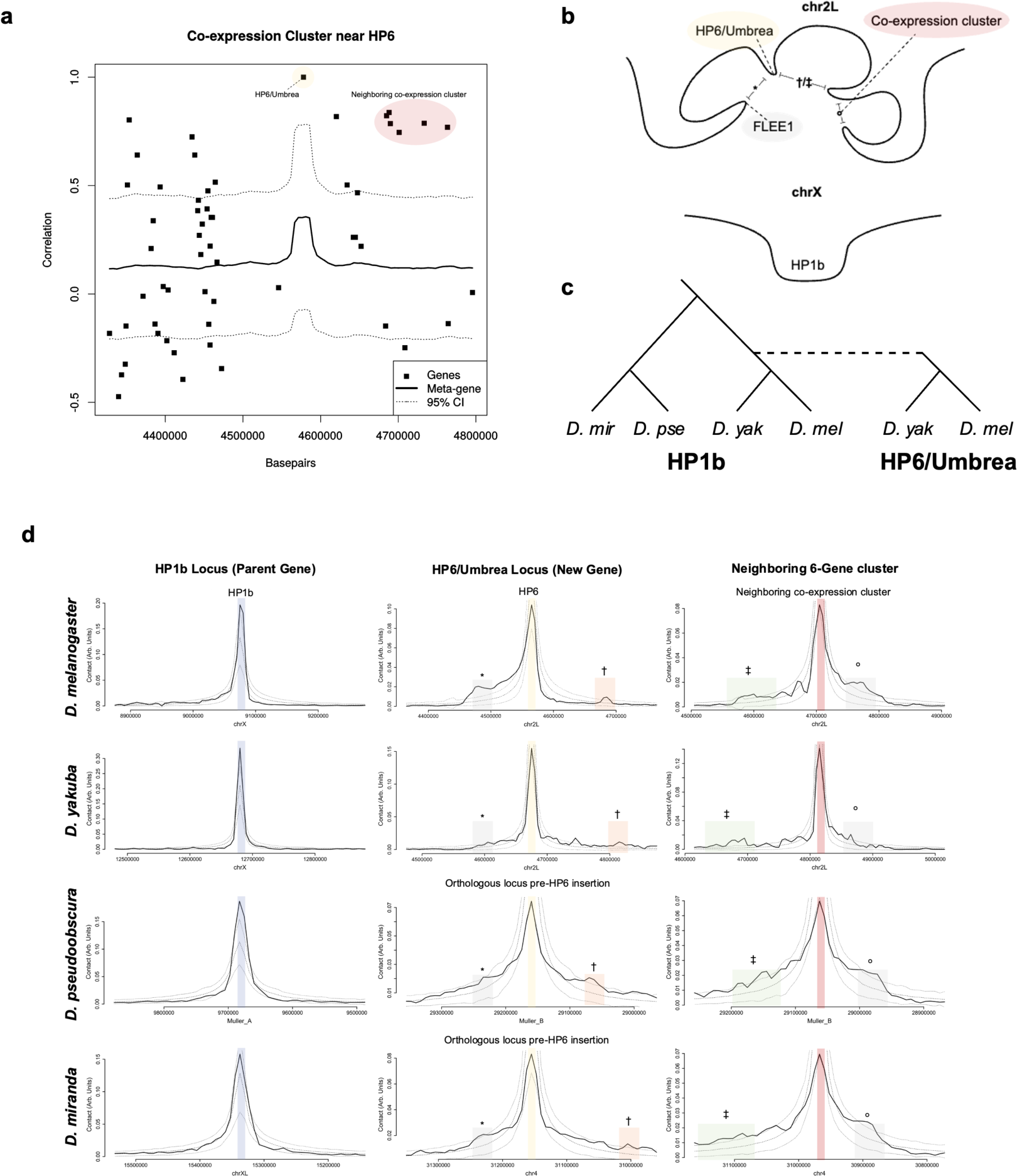
*HP6/Umbrea* co-expression is associated with conserved chromosomal looping that pre-dates its insertion. (a) Tissue co-expression analysis between *HP6/Umbrea* and neighboring genes reveals the presence of a co-regulated cluster of 6 neighboring genes. Note absence of other genes within *dumpy*’s intronic regions. (b) Two in-group species, *D. melanogaster* and *D. yakuba* (div. ∼ 6mya), contain *HP6/Umbrea*, while two out-group species, *D. pseudoobscura* and *D. miranda* (pse-mir div. ∼ 4mya, pse-mel div. ∼ 25mya), pre-date *HP6/Umbrea*’s insertion (∼ 12-15mya). (c) Cartoon legend illustrating features in (d). Not drawn to scale. (d) Hi-C data tracks for in-group (*D. mel, D. yak*) and out-group (*D. pse, D. mir*) species are shown for the parental gene *HP1b* (left column) *HP6/Umbrea*’s insertion site (middle column) and the co-regulated 6-gene cluster (right column), with a 95% confidence interval generated from genomic sampling plotted in dotted lines. On the vertical axis is contact in arbitrary units, and on the horizontal axis is genomic coordinates centered on the viewpoint location. Conserved feature (*) shows that *HP6/Umbrea*’s insertion site loops with the active larval enhancers contained in *dumpy*’s intronic gene-desert. Conserved features (†) & (‡) show that *HP6/Umbrea*’s insertion site reciprocally loops with the co-regulated 6-gene cluster. Conserved feature (^◦^) shows that the co-regulated gene cluster loops across the entire 6-gene cluster.

### HP6/Umbrea’s Chromosomal Conformations Are Determined by a Larval Enhancer Interaction

While the co-expression of *HP6/Umbrea*’s neighboring gene (*CR44609*) may be explained simply due to its proximity to *HP6/Umbrea*, the co-expression of the 6-gene cluster is not immediately evident as being a result of co-regulation. As the gene cluster of co-expressing genes is distally located along the chromosome beyond *HP6/Umbrea*’s 25kb region of influence, due to the 3-dimensional nature of the eukaryotic genome, these genes may in fact be proximally located near *HP6/Umbrea* in 3D space and thus be co-regulated. Similarly, while active enhancer marks correlating to the onset of expression appear ∼50-100kb away from *HP6/Umbrea*, it is not immediately clear that these putative enhancers are driving *HP6/Umbrea* expression, as their distance to *HP6/Umbrea* exceeds the 25kb region of influence. As the 3-dimensional conformations of the genome may still allow these distal genic elements to interact, we tested whether the putative larval enhancer, *HP6/Umbrea*, its neighboring gene (*CR44609*), and the 6-gene cluster are co-regulated by examining high-resolution Hi-C data for *D. melanogaster* (*24*) (Figure 4d).

Like co-expression, the frequency at which two genic elements make physical contact is expected to have a baseline, distance-dependent distribution. We may therefore test for co-regulation by predicting significant physical contact between *HP6/Umbrea*, its putative enhancer, and the cluster of co-expressed neighboring genes using Hi-C data in *D. melanogaster* (Supp. Figure S3). Such an interaction could be detected if contact between these two loci (i.e. *HP6/Umbrea* with enhancer and *HP6/Umbrea* with co-expressing genes) exceeds the baseline distance-dependent distribution of contact frequency. We generated an estimate of this baseline contact frequency distribution using 1000 independent loci that were sampled randomly from the genome, where contact data for the flanking regions were used to generate the baseline distance-dependent contact frequency distribution. We then extracted the contact frequency data for the *HP6/Umbrea* locus alone and compared this to the baseline genome-wide contact frequency distribution (Figure 4d).

We first note that after self-interactions are removed, we find that physical interactions in the genome generally remain highly localized, with most interactions lying near the focal locus as expected. Despite this, we find that *HP6/Umbrea*’s complex contact distribution shows significant contact with two key features: both with the putative larval enhancer as well as the neighboring 6-gene co-expression cluster (Figure 4d). Additionally, when this analysis is repeated for the 6-gene co-expression cluster, we find that this contact is reciprocated, as the 6-gene cluster shows significant contact across the cluster as well as with *HP6/Umbrea* (Figure 4d). Finally, *HP6/Umbrea* has enriched contact with the enhancer region that differentially activates at the onset of *HP6/Umbrea* expression. Combined with the tissue co-expression analysis, these results demonstrate that *HP6/Umbrea* and these 6 genes are likely co-regulated.

### The Complex 3D Genome Structure of the HP6/Umbrea Locus Is Conserved Over 25 Million Years, Pre-dating HP6/Umbrea Insertion

While we find evidence that *HP6/Umbrea*, the larval enhancer, and the 6-gene co-expression cluster are co-regulated, it is possible that these interactions evolved after *HP6/ Umbrea*’s insertion. To determine whether these interactions pre-date *HP6/Umbrea*’s insertion, we compared newly generated Hi-C data sets using a second in-group species, *D. yakuba* (Supp. Figure S4), and two out-group species, *D. pseudoobscura* and *D. miranda* (Supp. Figure S5, S6) (Figure 4c, d). While *HP6/Umbrea* inserted 12-15mya, the divergence between *D. melanogaster* and both outgroup species is 25mya (*25*). Within these clades, *D. melanogaster* and *D. yakuba* diverged 6 mya, while *D. pseudoobscura* and *D. miranda* diverged 4 mya. In comparing the Hi-C contact patterns for both *HP6/Umbrea* and its neighboring co-expression cluster, we find that key features of the local chromosomal conformation are conserved in the 3-dimensional structure despite 25 million years of evolution: contact with the larval enhancer, reciprocal contact between *HP6/Umbrea* and its co-expression cluster and contact across the entire co-expression cluster (Figure 4d). These features were found to be conserved in *D. miranda* even in the presence of a large-scale insertion between the future *HP6/Umbrea* locus and the neighboring co-expression cluster. The conservation of this chromosomal structure, despite the subsequent evolution of protein function of *HP6/Umbrea*, suggests that the neo-functionalization event driving the fixation of the original duplication was likely driven by enhancer capture. Specifically, the 3D structure driving enhancer contacts existed prior to *HP6/Umbrea*’s origination, and by duplicating into this region, *HP6/Umbrea* immediately captured this regulatory interaction.

### The HP6/Umbrea Locus Structure is Driven by a Tissue-Specific Larval Enhancer

Though the Hi-C data suggested that the captured larval enhancer would be located in the vicinity of chr2L:4500000, the interactions were observed at relatively low genomic resolution. To identify the location of the larval enhancer with higher precision, we utilized 4C-Seq (*26, 27*) on ∼400 dissected L3 larvae and pre-pupae from *D. melanogaster* using a viewpoint of the *HP6/Umbrea* coding sequence, revealing highly enriched contact with a single, distal 394bp locus located approximated 130kb away from the *HP6/Umbrea* locus (Supp. Figure S7). This locus was expanded by approximately 750bp on both 5’ and 3’ ends and was named the Four-C Larval Enhancer Element (FLEE1)(Supp. File S1). Interestingly, the 2165bp FLEE1 construct was found to be entirely contained within the coding regions of the genes *MFS18* and *Elp3*, which are both highly conserved, essential genes in *D. melanogaster*.

To validate whether FLEE1 contained a functional larval enhancer, we assayed pGreenRabbit reporter plasmids which we site-specifically integrated in *D. melanogaster* (BDSC 79604) (*28*). Compared to control, homozygote transformants that contain empty reporter vectors which drove basal levels of GFP expression, we found that FLEE1 directed GFP expression in the salivary glands of third instar larvae (Figure 5, Movie S1). The result is consistent with prior *in vivo* results of *HP6/Umbrea*’s known expression pattern and key role in polytene chromosome function in larval salivary glands. Anti-body staining of polytene chromosomes showed localization of *HP6/Umbrea* and its parental gene *HP1b*, while tissue-specific RNAi knockdown of *HP6/Umbrea* in the larval salivary glands demonstrated aberrant telomere-telomere attachments (*19*). Consistent with these observations, motif analysis of the FLEE1 locus using FIMO (*29*) and the CIS-BP database (*30*) revealed the presence of a *CrebA* motif, which is a leucine zipper transcription factor associated with regulation of tissue-specific genes in the salivary gland (*9*).

**Figure 5.**
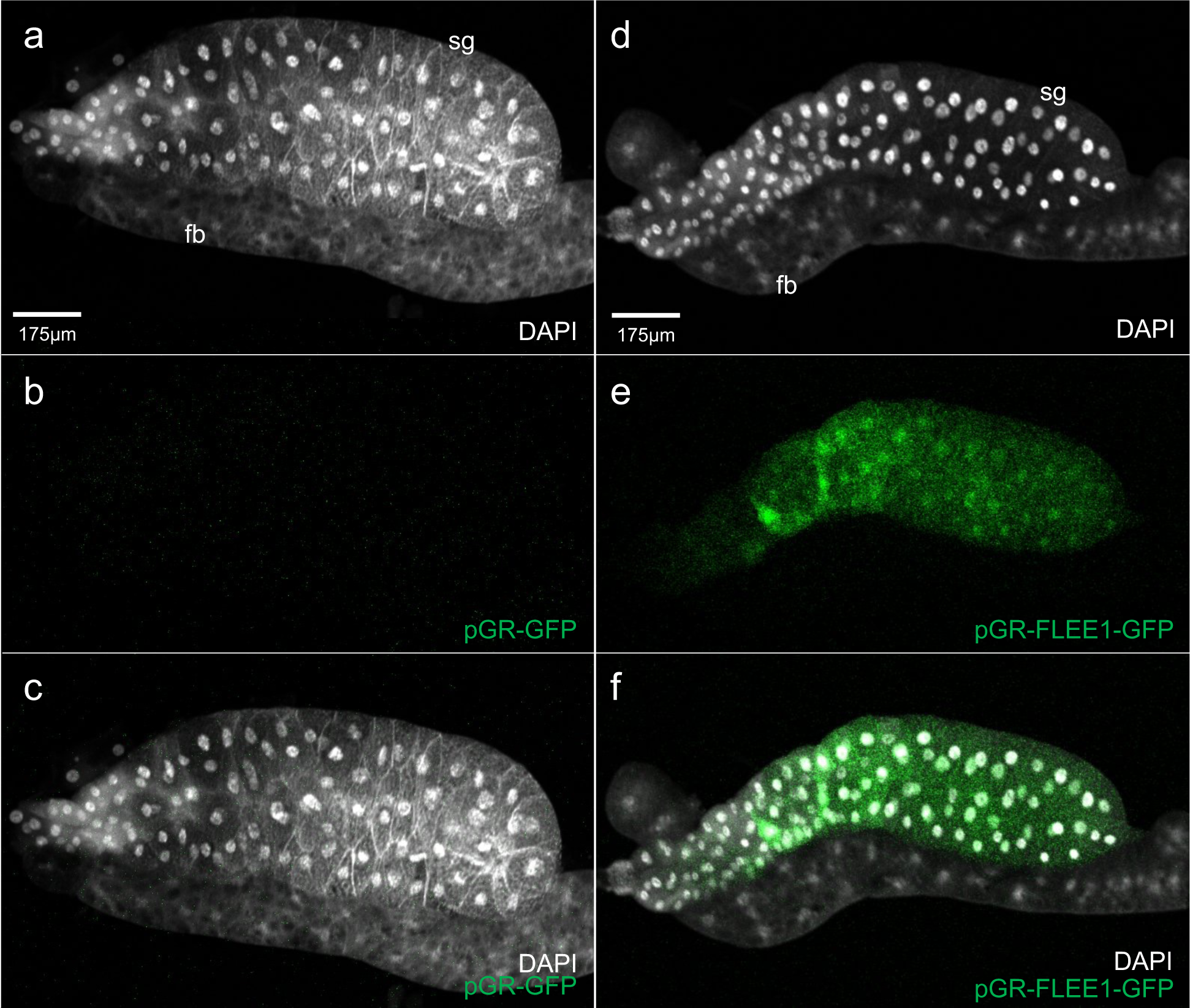
FLEE1 encodes a larval, salivary gland enhancer. Green, GFP. White, DAPI (DNA). sg = salivary gland. fb = fat body. The FLEE1 putative enhancer was tested for enhancer activity in third instar larvae using the pGreenRabbit reporter vector. (a-c) Basal GFP reporter expression from an empty reporter vector in a third instar salivary gland and fat body. (d-f) GFP reporter expression directed by FLEE1 in the salivary gland, with minimal expression in the fat body.

While FLEE1 contains the 3’ UTRs of both *MFS18* and *Elp3*, the entirety of FLEE1 is contained within the coding sequences of these two genes, excluding two short intronic regions of 54bp and 66bp within *MFS18* (Supp. Figure S8). Because *MFS18* and *Elp3* are essential genes, we were unable to perform further functional characterization. However, a population genetic analysis of the *MFS18* locus reveals that the coding sequence of *MFS18* is under selective pressure not only to maintain/conserve *MFS18* amino-acid sequence but also to maintain regulatory function as an active larval enhancer. Interestingly, the FLEE1 locus shows strong divergence from *D. yakuba* and *D. simulans* while maintaining low levels of polymorphism within natural populations in *D. melanogaster*, suggesting that the locus is under strong selective pressure (Supp. Figure S9). However, an analysis of the ratio of non-synonymous to synonymous substitution rates from *S. lebanonensis* to *D. melanogaster* for *MFS18* shows that the vast majority of these substitutions are synonymous substitutions (Ka/Ks = 0.033, p=0.0022)(*31*), demonstrating that this locus is under strong purifying selection. Alternatively, the *MFS18* locus fails to show signatures of directional selection, being unable to show significance in the correct direction under the Hudson-Kreitman-Aguadé (HKA)(*32*) and McDonald-Kreitman (MK)(*33*) tests (Table 1). These combined results suggest that the coding sequence of *MFS18* is under selective pressure to maintain/conserve *MFS18* amino-acid sequence while simultaneously maintaining regulatory function as an active larval enhancer, displaying a stereotypically high substitution rate as is common with enhancers under stabilizing selection(*34*). These results stand in sharp contrast to the *HP6/Umbrea* locus, which shows signatures of strong directional selection under both the HKA and MK tests (Table 1).

**Table 1.**
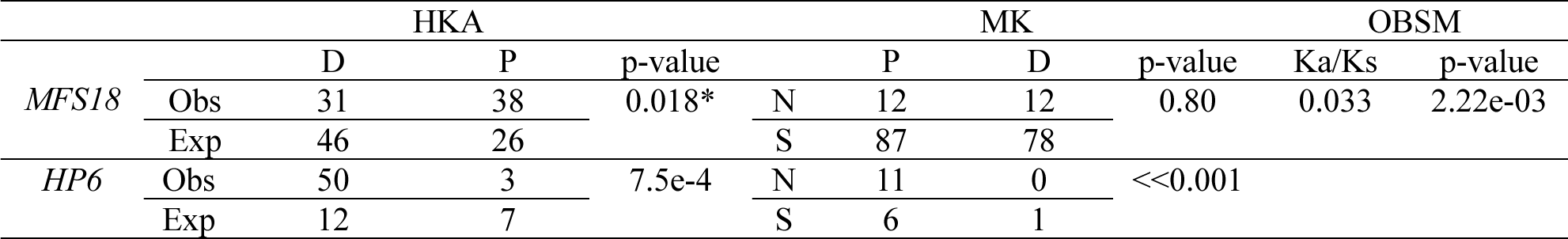
Population genetic analysis of enhancer and HP6/Umbrea locus. The test summaries of HKA, MK, and OBSM for positive selection of *MFS18* and *HP6/Umbrea*. “D, P, N, S, NS, Obs, and Exp” indicate divergence, polymorphism, nonsynonymous sites, synonymous sites, non-significant (p>0.05), observed numbers and proportions, and expected proportions, respectively. *indicates divergence is significantly *lower* than expected, one-sided Fisher’s Exact

### Co-expression of HP6/Umbrea and Neighboring Genes Pre-dates HP6/Umbrea Insertion

The distal 3D-dimensional contacts (spanning over 250kb from enhancer to co-expression cluster) we identified were found to pre-date *HP6/Umbrea*’s insertion, and these contacts were used to discover a previously uncharacterized enhancer element. However, it is still unclear whether the co-regulation of the co-expressed cluster pre-dates *HP6/Umbrea*’s insertion. The positioning of FLEE1 within a highly conserved, essential gene prevents *in situ* genetic manipulation to validate an ancestral co-regulatory environment. Therefore, to determine whether co-regulation of *HP6/Umbrea*’s insertion site and its neighboring genes pre-dates the insertion of *HP6/Umbrea*, we performed single-cell RNA-Sequencing (scRNA-Seq) using a panel of 3 closely related species: *D. melanogaster* and *D. yakuba*, both containing *HP6/Umbrea*, and *D. ananassae*, which pre-dates *HP6/Umbrea*’s origination. We performed scRNA-seq in the testis tissue because of its high evolutionary importance (*20, 35–37*), the existence of pre-existing high-quality cell type annotations (*38*), and the significantly higher expression levels of *HP6/Umbrea* and its co-expression cluster in this tissue type relative(*38*), and the significantly higher expression levels of HP6/Umbrea and its co-expression cluster compared to imaginal disc or salivary gland tissue (Supp. Figure S1).

After mapping and visualization of the scRNA-Seq data using previous cell type annotations (Figure 6a)(*38*) as well as data from all three species on the same, shared manifold (Figure 6b), it becomes clear that *HP6/Umbrea* is co-regulated on a cellular level with *CG11929, Taf12L* and *CG15631*, while overall expression of *Elba3* and *CG3251* are low and restricted mainly to germline stem cells (GSCs)/early spermatagonia. As an internal control, somatic and developmental cell types cluster together as expected. The bulk of the expression is shared across the co-regulated genes, while further cell type-specific expression is also shared within a sub-class of cyst cells, GSCs/early spermatogonia, and early as well as late spermatids. Importantly, the co-regulation of *CG11929, Taf12L*, and *CG15631* pre-dates the insertion of *HP6/Umbrea* as demonstrated by the shared cell type-specific co-expression of these genes in *D. ananassae*. While co-regulation of *CG11929, Taf12L*, and *CG15631* and very low expression of *Elba3* and *CG3251* remain evolutionarily conserved, *CG42523* shows significant divergence in its expression pattern within these species. Notably, while its expression pattern shows significant co-regulation with *CG11929, Taf12L*, and *CG15631* in *D. ananassae*, it appears that *CG42523* was down-regulated in the *D. yakuba* lineage, while it shows significant functional divergence in its regulation from the co-expression cluster in the *D. melanogaster* lineage.

**Figure 6.**
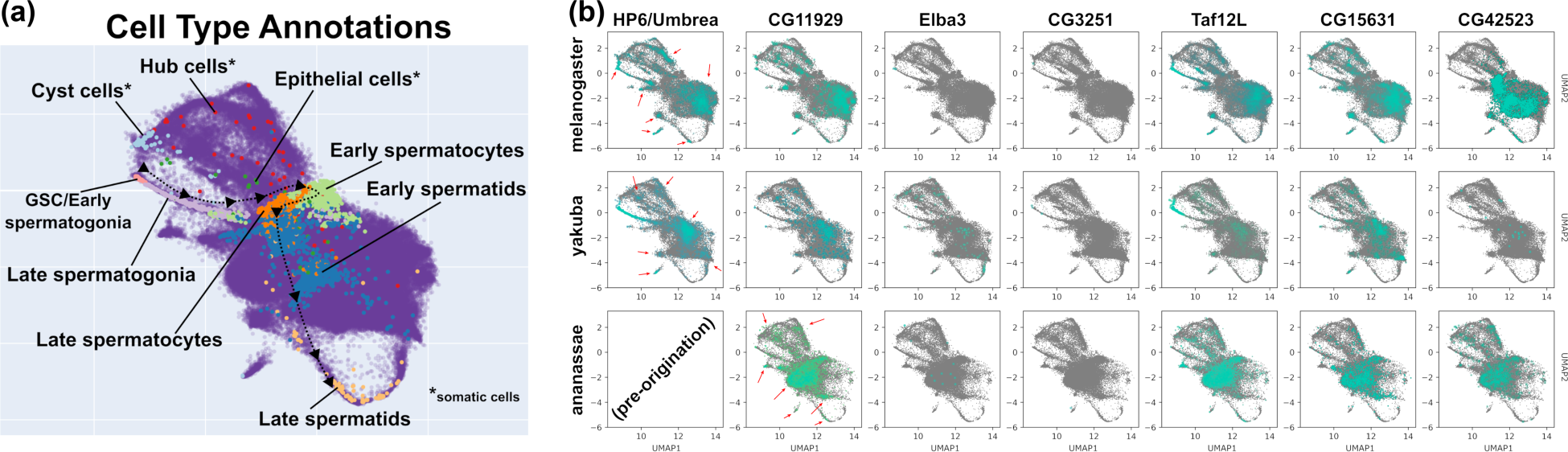
Regulation of *HP6/Umbrea* locus is ancestral. scRNA-Seq data from *D. melanogaster, D. yakuba* (both containing *HP6/Umbrea*) and *D. ananassae* (pre-dating *HP6/Umbrea* origination) are mapped to the same manifold and visualized. **(a)** Pre-existing data from and their corresponding labels from prior studies were included, allowing for precise identification of somatic (cyst, hub, and epithelial cells) and developmental (germline stem cell (GSC)/early spermatogonia, late spermatogonia, early and late spermatocyte, and early and late spermatid) cell type clusters both within and across species. The developmental trajectory of spermatogenesis is indicated using lines and arrows. **(b)** Expression data from *HP6/Umbrea* and the co-expression cluster of genes are plotted for all three species. While the bulk of expression in early spermatids is generally conserved within and across species for *HP6/Umbrea* (when present), *CG11929, Taf12L*, and *CG15631*, finer cell type-specific expression patterns (red arrows) are also conserved for these genes within each species, indicating conservation of co-regulation pre-dating the insertion of *HP6/Umbrea*. Furthermore, low levels of GSC/early spermatogonia-restricted expression of *Elba3* and *CG3251* are also conserved across species. Interestingly, *CG42523* shares the same cell type-specific expression patterns as *CG11929, Elba3*, and *CG15631* in *D. ananassae*, but diverges in expression both in *D. yakuba* and *D. melanogaster*.

## DISCUSSION

### Identification of Distal Larval Enhancer

FLEE1’s regulatory activity residing within the primarily exonic regions of the highly-conserved *MFS18* gene constitutes an example of how protein-coding regions of the genome may also have key regulatory functions (*39, 40*). Such pleiotropy demonstrates how the interpretation of synonymous substitution rates may not necessarily serve as good estimates of neutral evolution rates in commonly used codon table-based tests of molecular evolution. Rather, substitutions typically regarded as synonymous could alternatively be indicative of strong directional or stabilizing selection for the regulatory function of genomic enhancer elements. Furthermore, elucidation of the FLEE1-*HP6/Umbrea* interaction highlights the importance of identifying and characterizing the contributions of structural variations and chromosomal rearrangements in driving phenotypic evolution. Our results demonstrate how stabilizing selection for the conservation of large-scale chromosomal conformations drives the appearance of evolutionary novelty resulting in the development of novel, centromeric function as in the case of *HP6/Umbrea*. While further work will be required to reveal what evolutionary forces underlie the strong conservation of the long-distance interaction of the *HP6/Umbrea* locus prior to the new gene’s insertion, our findings demonstrate how complex chromosomal conformations are a key, underappreciated element in the evolution of the eukaryotic genome.

### Enhancer Capture Divergence Model

ECD joins various previously proposed models to interpret different evolutionary aspects of gene functionality. Whereas DDC and EAC, for example, explain the duplication-dependent subfunctionalization and resolution of adaptive-caused conflict from ancestral genes with multiple functions respectively, ECD interprets neofunctionalization for creating novel gene functions through duplication. ECD demonstrates how the manner of duplication itself may provide neo-functionalization in an asymmetric, tissue-specific manner. Such neo-functionalization provides a selective advantage in a direct, by the single-step 3D-facilitated acquirement of regulatory elements. The newly acquired elementary functions may maintain the new duplicate for adequate time until the new advantageous mutations occur to solve the Ohno’s dilemma.

In addition to partial duplication phenomena such as the generation of gene fusions (*41*) as well as favorable frame-shifts (*42*), our model highlights the under-appreciated evolutionary value of both the act of duplication itself and, more importantly, the genomic context in which these duplications occur. While the role of positional effects in gene regulation and evolution has long been appreciated (*43, 44*), the advent of new chromosomal conformation capture technologies allows us to directly connect the conservation of chromosomal domains (*45, 46*) and the origination of new genes under a strong conceptual framework.

Under the ECD model, a gene copy duplicates into a pre-existing regulatory context (Figure 7a), gaining a new regulatory interaction. This model thus provides a mechanistic explanation by which gene interaction networks may rapidly evolve (*36*). Under this model, we have two separate gene interaction sub-networks for both parental and neighboring genes (Figure 7b). As a new gene duplicates into a region near the neighboring gene, the new gene acquires the upstream regulatory function of the neighboring gene as well as the original parental gene’s downstream protein function (Figure 7c) while simultaneously preserving the pre-existing interactions from both parental and neighboring genes’ sub-networks. Since duplication has been observed to occur more frequently than point mutations (*4, 7*), enhancer capture provides a faster route to generating increased tissue-specific expression of a parental gene (Figure 1) than any set of mutations in the parental gene’s regulatory sequence. Duplication in the 3-dimensional looping architecture of the eukaryotic genome recombines genes and enhancers into new combinations, thus resulting in regulatory novelty (Figure 7c). As such, this model provides an explanation and mechanism for the well-described but poorly-understood phenomenon where new gene duplicates often possess highly tissue-specific expression patterns (Figure 2a)(*37, 47, 48*).

**Figure 7.**
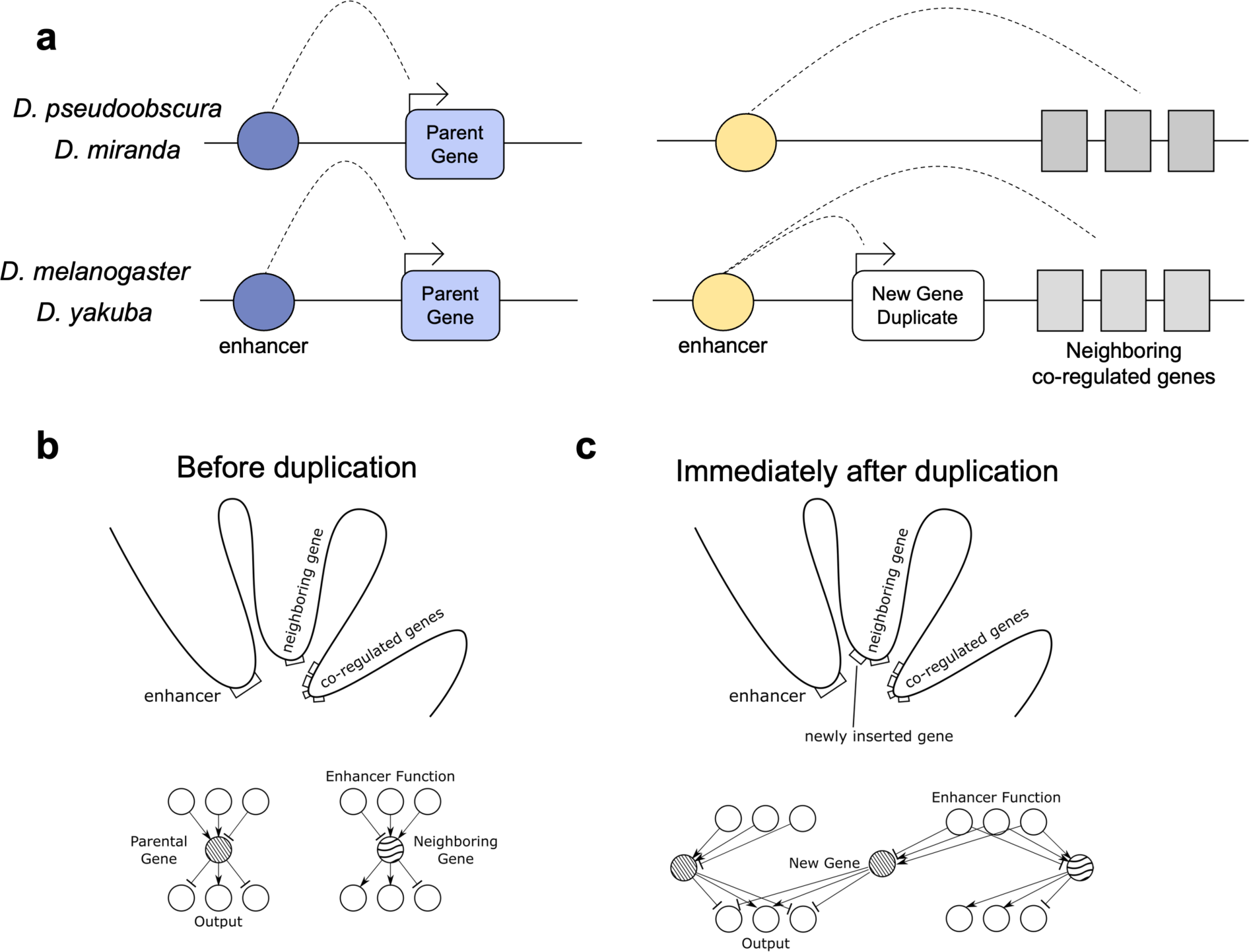
The 3D organization of the genome allows for rapid rearrangement of genetic networks. Panel (a) depicts a cartoon illustration of the action of the larval enhancer on the neighboring cluster of co-regulated genes as well as the future insertion site of *HP6/Umbrea*. Preceding insertion of *HP6/Umbrea*, the larval enhancer was in contact with both *HP6/Umbrea*’s neighboring gene as well as with the co-regulated 6-gene cluster. (b, c) This looping structure remains conserved following *HP6/Umbrea*’s insertion, allowing for a rapid recombination of elements upstream of *HP6/Umbrea*’s neighboring gene (i.e., larval enhancer) with elements downstream of *HP6/Umbrea*’s parental gene (i.e., *HP1b*’s protein function). A sample gene interaction network, both pre-& post-duplication, is also depicted above. Note that the parental gene and neighboring gene’s original interactions remain intact, preserving previous function.

One key aspect of the ECD model is the selective advantage imparted by increased tissue-specific expression. The resolution of genetic conflict, such as sexual antagonism, is becoming increasingly appreciated as a driver of the evolution of new genes (*49, 50*). While most new genes have highly tissue-specific expression patterns, these often favor either the female or male reproductive organs/germlines in *D. melanogaster* (*37*). A close examination of the expression pattern of *HP6/Umbrea* demonstrates the same – *HP6/Umbrea* is expressed primarily in the imaginal discs, larval salivary gland, and the male reproductive organs. As such, it is possible that the selective advantage imparted by *HP6/Umbrea*’s original duplication may have been a result of regulatory sexual antagonism and, given that most new genes show expression specific to reproductive organs, enhancer capture may be a widespread mechanism for the resolution of sexual antagonism. Furthermore, *HP6/Umbrea*’s repeated ancestral loss suggested it was originally non-essential follow duplication, but later gained its semi-lethal phenotype in a step-wise manner. The enhancer capture model may provide a key mechanism by which new essential genes gain their essential phenotype(*20*).

### Evidence for the generality of the ECD mechanism

One context in which the mechanism of enhancer capture has been well studied is in the development of human malignancies. The first discovered example of enhancer capture occurring in cancer is the t(8;14) translocation in Burkitt’s lymphoma, allowing for the oncogene *Myc* to be expressed under the regulatory control of the immunoglobulin heavy chain gene (*IGH*), which is expressed in lymphoid cells (*51, 52*). To date, there have been a variety of other examples of enhancer capture re-arrangements involved in oncogenesis occurring in diverse tissues (*53–57*). Although most re-arrangements bring oncogenes into proximity with constitutive regulatory elements of a given cell type, they may also be brought into proximity with context-specific regulatory regions. One such translocation in prostate cancer involves the translocation of the oncogenes *ETV1* or *ERG* within proximity of the promoter region of *TMPRSS2* which contains several androgen receptor binding sites. In this instance, *ETV1* or *ERG* gains new androgen-dependent expression, which can be abrogated by androgen deprivation therapy, a common treatment for prostate cancer(*58, 59*).

One longstanding question in this literature is why particular re-arrangements are commonly associated with specific cancers. Are common re-arrangements observed because they are the few examples that confer a selective advantage in select cell or does the cancer cell type have a structural predisposition to favor those re-arrangements? Although common rearrangements do confer a relative fitness advantage to the cancer cells, the evidence has become clear that re-combining loci are likely to be within physical proximity of one another(*60, 61*). It has been shown that chromosomes 9 and 22 neighbor each other in hematopoetic cells which may explain the frequency of the t(9:22) translocation in chronic lymphocytic leukemia which produces the *BCR-ABL* fusion protein(*62, 63*). Additionally, it has been shown that the *Myc* and *IGH* genes are brought within close physical proximity during B-cell stimulation (*64*) highlighting the importance of cell-context specific genomic arrangements in cancer.

The primary difference between enhancer capture in cancer and organismal evolution is the lack of necessity for cancer cells to preserve an oncogene’s previous function via gene duplication prior to translocation. Additionally, cancer cells typically experience selection at the clonal level so re-arrangements do not need to confer optimized gene expression within multiple tissue contexts. However, the cancer literature is clear that enhancer capture is a commonly occurring one-step mechanism that allows individual cells to gain fitness advantages, and that cell-type specific 3D genome confirmations selectively favor certain re-arrangements. Given our finding that the enhancer capture-divergence model is a significant driver of new gene evolution, it is likely that the inherent 3D configuration of the germline genome imposes a significant and previously unappreciated constraint on evolutionary novelty.

## METHODS AND MATERIALS

### Tissue expression data and analysis

Tissue expression data were retrieved from FlyBase. Pre-computed RPKM data files were downloaded, with RPKM values for each FlyBase transcript being reported for 29 tissues (*8*). As many of the tissue-types were repetitive, data from the head, ovary, carcass, and digestive system were averaged to reduce over-representation bias in further correlational analyses. Gene map data was also obtained from FlyBase to properly identify neighboring genes (*9*). Parental/new gene pair information was retrieved from (*20*). Spearman correlation coefficients were calculated using the tissue expression data between parental and new gene pairs. Due to intronic structures and variation in gene length, two neighboring genes for each new gene on each side were assessed using Spearman correlation coefficients and the maximum value of the four neighbors was recorded. Additionally, correlation coefficients for all genes within 500kb of *HP6/Umbrea* were reported. To generate a baseline distance-dependent genomic estimate of co-expression, 1000 random genic loci were chosen and co-expression values (Spearman) between the randomly selected gene and all neighbors within a 500kb range were calculated. This 500kb region was then divided into 100 non-overlapping windows where means and variances in correlation coefficients were calculated across all randomly selected loci.

### ChIP-Seq data

ChIP-Seq or ChIP-Chip data were obtained for H3K4me1 and H3K27ac for S2 cells as well as whole L3 larvae from modENCODE (*23*). H3K4me1 ChIP-Chip data for S2-DRSC cells were obtained using data ID 304 and 3760. H3K27ac ChIP-Chip data for S2-DRSC cells were obtained using data ID 296 and 3757. H3K4me1 ChIP-Seq data for whole Oregon-R L3 larvae were obtained using data ID 4986. H3K27ac ChIP-Seq data for whole Oregon-R L3 larvae were obtained using data ID 5084. For all data sets, data was obtained in .gff3 format and visualized using the UCSC Genome Browser.

### Hi-C data

We generated Hi-C data of *D. yakuba, D. pseudoobscura,* and *D. miranda*, and used publicly available Hi-C libraries obtained from NCBI for *D. melanogaster*, PRJNA393992. *D. melanogaster* source tissue was S2 cells, *D. yakuba* from adult females (SRR12331759), and *D. pseudoobscura* and *D. miranda* were L3 larvae. Hi-C libraries were preprocessed, mapped, and filtered using HiCUP version 0.8.0 (*65*). Specifically, reads from fastq files were trimmed at ligation junctions, and subsequently each mate of paired-end sequences were independently mapped to the respective genomes using bowtie2 version 2.2.9 (*66*). Reads were mapped to genomes consisting of canonical chromosomes only (i.e., excluding scaffolds and other unplaced sequences). *D. melanogaster* reference genome was dm6 and obtained from FlyBase (*9*). The *D. yakuba* reference genome for the NY73PB line was generated by meta-assembly of two PacBio long read assemblies (FALCON and Canu) using quickmerge, followed by polishing with Quiver, PILON, and a custom FreeBayes homopolymer frameshift polishing step. It can be obtained from NCBI (PRJNA310215). The *D. pseudoobscura* reference genome was obtained directly from Ryan Bracewell (https://www.ryanbracewell.com/data.html) (*67*) and the *D. miranda* reference genome was obtained from NCBI (PRJNA474939), (*68*). HiCUP was used further to remove experimental artifacts based on an *in silico* genome digest as previously described (*65*). HiCUP mapped and filtered .sam files were then converted to formats compatible with HOMER version 4.11 (*69*) and juicer tools version 1.22.01 (*70*). To create matrices, HOMER was used to tile the genome into matrices of fixed-size bins, and assign reads to their correct intersecting bins. HOMER was also used to normalize contact counts in these matrices based on known Hi-C biases, as previously described (*69*). JuicerTools was used to produce .hic files at resolutions of 5kb for *D. melanogaster* and *D. yakuba* and 7.5kb for *D. pseudoobscura* and *D. miranda*, and to create normalized matrices.

Using Hi-C contact matrices, data rows for *HP6/Umbrea* and its neighboring cluster were pulled for a 400kb region centered on *HP6/Umbrea* and self-self interactions were removed. To generate a genome-wide distance-dependent distribution of contact, 1000 random loci were sampled. Contact data for each locus was then normalized with total contact (arb. units) being equal for all loci. The means and variances for each non-overlapping window were calculated and reported and compared to *HP6/Umbrea* and the co-expression clusters’ data. To generate genomic coordinates for *HP6/Umbrea* prior to duplication, *D. melanogaster* sequences flanking *HP6/Umbrea*’s insertion site were aligned to the *D. yakuba, D. pseudoobscura* and *D. miranda* reference genomes using BLAST. Similarly, the promoter region of CG11929 was aligned to *D. yakuba, D. pseudoobscura* and *D. miranda* reference genomes to represent the co-expression cluster.

### 4C-Seq Data

About 400 *D. melanogaster* L3 larvae and pre-pupae were freshly dissected in 10-minute intervals on ice. A single cell suspension was generated from imaginal disc tissue using collagenase. These suspensions were pooled and formaldehyde-fixed for 10 minutes, followed by glycine quenching. Aliquots of these suspensions were quantified and snap frozen with liquid nitrogen and stored at -80°C until 10^7^ cells were accumulated. All cells were then collected and resuspended in a lysis buffer containing Triton X-100, NP-40, and protease inhibitors followed by homogenization via douncing. Nuclei were then gently lysed using a SDS and Triton-X while shaking (900 RPM) at 37°C for 1 hour each. Restriction enzyme digests were then performed using DpnII. After enzymatic deactivation at 65°C, the resulting solution was diluted in 7 mL of water, and proximity ligation was performed using T4 ligase overnight. This was followed by overnight de-crosslinking using proteinase K. A second restriction enzyme digest was performed with Csp6i followed by a second proximity ligation step performed in 14 mL solution. The resulting circularized library was extracted with ethanol and then purified using a HiPure PCR Cleanup kit. The cleaned library was then amplified using primers specific to HP6/Umbrea with attached Illumina P5/P7 adapters and sequenced on the Illumina HiSeq 2500 platform (PRJNA948431). Results were subsequently aligned to the FlyBase dm6 reference genome, and raw coverage was visualized in R using rtracklayer.

### Fly stocks, genetic manipulations and microscopy

All *D. melanogaster* lines were grown on a modified Bloomington cornmeal-molasses formulation. Fly lines for site-specific integration were obtained from Bloomington Drosophila Stock Center. pGreenRabbit reporter plasmids were site-specifically integrated into y[1] w[*] P{y[+t7.7]=nanos-phiC31\int.NLS}X; P{y[+t7.7]=CaryP}attP40 (BDSC 79604). FLEE1 (2L:4444468-4450632) was amplified by PCR and cloned into the pGreenRabbit vector, following traditional cloning methods. We injected an empty pGreenRabbit vector as a negative control and pGreenRabbit with the FLEE1 insert into BDSC 79604 pre-blastoderm embryos. Flies with successful integration were screened for the red eyes phenotype (presence of mini-white). We dissected salivary glands from third instar larvae of homozygous transformants in 1X PBS, fixed in 5% PFA in 1XPBS for 5 minutes, and washed 4 times in 1X PBS for 5 minutes. Fixed salivary glands were stained with DAPI (1:1000) for 10 minutes. All imaging was carried out on an upright laser scanning confocal microscope (Zeiss LSM 710) and similarly processed using ImageJ software.

### Population Genetic Analysis

*The data analysis.* The genomic variants were called from whole genome sequencing of 25 samples of *D. melanogaster* (DRM36, EA87, EA87N, ED10N, EF10N, EF126N, GA01, GA03, GA06, GA07, GH01, GH06, GH12, GH16, GH17, MC23, MC28, RAL900, RG18N, RG4N, UM118, UM37, UM526, ZH16, ZH20), ten samples of *D. simulans* (F11R4, F11R5, F21R2, F21R3, F31R2, F31R3, F31R4, F31R5, F41R1, F41R2), and five samples of D. yakuba (CY02B5, CY08A, CY13A, CY17C, CY22B), with sequencing depths >10(*71*). All these publicly available raw reads were downloaded from NCBI and cleaned with fastp(*72*). The cleaned reads were then mapped to the reference genome of BDGP6.32 with bwa mem v0.7.12(*73*). The variants-calling steps included marking duplicates, recalibrating base quality scores, per-sample calling with HaplotypeCaller, joint-calling with GenotypeGVCFs, and SNPs annotation with snpEff(*71, 74*). Only the biallelic sites with quality score > 30, minimum coverage of 10X, minimum genotype quality of 30, a maximum of 25% missing data were kept.

HKA-like tests (*32, 75*) and MK tests (*33*) were conducted using polarized SNPs by focusing on fixed homologous sites in all outgroup samples (*D. yakuba* and *D. simulans*). The allele frequencies for D. melanogaster and outgroups were estimated with PLINK v1.9(*76*). The expected proportions of diverged and polymorphic sites were calculated using the entirety of chromosome 2L (547951/ 307551=1.78). The proportions of diverged and polymorphic sites for genes were compared against the chromosome-wide ones with χ2 test (d.o.f.=1).

To detect signals of natural selection based on Ka/Ks (also ω) at the loci of *MFS18*, we collected orthologous sequences of these two genes in 10 Drosophilid species (*D. ananassae, D. erecta, D. melanogaster, D. mojavensis, D. pseudoobscura, D. simulans, D. virilis, D. willistoni, D. yakuba, Scaptodrosophila lebanonensis*) from OrthoDB v11(*77*). For *HP6/Umbrea*, Ka/Ks ratio was not computed due to incomplete ORFs in outgroup species. We used a codon-based alignment computed with TranslatorX and MAFFT(*78, 79*) for *MFS18* to generate gene trees and conducted the branch model test implemented by PAML(*80*). To determine the optimal branch model for substitution rate estimation, we used a dynamic programming method by Zhang et. al. (*31*) to select the optimal model according to log likelihoods.

*The sojourn time of a neutral polymorphic duplicate before loss in a population:* The question to address is how long a newly formed duplicate, if slightly deleterious (as was previously shown for various polymorphic duplicates (*1*)), can stay in a form of polymorphism in a population before loss due to genetic drift. The fixation probabilities for various polymorphic duplicates were calculated using the equation: u/u_o_ = S/(1-e^-S^) where u_o_ = 1/2Ne as the fixation probability of a neutral mutation, S = 4N_e_s and s the selection coefficient (*81*). The selection component, *γ* = 2N_e_s, for various polymorphic duplicates in *D. melanogaster* were experimentally measured(*1*). The average sojourn time before a neutral duplicate mutation disappears from a population was calculated as T_0_(1/2N) = 2(Ne/N)In(2N) where Ne is effective population size and N actual population size (*82*). The average ratio Ne/N was reported in *D. melanogaster* as 0.027 (*83*) and a general estimate for metazoans as 0.10(*84*). The T_0_(1/2N) < 1.04 ∼ 3.60 generations (the median as 2.32 generations) (Ne = 3,300,000 in *D. melanogaster*(*2*)), because all the duplicate variants are slightly deleterious (*1*) and could disappear even sooner. Furthermore, the point mutation rate, as reported previously (e.g.(*2, 3*), is in the orders of 10^-8^ ∼ 10^-9^ per site per generation and the advantageous ones even much more rare, is unlikely to generate any genetic change that can rescue the duplicate from extinction in so short a time.

### scRNA-Seq

Testes from *D. melanogaster* (*38*), *D. yakuba* (newly generated), and *D. ananassae* (newly generated) were dissected in drops of cold PBS using forceps on Petri dishes before being transferred on ice to reduce degradation. We then desheathed testes in lysis buffer (196 µL 1X TrypLE + 4 µl 100mg/ml collagenase). After spinning down briefly and incubating at room temperature for 30 minutes with mild vortexing every 10 minutes, the samples were passed through 35 µm filters before centrifuging for 7 minutes at 163g (1200rpm) at 4°C. We removed the supernatant, washed the cell pellet with 200 µL cold HBSS, and centrifuged again for 7 minutes at 163g (1200rpm) at 4°C. We then removed the supernatant before resuspending the cell pellet in 35 µL cold HBSS. We counted cells and checked viability on an automated cell counter using 5 µL of the single cell suspension with 5 µL of trypan blue. Samples were then sent to Rockefeller Genomics Center for 10X single-cell library preparation and sequencing.

The resulting libraries were processed using cellranger (v7.1.0) and aligned to RefSeq genomes obtained from NCBI (*mel*: GCF_000001215.4, *yak*: GCF_016746365.2, *ana*: GCF_017639315.1). Pair-wise alignments were performed for each species’ transcriptomes using BLAST and subsequently imported into SAMAP (v0.3.0) (*85*). Additionally, raw, unprocessed count data from cellranger was converted into Seurat (v4.3.0) objects in R and then exported as .h5ad AnnData files. These unprocessed AnnData files were then imported into SAMAP, where expression data for all three species were mapped to the same manifold and subsequently visualized. Cell type labels from (*38*) were imported into SAMAP and visualized, allowing for cell type classification. As overall expression levels of the genes within the visualized cluster varied, resulting in certain genes’ visualizations being saturated and therefore uninterpretable, the scale factor for these genes was manually adjusted in SAMAP to allow for a consistent interpretation across genes.

### RNAi and lethality measurements

We used lethality data previously published by our lab (*20, 21*) that was based on RNAi lines obtained from the Vienna Drosophila Resource Center (VDRC). A quarter of all KK RNAi lines from VDRC carry an inverted repeat sequence insertion at 30B3. However, a proportion (23–25%) of KK lines also carry an insertion at 40D3, which is housed within the *tio* locus and produces a confounding lethal phenotype. To avoid this, we updated the lethality data of new genes reported in (*20*) by removing the *tio* insertion site in KK lines using a recombination-based approach (*21, 86*) and finally derived lethality data for the new genes. The lethality results for all lines without insertion in the *tio* locus were reproducible, previously having been analyzed using four replicates, and again in our analysis in duplicate. Distally duplicated genes had 90% fewer offspring relative to control flies after Act5c-GAL4 induction were labeled as essential.

## Acknowledgements

We thank Dr. Marc Halfon and Dr. Sarah Bray for sharing the pGreenRabbit plasmid. We thank Hong Duan and Connie Zhao at the Genomics Resource Center of Rockefeller University for their help with the scRNA-seq libraries. This manuscript is dedicated in memory of Frances Lee (F. LEE).

## Funding

National Institutes of Health grant GM7197-42 (UL)

National Institutes of Health grant 1R01GM116113-01A1 (ML)

National Science Foundation MCB2020667 (ML)

National Institutes of Health postdoctoral fellowship F32GM146423 (DA)

National Institutes of Health grant R01-GM115523 (PA)

National Institutes of Health MIRA R35GM133780 (LZhao)

## Author contributions

Conceptualization: UL, ML

Methodology: UL, ML

Investigation: UL, DA, SX, MA, DRS, IE, DS, JC, PR, NS, CL, CBL, JJE, LZhang

Visualization: UL, DA, PR

Funding acquisition: PA, QZ, LZhao, ML

Project administration: UL, PA, QZ, LZhao, ML

Supervision: PA, QZ, JJE, LZhao, ML

Writing – original draft: UL, DA, AG, ML

Writing – review & editing: UL, DA, PR, AG, NS, LZhao, ML

## Data and materials availability

Publicly available *D. melanogaster* Hi-C libraries form S2 cells: PRJNA393992. Newly generated *D. yakuba* Hi-C libraries from adult females: SRR12331759. Newly generated *D. pseudoobscura* and *D. miranda* Hi-C libraries from L3 larvae: PRJNA948678. Newly generated *D. yakuba* reference genome: PRJNA310215. Publicly available *D. pseudoobscura* reference genome: https://www.ryanbracewell.com/data.html. Publicly available *D. miranda* reference genome: PRJNA474939. Newly generated *D. melanogaster* 4C-Seq data from L3 larve: PRJNA948431. scRNA-Seq data may: PRJNA995212.

## Supplementary Materials

Figs. S1-S9

File S1

Table S1

Movie S1

## SUPPLEMENTARY INFORMATION

### Prior Models and Genomic Symmetries

The first models describing new gene evolution proposed that all new genes likely evolve via duplication-based mechanisms (*1, 2*), including: the duplication, divergence, complementation (DDC)/sub-functionalization model (*3*), the escape from adaptive conflict (EAC) model (*4*), the innovation, amplification, and divergence (IAD) model. To address how a duplicate, redundant gene copy may rise to fixation, these models all assume multiple functions for any studied gene. In the DDC (sub-functionalization) model, symmetric (identical) copies of a duplicated gene lose function in complementary fashion, resulting in retention of duplicate gene copies with separate but complementary functions. While the DDC model allows each duplicate copy to possess a subset of the parental gene’s original functions, the EAC model allows for increased optimization of one or more of the original parental gene’s functions that are partitioned to each paralogous copy. The EAC model assumes internal genetic conflict within the parental gene preventing simultaneous optimization of its multiple functions, and duplication thus allows for the resolution of this evolutionary constraint, conferring a selective advantage in both parental and new genes. While the DDC and EAC models can explain how prior gene functions can be partitioned amongst duplicate copies, these models both assume that newly evolved duplicated genes can only retain pre-existing, essential functions from their parental genes and thus fail to describe a mechanism for how truly novel gene function emerges. In contrast, the IAD model proposes that changes in selection pressures may favor the increased expression of a given gene with an auxiliary function. This provides a selective advantage for increased gene dosage through an increase in gene copy number. Following the initial increase of auxiliary function through gene amplification, subsequent relaxation of selection pressure will allow for changes to accumulate on the various copies, allowing the new copies to diverge and potentially gain a new function (*5*). While the IAD model provides a solution for Ohno’s dilemma for gene family expansions in microbial organisms while encountering environmental changes (*6*), the model cannot be applied to metazoans due to often conflict effects for same genes in different tissues or cells.

A key factor missing in these previous models is the effect of chromosomal and regulatory context on a gene duplicate’s function and spatiotemporal expression. In the DDC, EAC and IAD models, the evolution of new gene duplicates is assumed to occur in a regulatory-independent context and do not describe how the regulatory sequences may shape the evolution of a new gene duplicate. Here, we explain how the regulatory context can promote neofunctionalization of newly duplicated genes through the enhancer-capture divergence (ECD) model. In the ECD model, the duplication of a pre-existing gene into a new regulatory context through a preexisting 3-dimenssional (3D) genome structure results in unique expression pattern from that of its parent gene controlled by a combination of regulatory elements from both the native and new contexts. The single-step evolutionary process of ECD thus allows for rapid neofunctionalization and is dependent on the regulatory architecture of the three-dimensional eukaryotic genome.

Similar to the IAD model, the ECD model first proposes that selective pressures change for the increased expression of a pre-existing (parental) gene within a specific tissue or set of tissues. To achieve this, there are two possible scenarios: 1) the evolution of a new enhancer in the parental gene’s locus, either through duplication or substitution, or in the case of the ECD model, 2) the duplication of the parental gene into a distal region of the genome that is already under the control of a pre-existing, tissue-specific enhancer. While the first scenario is possible, this would require multiple neutral *de novo* substitutions or insertions to generate one or more necessary transcription factor binding sites that fix within a population and modulates the expression of the new gene duplicate without disrupting parent gene’s expression pattern.

In the second scenario under *enhancer capture*, the duplication of the parental gene into another regulatory environment under the control of a pre-existing, tissue-specific enhancer is a solution that requires far fewer genomic changes and can occur in a single step. As the new selection pressures recur, the duplicate copy that is under new regulatory control will increase in frequency in the population, allowing it to fix. If the selection pressures change such that the increased tissue-specific expression of the new gene is no longer advantageous or compensatory mutations appear in the original parent locus, selective pressures will relax on the new gene copy allowing for *divergence*. While loss of the new gene copy by drift or negative selection is one possible fate, if the duplicate gene copy is at high enough frequency within a population, substitutions may accumulate and result in the gain of new, tissue-specific function.

There are several distinctions between the ECD model and previously classic models of gene duplication. First, the DDC, EAC and IAD models do not consider the effect of the pre-existing regulatory and chromosomal environment on a new, distally duplicated gene. Second, compared to the DDC and EAC models but like IAD, the ECD model is a single-step process in which the initial duplication event provides a selective advantage. However, unlike IAD, a duplicate gene copy can immediately integrate into a tissue-specific regulatory network separate from that of its parent under ECD, providing a fast evolutionary solution to “Ohno’s Dilemma.” A final and critical distinction between the ECD model and previous classical models, to address the dilemma, is that they explain the evolution and retention of different classes of gene duplications. The previous models are symmetric models of duplication-based evolution which assume that the original parental gene function is randomly partitioned or entirely retained between identical duplicate copies, making parent and new gene copies indistinguishable from one another. A similar genomic symmetry is also seen in tandem duplications, where duplicate copies cannot be definitively identified as the “parent” or “new” gene copy through synteny. As a result, the DDC, EAC, and IAD models provide reasonable mechanistic explanations for why a large number of duplicate gene copies are retained, applying particularly well to tandem and other symmetric gene duplications. However, these previous models do not consider the role of regulatory and chromosomal context on newly evolved, asymmetric duplicates and thus cannot explain the origination of a large number of evolutionarily important genes.

Genes evolving under ECD are asymmetric, as the parental gene remains in its original locus while the new copy resides in a distal region of the genome under the control of a different, pre-existing regulatory context. This genomic asymmetry allows for clear distinction between parent and new gene copies through synteny. A similar asymmetry is also seen in protein and regulatory function, where the parent gene retains its entire function and spatiotemporal expression pattern, while the auxiliary tissue-specific function and expression pattern is restricted to the new gene copy. The asymmetry of both 1) distinguishable gene identity and 2) segregation of expression and function is a key feature of the ECD model that distinguishes it from the DDC, EAC and IAC models, and allows for clear identification of genes that evolved under enhancer capture and the application of genomic tests regarding retention of essential gene function. We utilize these features of the ECD model to show a statistical enrichment of distally duplicated genes that have evolved via enhancer capture-divergence within *Drosophila melanogaster*. Under the ECD model, we predict that newly evolved genes will be enriched for two elements: 1) we predict high degrees of co-expression with neighboring genes combined with low co-expression with its parent gene and 2) we predict that genes evolving under ECD should originate as non-essential, as all essential function should be asymmetrically retained by the parent gene while the auxiliary, non-essential function is retained in the new copy. As the ECD process can occur in a single step in a 3D world of genome, we also predict that the enhancer capture process should be a key mechanism for the evolution of distally duplicated genes alongside the DDC, EAC, and IAD models due to its rapid evolvability. This is supported by the observation that gene duplication occurs more frequently than point mutation (*5, 6*), where ECD requires fewer genomic alterations than the *de novo* evolution of a new enhancer via substitution.

Under the IAD model, a full duplication of the parent gene function and expression pattern drives the duplicate copies to fixation as it provides the most evolvable solution to new conditions. In contrast, under the enhancer capture-divergence model, a copy of the parent gene duplicates into a region of the genome containing an active enhancer(s) that modulates the new gene copy’s expression in a tissue-specific manner. Alternatively, the new gene may duplicate into an inactive region of the genome containing unbound transcription factor binding sites, thus activating a previously inert non-coding sequence into a *de novo* enhancer.

---

**Figure S1.**
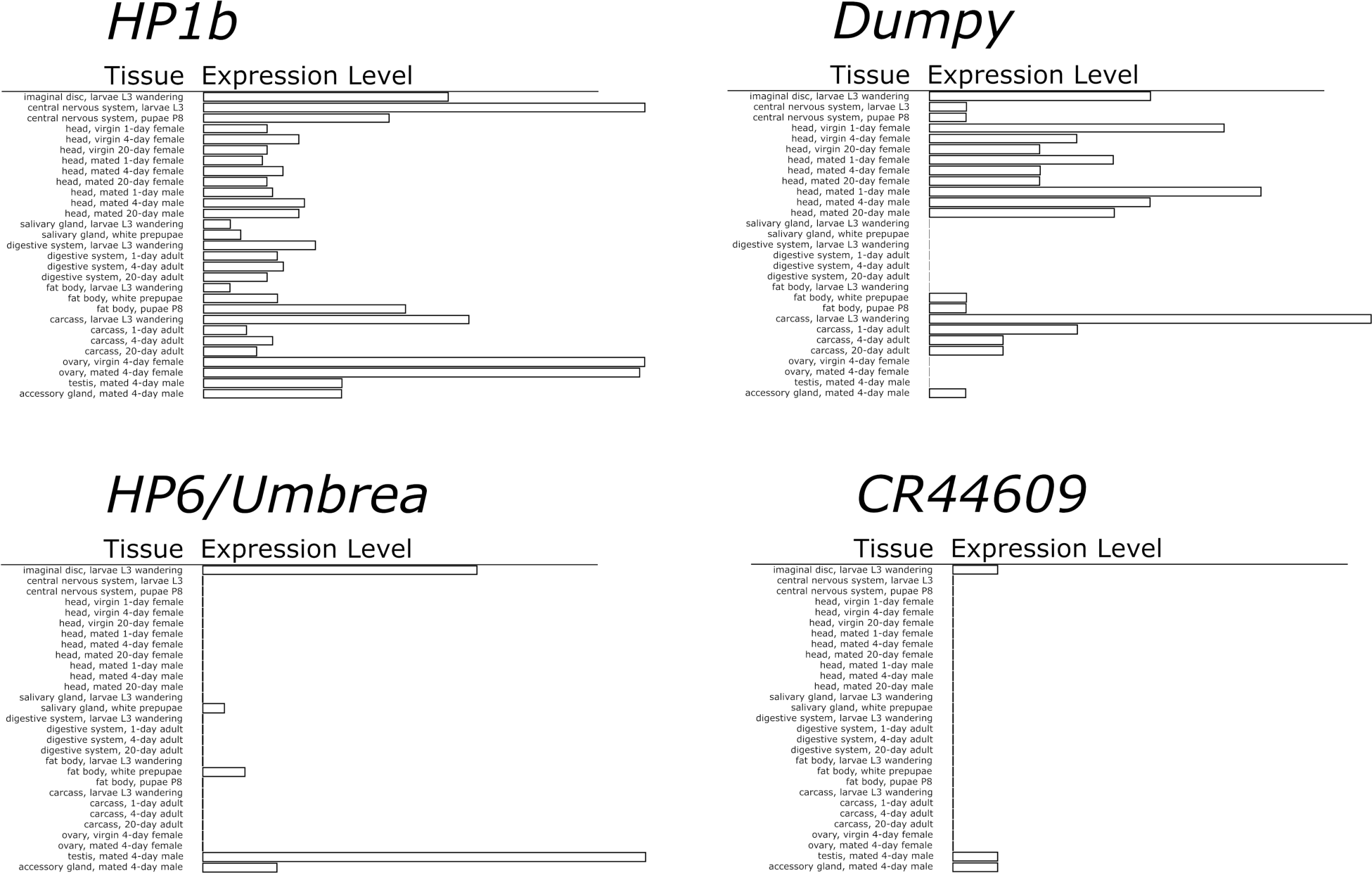
Expression patterns of *HP6/Umbrea* and other genes. Unlike the broad expression pattern of parental gene *HP1b*, the tissue expression pattern of *HP6/Umbrea* is stereotypical of new gene expression patterns, with high tissue specificity, restricted in this case to primarily the imaginal discs, larval salivary glands, and male reproductive organs. While *HP6/Umbrea* was inserted into an intronic region of the larger gene *dumpy, HP6/Umbrea*’s expression pattern is shared with *HP6/Umbrea*’s neighboring gene *CR44609*.

**Figure S2.**
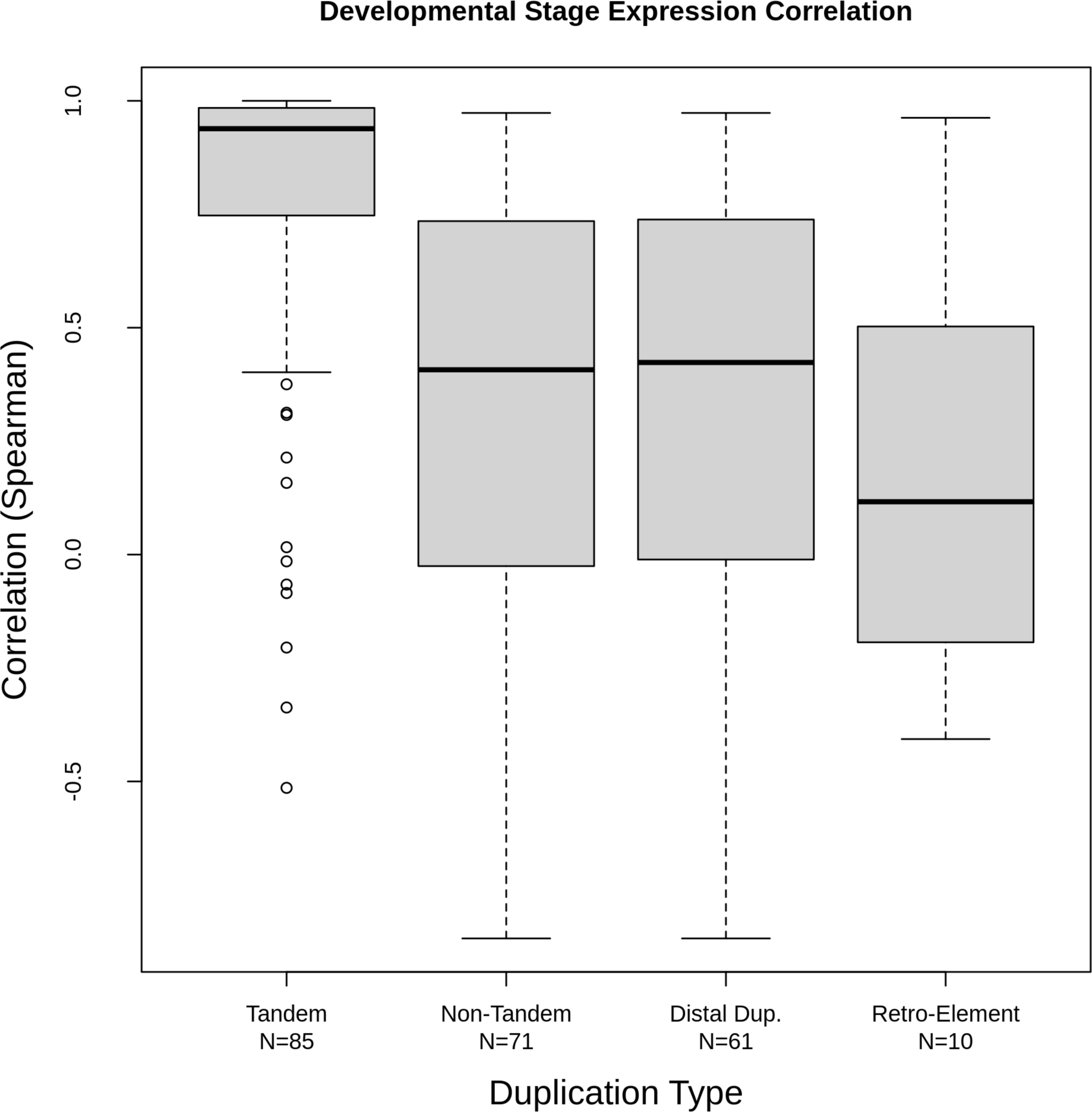
Enhancer Capture-Divergence drives regulatory neo-functionalization of new duplicate genes. Parental and new gene co-expression for new duplicate genes arising by tandem, distal duplicates, retro-transposons, and non-tandem (distal + retro-transposons) duplicates were calculated using gene expression data for 30 developmental stages in *D. melanogaster* (“developmental co-expression”). The development co-expression of non-tandem duplicates was significantly lower than the developmental co-expression of tandem duplicates (p=3.45 x 10^-10^) as well as distal duplicates and retro-transposons alone (distal: p=8.99 x 10^-9^, retro-transposition: p=5.41 x 10^-3^). These combined results demonstrate how Enhancer Capture-Divergence is a significant driver of regulatory neo-functionalization in new duplicate genes, which cannot be explained by symmetric models of new duplicate gene evolution.

**Figure S3.**
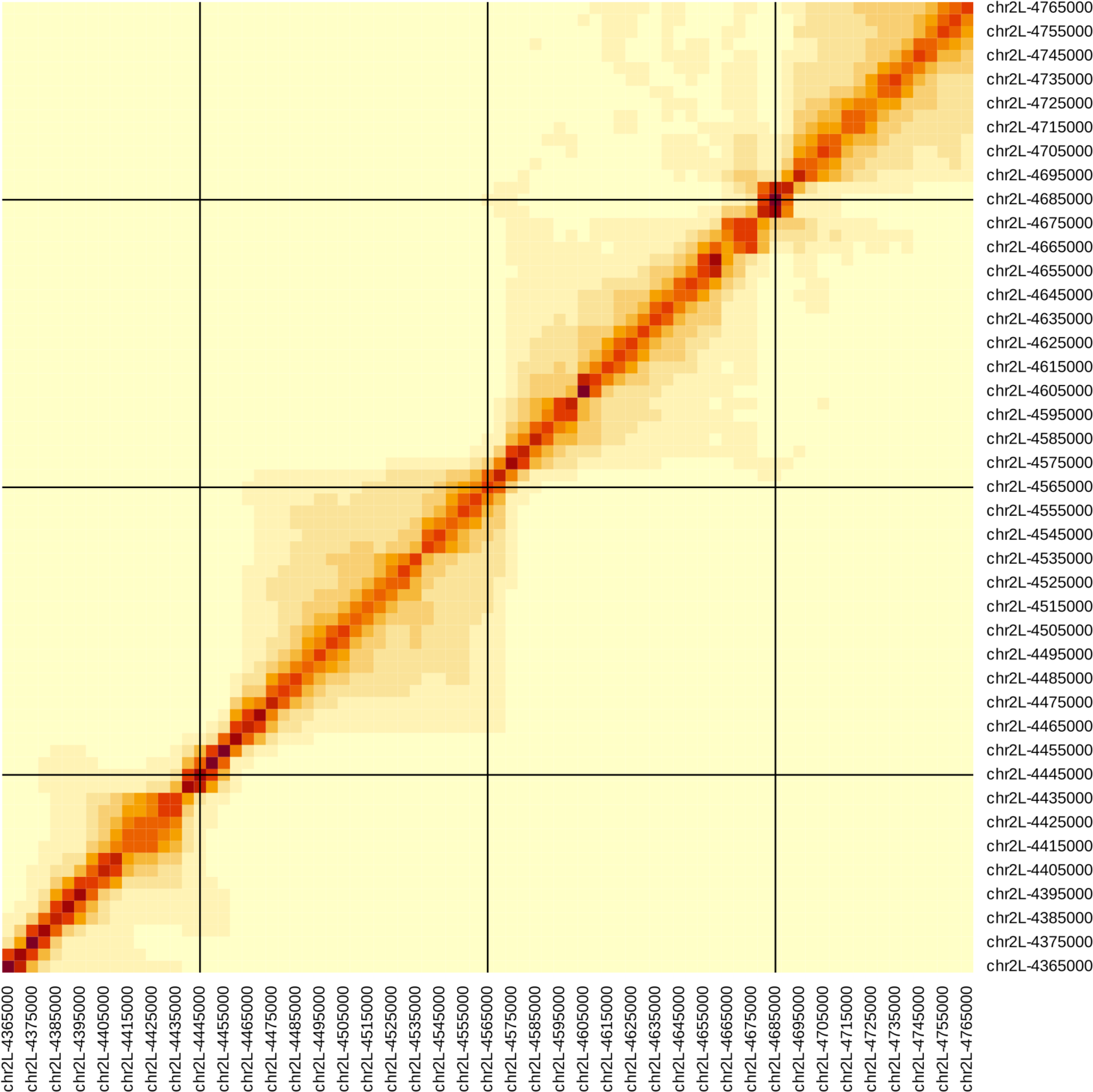
Local Hi-C heatmap for *D. melanogaster.* Shown above is the local chromosomal configuration of chromosome 2L in the vicinity of *HP6/Umbrea* (chr2L:4570000, center).

**Figure S4.**
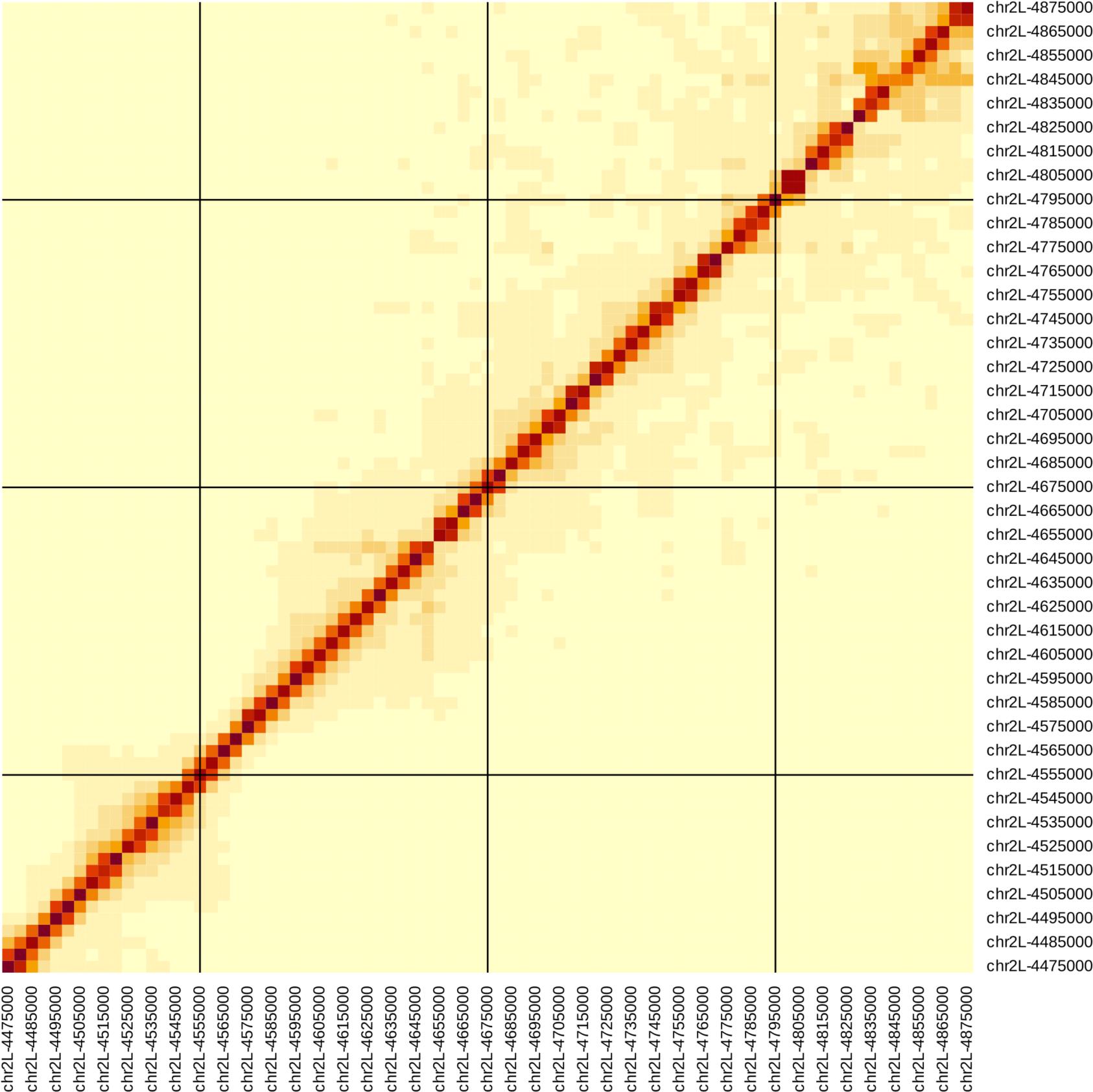
Local Hi-C heatmap for *D. yakuba*. Shown above is the local chromosomal configuration of chromosome 2L in the vicinity of *HP6/Umbrea* (chr2L:4680000, center).

**Figure S5.**
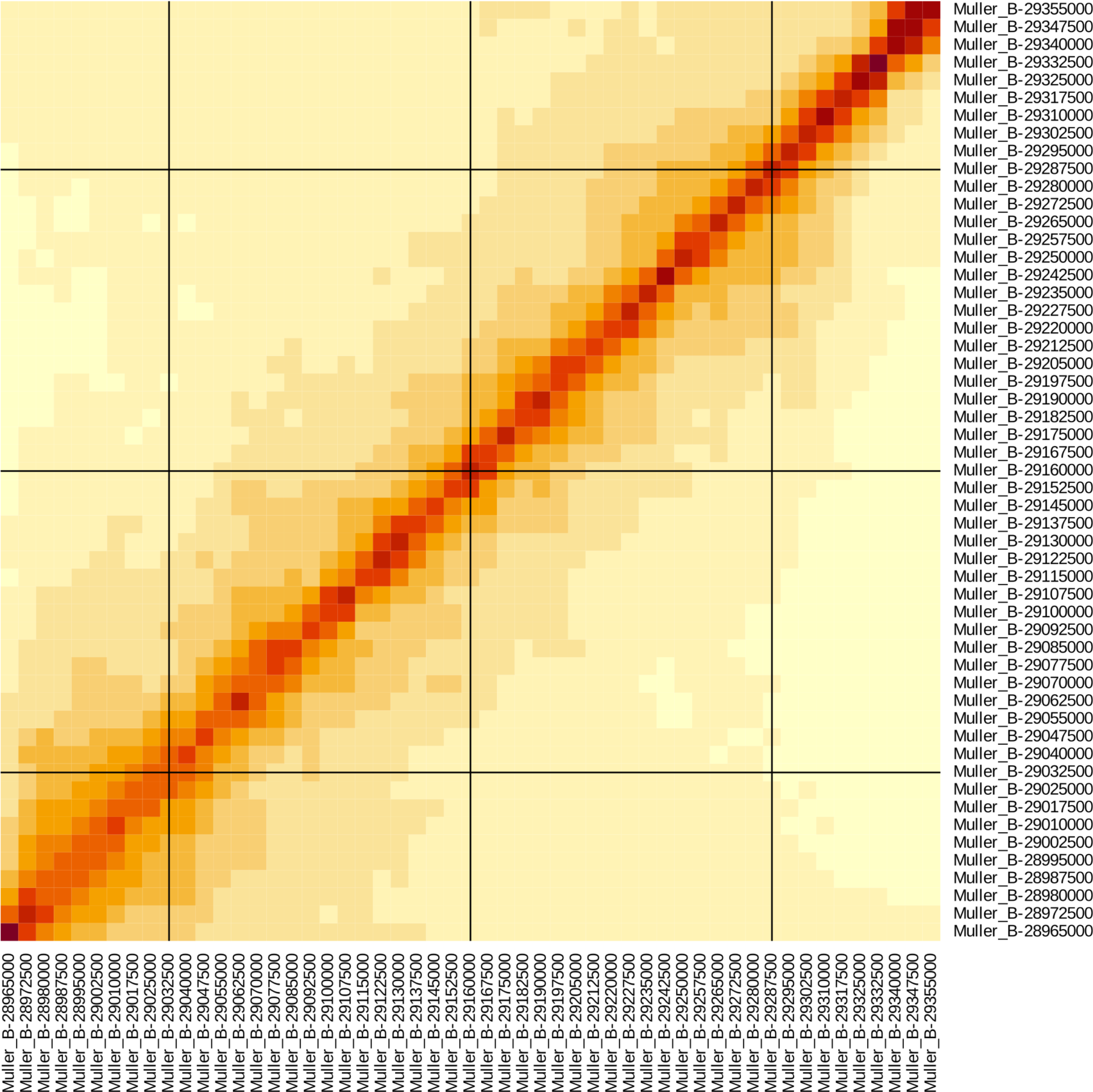
Local Hi-C heatmap for *D. pseudoobscura*. Shown above is the local chromosomal configuration of Muller Element B in the vicinity of *HP6/Umbrea*’s future insertion site (Muller B:29165000, center). Note a large-scale chromosomal inversion event occurred between *D. melanogaster* and *D. pseudoobscura* (not shown).

**Figure S6.**
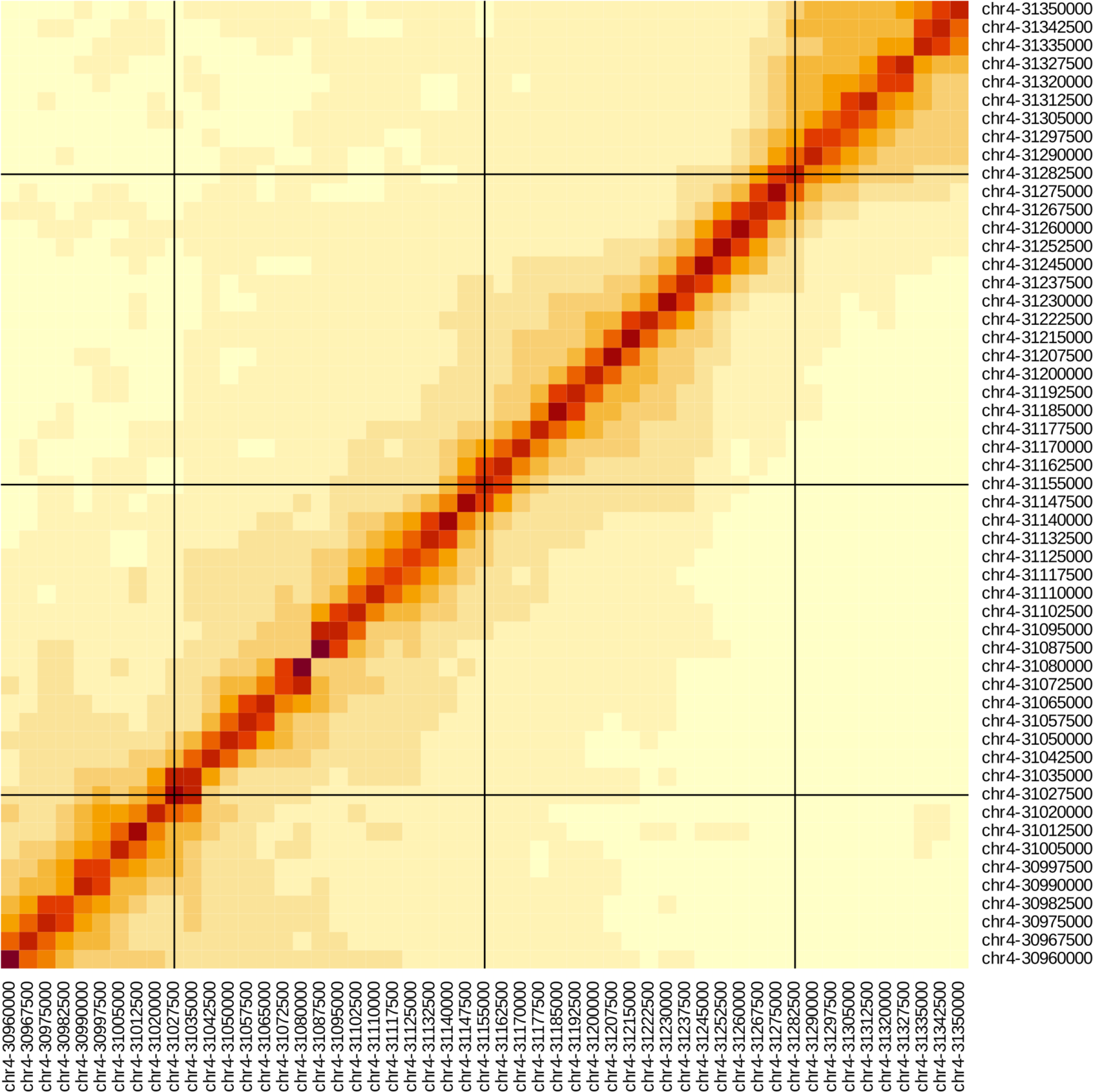
Local Hi-C heatmap for *D. miranda*. Shown above is the local chromosomal configuration of chromosome 4 in the vicinity of *HP6/Umbrea*’s future insertion site (chr4:31160000, center).

**Figure S7.**
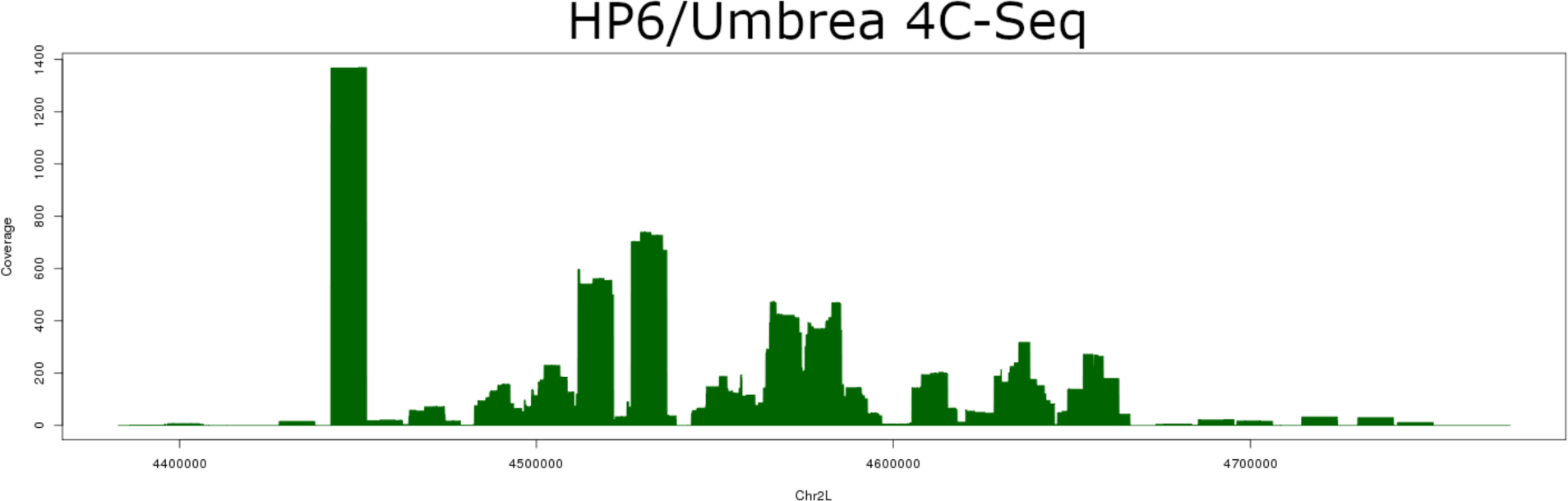
4C-Seq in *D. melanogaster*. Shown above are raw read coverage results from 4C-Seq derived from *D. melanogaster* larval tissue with self-self interactions removed, centered on *HP6/Umbrea*. The strong peak on the left shows the location of FLEE1.

**Figure S8.**
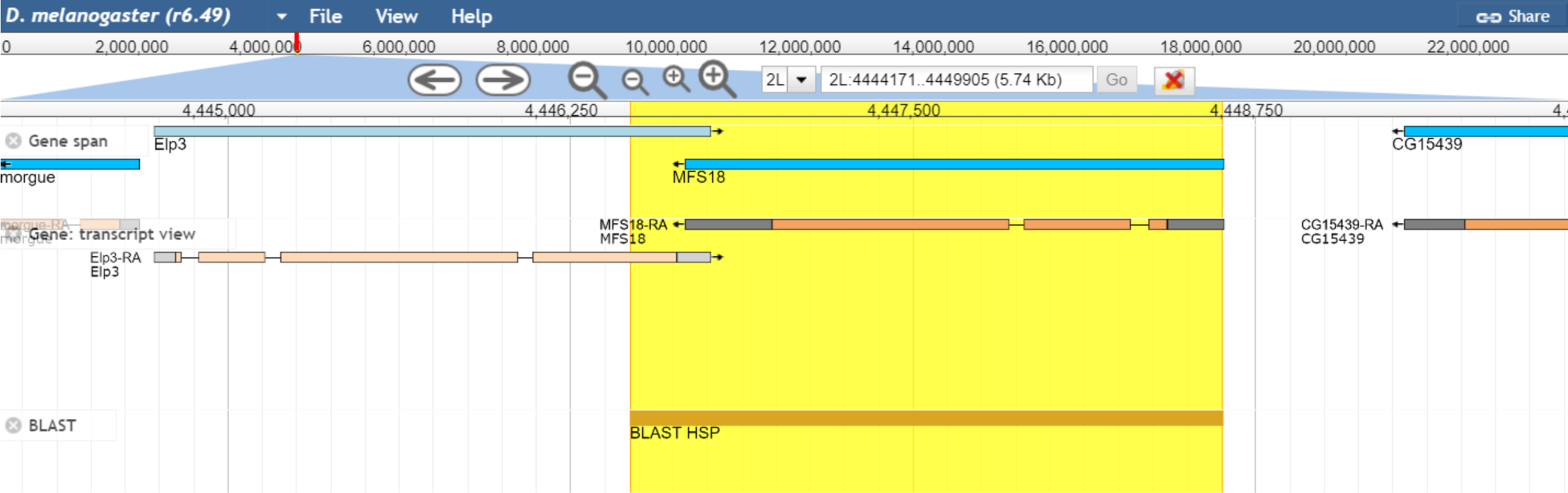
FLEE1 is located within exonic sequence. The FlyBase gene track and transcript view for FLEE1 shows that it is contained nearly entirely within the coding sequences of *MFS18* and *Elp3* on chromosome 2L.

**Figure S9.**
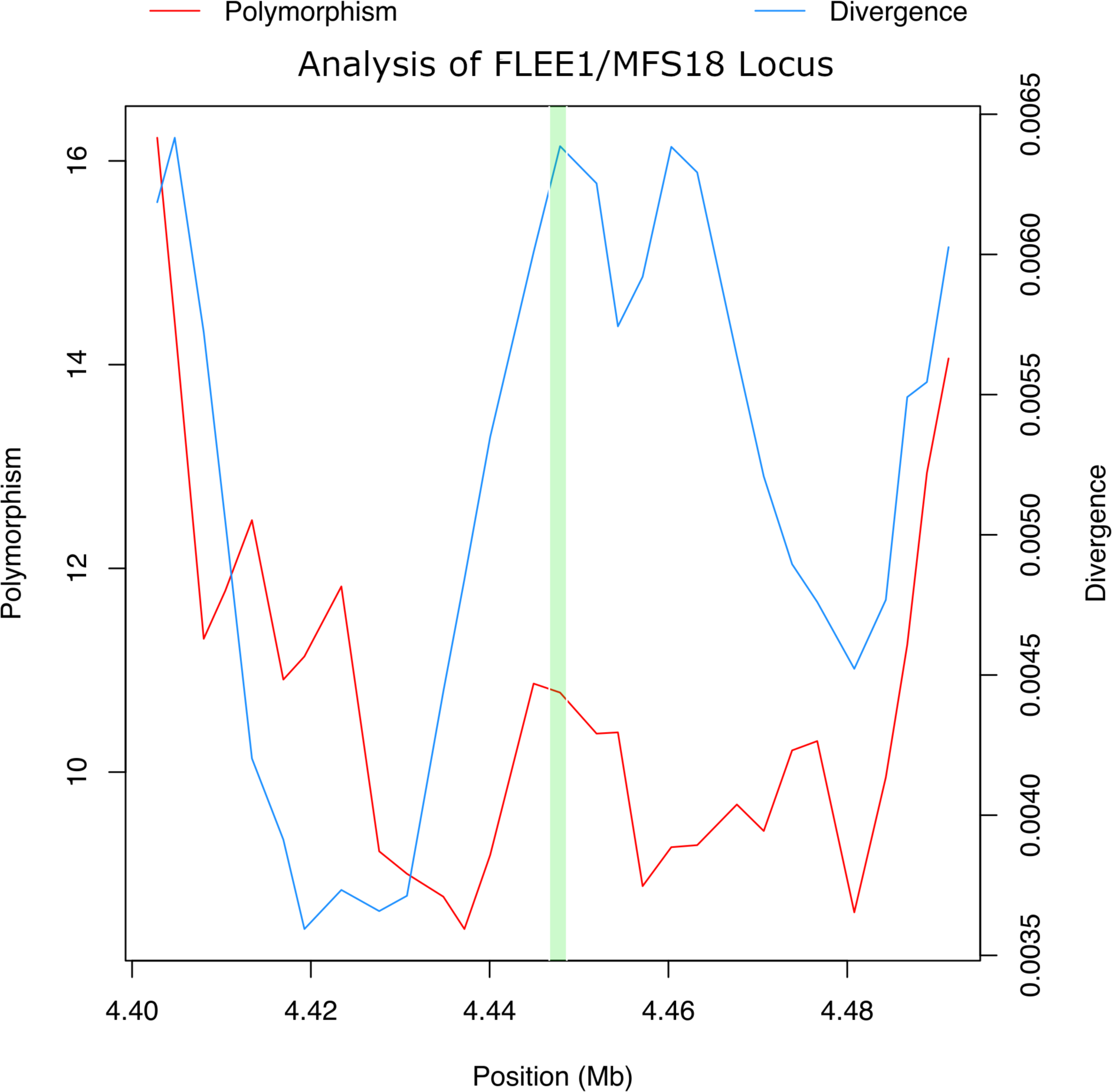
Polymorphism and divergence in FLEE1/*MFS18* locus. Polymorphism and divergence calculations on chromosome 2L with a 2.5kb window, with the *MFS18* locus highlighted in green, show how the relative number of polymorphisms is low when compared to sequence divergence.

**File S1. Sequence for FLEE1 enhancer.** Sequence for the cloned FLEE1 element in FASTA format.

>FLEE1

ACCAGGGCTTCGGTATGTTGCTGATGGAGGAGGCGGAGCGAATTGCTCGAGAGGAGCACGGCAGCAC

AAAACTGGCGGTCATATCGGGAGTGGGCACCAGAAACTACTATCGCAAAATGGGATACCAACTTGAC

GGACCCTACATGTCAAAGAGCATAGAAGAAAATAACTAGGTATAGCGTTAAATGACTGTCTTGGTGGT

ATGTTGGAGGATTAAATATTTGTATTTTATCACGCTAGGAGCAGTAAGATTTCGCTACTTAAAACTACT

CTCTTAAATATATACATTAATATATAGAATGAATCGATTTATTGGCTAAAACTCACAGGGTCCTTTAAA

GTATCAATGATACCACATATTTTTTGGCTTTAACATCTCACAAGAACAACTAAATGATCGTCAATCATA

AACGTGTATACTAAATAATATAAGCAGCATGAACATAAATCGATCCACTCCAATATACCCCACACATA

AATAAATAGGTTAGTTTTTCTGAGGAAAGTGTGCAAGAGAATGTTAAACGATGGCTTCCGCCGAACCA

AAGACTATAAATATGATCCAGCCAACCAAATTGATGCCAGCAGCGGCGCTGAACACCATCGGCCAGC

TTTGTGTGAGCTCCAGAATGTGTCCGGCCAAGTATACTCCGAGAAAGCCAGGAATCGCGCCCACTGTG

TTCATCAGGCCAAAGACGCTGCCCGAATGCAGAGGTGCCAGGTCTTGGGGATTCACTGTTACCGCGTT

GTTGTGGAAGCCCGTGCCGCCAATGATAATGGTCATGCAGATGAGCGCCGTATGGAAGTCCGAGGTG

CGGCTCATCACAAACAGGGCCAGATTCTGAGCGGCAAAGCAGCAACTTTGGATGACCTTGCGCACCGT

CGTCGTGTGCCATTCGCGAGCGAGTAATCTGGTGGTCAAGTACTTGGCGAATAGCGTGCACGGTGGCA

GGGCAAGCCACGGGATCATGTTCACTACCCAACCCTTGGCGTGTGGAAAGCCGTCGTGGAAGTATGTA

GGCAGCCAGGAGAGTAGCACGAAGAAGCAGTTCATCTCGCAGGCGTGAGTCAGCACACAGGCCCAGA

AGGACAGCCTACGAAAGTATCGCAACCAAGGCACGGCTGACGTCTCTGCCGGACTCTTGTTCGCGCAC

AGTCGGGATGGCGTGGCAATATTAATGATTCGGTTTCGCTCGCCGGCCATTGCATAGTAGCGCAGCAC

CAGCGCCCATGCGATGCCCATCAGTCCTATCACCCGGAATACATACGACCAGCCGAAGTAGTCCAGCA

GAAAAGATCCCATAATCCCAGTCAGAAGAGTACCTAGAGCCGATCCCGCTGTGAGCAGCCCAAAGAA

GCTGCTTCTCTCATTGGGGCACAAATTCTGCAAACGATTAAGTTATAGTTTATGTGTAAATTTATAAAA

TTAGCTAAGCACCTGACTGGTTAGACTAATCATGCTAGGAAAGTGCACGCCCTGAAGAGCGCCGTTCA

GGATTCGAATGGCAACAATGAAGGGAATAGCGTAACTCTTGATGGAGCCCGCCGTCCAGATGATAGT

GGGCATTAGGAATGTGATAAGCGACCAGCCGATTGCGGCAAACAGAATGACTCGCTGGCCTCCAAAG

CGGTCGCTGAAGTAGCCGCCCACAACCTGCGTGAGTGTGTAGCCCCAGAAGAAGGAGCTGAGCACAG

TGCCCGAGTCGGTTTTGCTCCACTTTTGGGCGGATGCCACGGCCGGCACAAGAAGTGGCATAGTGGTG

CGGGTGGAGTACAGCATACAGGTGCCCGTAATAAGGGTGATGAACCAGACACGCTTCTCATGCCTGC

AATGATCCGACAAAGGAGTTGTACTTGGGGAGGTTTAGTGAGCTGTATGCTGTGAGACCCACCTGGTC

CAAATGCTCTGCGTGTCCACCAGTTCCCCGCGCAGCAGAGAATATTTTAGCTTCTCGTCCATGGTCACA

AACTGGGTCCGGAACTATTGCCTTTCCTTCACGTCACATATCAACTCCAACTGCTTCGTTGCTTGCCGC

TGTGGCATATTTTACTGCCCTTTGTTTACTTTCATTCACGTTGGCGACTAGACACGCCAAGTATTTGCG

CCTGTTAAAATTATGTTTTTACGTGGCCGTTTTTCCAACAGCCGCTGGACTAGAGCATAG

**Table S1.**
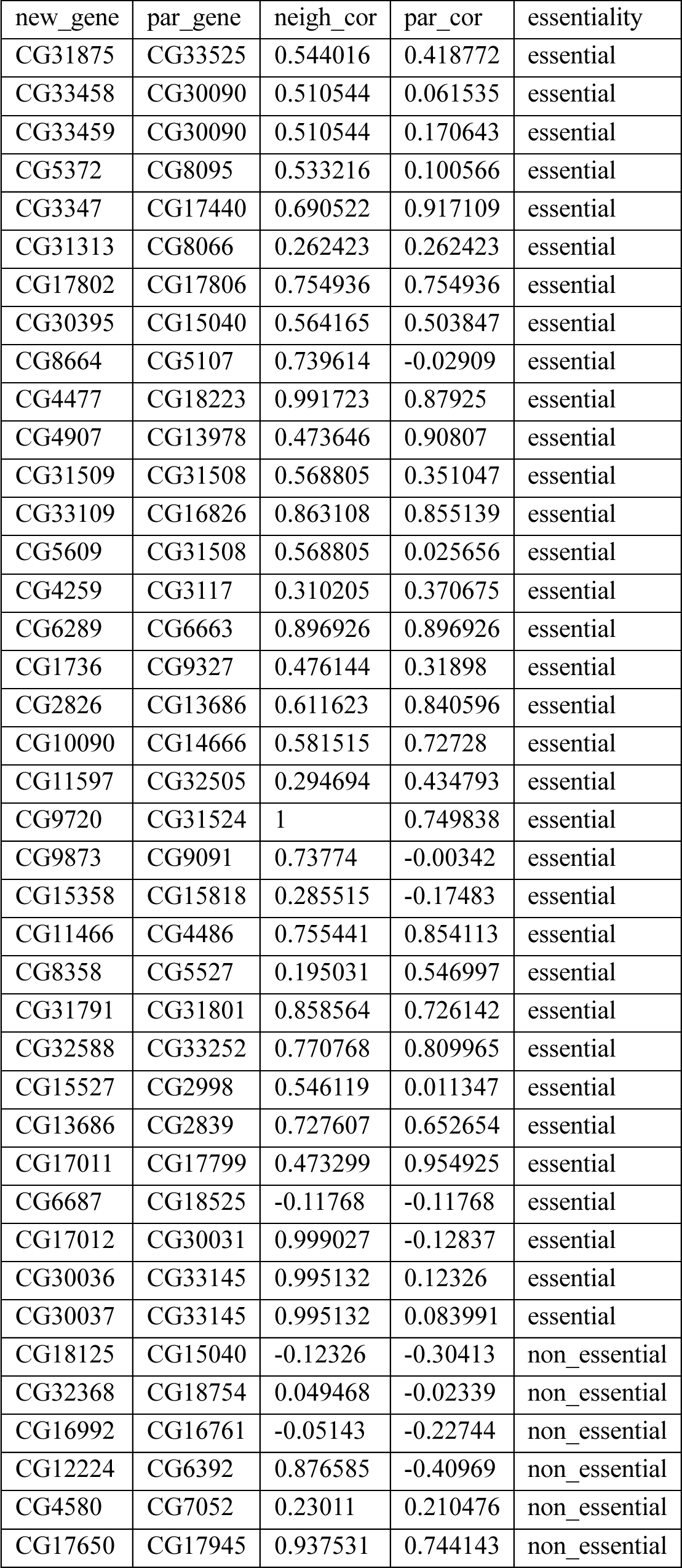
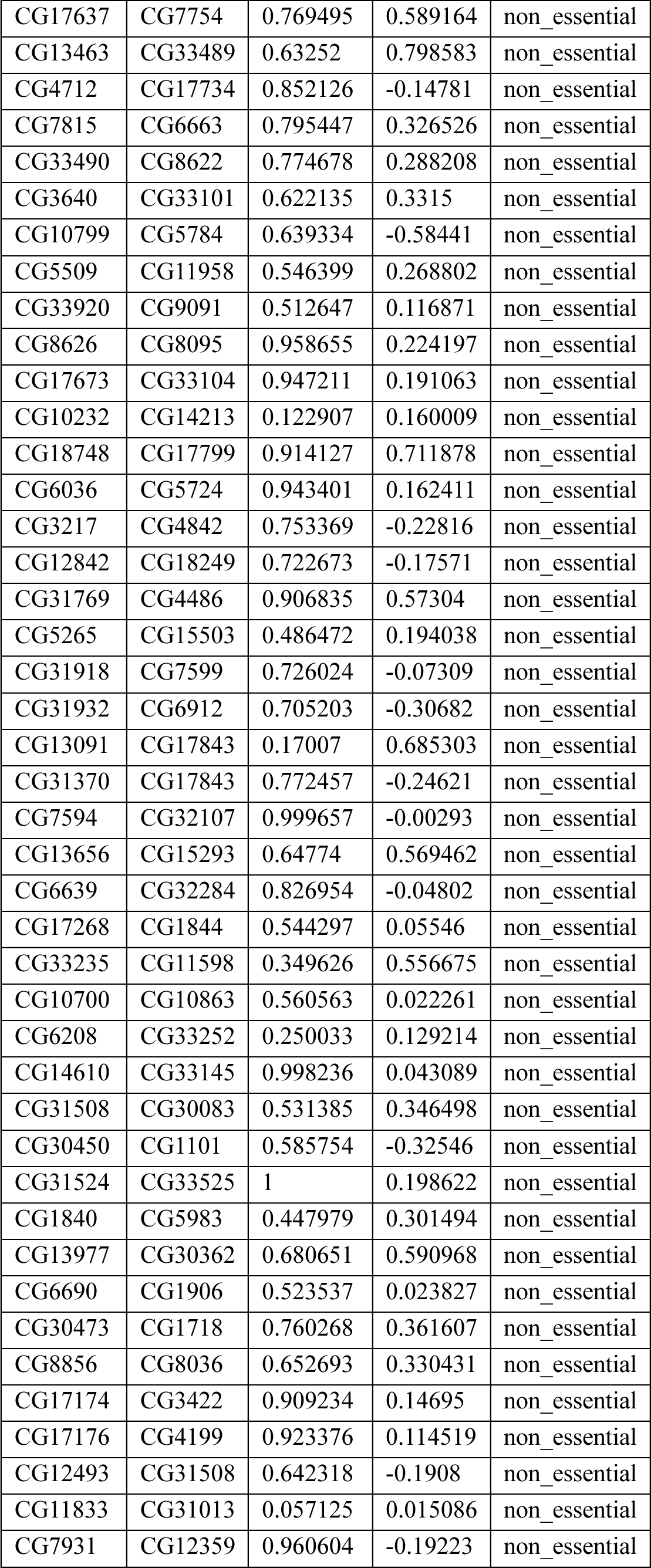
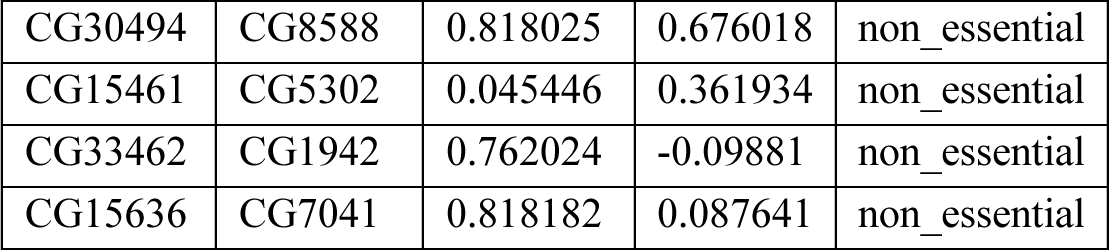
Co-expression and essentiality data for newly evolved distal duplicates. Data for each new gene/parent gene pair with parental coexpression, maximal neighboring coexpression, and essentiality data reported.

**Movie S1. Live GFP expression.** Video of live larvae with FLEE1 under the control of an enhancer-reporter vector. Expression is seen restricted to the larval salivary glands. (MovieS1.mp4)

## References and Notes

1. J. J. Emerson, M. Cardoso-Moreira, J. O. Borevitz, M. Long, Natural Selection Shapes Genome-Wide Patterns of Copy-Number Polymorphism in Drosophila melanogaster. Science 320, 1629–1631 (2008).

2. M. Kreitman, Nucleotide polymorphism at the alcohol dehydrogenase locus of Drosophila melanogaster. Nature 304, 412–417 (1983).

3. Y. Wang, P. McNeil, R. Abdulazeez, M. Pascual, S. E. Johnston, P. D. Keightley, D. J. Obbard, Variation in mutation, recombination, and transposition rates in Drosophila melanogaster and Drosophila simulans. Genome Res. 33, 587–598 (2023).

4. U. Bergthorsson, D. I. Andersson, J. R. Roth, Ohno’s dilemma: Evolution of new genes under continuous selection. Proc. Natl. Acad. Sci. USA 104, 17004–17009 (2007).

5. A. Force, M. F. Lynch, B. Pickett, A. Amores, Y. Yan, J. Postlethwait, Preservation of Duplicate Genes by Complementary, Degenerative Mutations. Genetics 151, 1531–1545 (1999).

6. C. T. Hittinger, S. B. Carroll, Gene duplication and the adaptive evolution of a classic genetic switch. Nature 449, 677–681 (2007).

7. J. Nasvall, L. Sun, J. R. Roth, D. I. Andersson, Real-Time Evolution of New Genes by Innovation, Amplification, and Divergence. Science 338, 384–387 (2012).

8. J. B. Brown, N. Boley, R. Eisman, G. E. May, M. H. Stoiber, M. O. Duff, B. W. Booth, J. Wen, S. Park, A. M. Suzuki, K. H. Wan, C. Yu, D. Zhang, J. W. Carlson, L. Cherbas, B. D. Eads, D. Miller, K. Mockaitis, J. Roberts, C. A. Davis, E. Frise, A. S. Hammonds, S. Olson, S. Shenker, D. Sturgill, A. A. Samsonova, R. Weiszmann, G. Robinson, J. Hernandez, J. Andrews, P. J. Bickel, P. Carninci, P. Cherbas, T. R. Gingeras, R. A. Hoskins, T. C. Kaufman, E. C. Lai, B. Oliver, N. Perrimon, B. R. Graveley, S. E. Celniker, Diversity and dynamics of the Drosophila transcriptome. Nature 512, 393–399 (2014).

9. A. Larkin, S. Marygold, G. Antonazzo, H. Attrill, G. dos Santos, P. Garapati, J. Goodman, L. Gramates, G. Millburn, V. Strelets, C. Tabone, J. Thurmond, T. F. Consortium, FlyBase: updates to the Drosophila melanogaster knowledge base. Nucleic Acid Research 49, D899–D907 (2021).

10. T. Sexton, E. Yaffe, E. Kenigsberg, F. Bantignies, B. Leblanc, M. Hoichman, H. Parrinello, A. Tanay, G. Cavalli, Three-Dimensional Folding and Functional Organization Principles of the Drosophila Genome. Cell 148, 458– 472 (2012).

11. G. Fasano, A. Franceschini, A multidimensional version of the Kolmogorov–Smirnov test. Monthly Notices of the Royal Astronomical Society 225, 155–170 (1987).

12. C. Puritz, E. Ness-Cohn, R. Braun, fasano.franceschini.test: An Implementation of a Multidimensional KS Test in R. arXiv arXiv:2106.10539 [Preprint] (2022). 10.48550/arXiv.2106.10539.

13. Q. Zhou, G. Zhang, Y. Zhang, S. Xu, R. Zhao, Z. Zhan, X. Li, Y. Ding, S. Yang, W. Wang, On the origin of new genes in Drosophila. Genome Res., doi: 10.1101/gr.076588.108 (2008).

14. B. R. Graveley, A. N. Brooks, J. W. Carlson, M. O. Duff, J. M. Landolin, L. Yang, C. G. Artieri, M. J. van Baren, N. Boley, B. W. Booth, J. B. Brown, L. Cherbas, C. A. Davis, A. Dobin, R. Li, W. Lin, J. H. Malone, N. R. Mattiuzzo, D. Miller, D. Sturgill, B. B. Tuch, C. Zaleski, D. Zhang, M. Blanchette, S. Dudoit, B. Eads, R. E. Green, A. Hammonds, L. Jiang, P. Kapranov, L. Langton, N. Perrimon, J. E. Sandler, K. H. Wan, A. Willingham, Y. Zhang, Y. Zou, J. Andrews, P. J. Bickel, S. E. Brenner, M. R. Brent, P. Cherbas, T. R. Gingeras, R. A. Hoskins, T. C. Kaufman, B. Oliver, S. E. Celniker, The developmental transcriptome of Drosophila melanogaster. Nature 471, 473–479 (2011).

15. H. Kaessmann, N. Vinckenbosch, M. Long, RNA-based gene duplication: mechanistic and evolutionary insights. Nat Rev Genet 10, 19–31 (2009).

16. D. Jangam, C. Feschotte, E. Betran, Transposable Element Domestication As an Adaptation to Evolutionary Conflicts. Trends in Genetics 33 (2017).

17. B. D. Ross, L. Rosin, A. W. Thomae, M. A. Hiatt, D. Vermaak, A. F. A. de la Cruz, A. Imhof, B. G. Mellone, H. S. Malik, Stepwise evolution of essential centromere function in a Drosophila neogene. Science 240, 1211–1214 (2013).

18. F. Greil, E. de Wit, H. J. Bussemaker, B. van Steensel, HP1 controls genomic targeting of four novel heterochromatin proteins in Drosophila. EMBO Journal 26 (2007).

19. C. Joppich, S. Scholz, G. Korge, A. Schwendemann, Umbrea, a chromo shadow domain protein in Drosophila melanogaster heterochromatin, interacts with Hip, HP1 and HOAP. Chromosome Res 17, 19–36 (2009).

20. S. Chen, Y. E. Zhang, M. Long, New genes in Drosophila quickly become essential. Science 330, 1682–1685 (2010).

21. S. Xia, N. W. VanKuren, C. Chen, L. Zhang, C. Kemkemer, Y. Shao, H. Jia, U. Lee, A. S. Advani, A. Gschwent, M. D. Vibranovski, S. Chen, Y. E. Zhang, M. Long, Genomic analyses of new genes and their phenotypic effects reveal rapid evolution of essential functions in Drosophila development. PLoS Genetics 17, e1009654 (2021).

22. S. Grill, A. Riley, M. Selvaraj, R. Lehmann, HP6/Umbrea is dispensable for viability and fertility, suggesting essentiality of newly evolved genes is rare. Proceedings of the National Academy of Sciences 120, e2309478120 (2023).

23. S. E. Celniker, L. A. L. Dillon, M. B. Gerstein, K. C. Gunsalus, S. Henikoff, G. H. Karpen, M. Kellis, E. C. Lai, J. D. Lieb, D. M. MacAlpine, G. Micklem, F. Piano, M. Snyder, L. Stein, K. P. White, R. H. Waterston, modENCODE Consortium, Unlocking the secrets of the genome. Nature 459, 927–930 (2009).

24. Q. Wang, Q. Sun, D. M. Czajkowsky, Z. Shao, Sub-kb Hi-C in D. melanogaster reveals conserved characteristics of TADs between insect and mammalian cells. Nature Communications 9, 188 (2018).

25. C. A. Russo, N. Takezaki, M. Nei, Molecular phylogeny and divergence times of drosophilid species. Molecular Biology and Evolution 12, 391–404 (1995).

26. B. Tolhuis, M. Blom, R. M. Kerkhoven, L. Pagie, H. Teunissen, M. Nieuwland, M. Simonis, W. de Laat, M. van Lohuizen, B. van Steensel, Interactions among Polycomb Domains Are Guided by Chromosome Architecture. PLOS Genetics 7, e1001343 (2011).

27. H. J. G. van de Werken, P. J. P. de Vree, E. Splinter, S. J. B. Holwerda, P. Klous, E. de Wit, W. de Laat, 4C technology: protocols and data analysis. Methods Enzymol 513, 89–112 (2012).

28. B. E. Housden, K. Millen, S. J. Bray, Drosophila Reporter Vectors Compatible with PhiC31 Integrase Transgenesis Techniques and Their Use to Generate New Notch Reporter Fly Lines. G3 (Bethesda) 2, 79–82 (2012).

29. T. L. Bailey, M. Boden, F. A. Buske, M. Frith, C. E. Grant, L. Clementi, J. Ren, W. W. Li, W. S. Noble, MEME Suite: tools for motif discovery and searching. Nucleic Acids Research 37, W202–W208 (2009).

30. M. T. Weirauch, A. Yang, M. Albu, A. G. Cote, A. Montenegro-Montero, P. Drewe, H. S. Najafabadi, S. A. Lambert, I. Mann, K. Cook, H. Zheng, A. Goity, H. van Bakel, J.-C. Lozano, M. Galli, M. G. Lewsey, E. Huang, T. Mukherjee, X. Chen, J. S. Reece-Hoyes, S. Govindarajan, G. Shaulsky, A. J. M. Walhout, F.-Y. Bouget, G. Ratsch, L. F. Larrondo, J. R. Ecker, T. R. Hughes, Determination and Inference of Eukaryotic Transcription Factor Sequence Specificity. Cell 158, 1431–1443 (2014).

31. C. Zhang, J. Wang, W. Xie, G. Zhou, M. Long, Q. Zhang, Dynamic programming procedure for searching optimal models to estimate substitution rates based on the maximum-likelihood method. Proceedings of the National Academy of Sciences 108, 7860–7865 (2011).

32. R. R. Hudson, M. Kreitman, M. Aguadé, A Test of Neutral Molecular Evolution Based on Nucleotide Data. Genetics 116, 153–159 (1987).

33. J. H. McDonald, M. Kreitman, Adaptive protein evolution at the Adh locus in Drosophila. Nature 351, 652–654 (1991).

34. M. Z. Ludwig, C. Bergman, N. H. Patel, M. Kreitman, Evidence for stabilizing selection in a eukaryotic enhancer element. Nature 403, 564–567 (2000).

35. M. D. Vibranovski, Y. E. Zhang, C. Kemkemer, H. F. Lopes, T. L. Karr, M. Long, Re-analysis of the larval testis data on meiotic sex chromosome inactivation revealed evidence for tissue-specific gene expression related to the drosophila X chromosome. BMC Biology 10, 49 (2012).

36. W. Zhang, P. Landback, A. R. Gschwend, B. Shen, M. Long, New genes drive the evolution of gene interaction networks in the human and mouse genomes. Genome Biology 202 (2015).

37. M. Long, N. W. VanKuren, S. Chen, M. D. Vibranovski, New gene evolution: little did we know. Annual Review of Genetics 47, 307–333 (2013).

38. E. Witt, S. Benjamin, N. Svetec, L. Zhao, Testis single-cell RNA-seq reveals the dynamics of de novo gene transcription and germline mutational bias in Drosophila. eLife 8, e47138 (2019).

39. R. Y. Birnbaum, E. J. Clowney, O. Agamy, M. J. Kim, J. Zhao, T. Yamanaka, Z. Pappalardo, S. L. Clarke, A. M. Wenger, L. Nguyen, F. Gurrieri, D. B. Everman, C. E. Schwartz, O. S. Birk, G. Bejerano, S. Lomvardas, N. Ahituv, Coding exons function as tissue-specific enhancers of nearby genes. Genome Res. 22, 1059–1068 (2012).

40. S. Zhenilo, E. Khrameeva, S. Tsygankova, N. Zhigalova, A. Mazur, E. Prokhortchouk, Individual genome sequencing identified a novel enhancer element in exon 7 of the CSFR1 gene by shift of expressed allele ratios. Gene 566, 223–228 (2015).

41. W. Wang, J. Zhang, C. Alvarez, A. Llopart, M. Long, The origin of the Jingwei gene and the complex modular structure of its parental gene, yellow emperor, in Drosophila melanogaster. Molecular Biology and Evolution 17, 1294–1301 (2000).

42. W. Wang, H. Zheng, S. Yang, Y. Haijing, J. Li, H. Jiang, J. Su, L. Yang, J. Zhang, J. McDermott, R. Samudrala, J. Wang, H. Yang, J. Yun, K. Kristiansen, G. K. S. Wong, J. Wang, Origin and evolution of new exons in rodents. Genome Research 15, 1258–1264 (2005).

43. C. B. Bridges, The Bar “Gene” a Duplication. Science 83, 210–211 (1936).

44. H. J. Muller, Bar Duplication. Science 83, 528–530 (1936).

45. N. Harmston, E. Ing-Simmons, G. Tan, M. Perry, M. Markenschlager, B. Lenhard, Topologically associating domains are ancient features that coincide with Metazoan clusters of extreme noncoding conservation. Nature Communications 8, 441 (2017).

46. J. Krefting, M. A. Andrade-Navarro, J. Ibn-Salen, Evolutionary stability of topologically associating domains is associated with conserved gene regulation. BMC Biology, 87 (2018).

47. J. Y. Zhang, Q. Zhou, On the Regulatory Evolution of New Genes Throughout Their Life History. Molecular Biology and Evolution 36, 15–27 (2019).

48. H. Dai, T. F. Yoshimatsu, M. Long, Retrogene movement within- and between-chromosomes in the evolution of Drosophila genomes. Gene 385, 96–102 (2006).

49. L. E. Kursel, H. McConnel, A. F. A. de la Cruz, H. S. Malik, Gametic specialization of centromeric histone paralogs in Drosophila virilis. Life Science Alliance 4, e202000992 (2021).

50. N. W. VanKuren, M. Long, Gene Duplicates Resolving Sexual Conflict Rapidly Evolved Essential Gametogenesis Functions. Nature Ecology and Evolution 2, 705–712 (2018).

51. L. Zech, U. Haglund, K. Nilsson, G. Klein, Characteristic chromosomal abnormalities in biopsies and lymphoid-cell lines from patients with burkitt and non-burkitt lymphomas. International Journal of Cancer 17, 47–56 (1976).

52. V. K. Jain, J. G. Judde, E. E. Max, I. T. Magrath, Variable IgH chain enhancer activity in Burkitt’s lymphomas suggests an additional, direct mechanism of c-myc deregulation. J Immunol 150, 5418–5428 (1993).

53. B. Gryder, P. C. Scacheri, T. Ried, J. Khan, Chromatin Mechanisms Driving Cancer. Cold Spring Harb Perspect Biol 14, a040956 (2022).

54. J. Zheng, Oncogenic chromosomal translocations and human cancer (review). Oncol Rep 30, 2011–2019 (2013).

55. R. C. Hennessey, K. M. Brown, Cancer regulatory variation. Curr Opin Genet Dev 66, 41–49 (2021).

56. X. Wang, J. Xu, B. Zhang, Y. Hou, F. Song, H. Lyu, F. Yue, Genome-wide detection of enhancer-hijacking events from chromatin interaction data in rearranged genomes. Nat Methods 18, 661–668 (2021).

57. L. E. Montefiori, S. Bendig, Z. Gu, X. Chen, P. Pölönen, X. Ma, A. Murison, A. Zeng, L. Garcia-Prat, K. Dickerson, I. Iacobucci, S. Abdelhamed, R. Hiltenbrand, P. E. Mead, C. M. Mehr, B. Xu, Z. Cheng, T.-C. Chang, T. Westover, J. Ma, A. Stengel, S. Kimura, C. Qu, M. B. Valentine, M. Rashkovan, S. Luger, M. R. Litzow, J. M. Rowe, M. L. den Boer, V. Wang, J. Yin, S. M. Kornblau, S. P. Hunger, M. L. Loh, C.-H. Pui, W. Yang, K. R. Crews, K. G. Roberts, J. J. Yang, M. V. Relling, W. E. Evans, W. Stock, E. M. Paietta, A. A. Ferrando, J. Zhang, W. Kern, T. Haferlach, G. Wu, J. E. Dick, J. M. Klco, C. Haferlach, C. G. Mullighan, Enhancer Hijacking Drives Oncogenic BCL11B Expression in Lineage-Ambiguous Stem Cell Leukemia. Cancer Discovery 11, 2846–2867 (2021).

58. S. A. Tomlins, D. R. Rhodes, S. Perner, S. M. Dhanasekaran, R. Mehra, X.-W. Sun, S. Varambally, X. Cao, J. Tchinda, R. Kuefer, C. Lee, J. E. Montie, R. B. Shah, K. J. Pienta, M. A. Rubin, A. M. Chinnaiyan, Recurrent fusion of TMPRSS2 and ETS transcription factor genes in prostate cancer. Science 310, 644–648 (2005).

59. M. S. Litwin, H.-J. Tan, The Diagnosis and Treatment of Prostate Cancer: A Review. JAMA 317, 2532–2542 (2017).

60. K. J. Meaburn, T. Misteli, E. Soutoglou, Spatial genome organization in the formation of chromosomal translocations. Semin Cancer Biol 17, 80–90 (2007).

61. F. Dubois, N. Sidiropoulos, J. Weischenfeldt, R. Beroukhim, Structural variations in cancer and the 3D genome. Nat Rev Cancer 22, 533–546 (2022).

62. H. Neves, C. Ramos, M. G. da Silva, A. Parreira, L. Parreira, The nuclear topography of ABL, BCR, PML, and RARalpha genes: evidence for gene proximity in specific phases of the cell cycle and stages of hematopoietic differentiation. Blood 93, 1197–1207 (1999).

63. S. Kozubek, E. Lukásová, A. Marecková, M. Skalníková, M. Kozubek, E. Bártová, V. Kroha, E. Krahulcová, J. Slotová, The topological organization of chromosomes 9 and 22 in cell nuclei has a determinative role in the induction of t(9,22) translocations and in the pathogenesis of t(9,22) leukemias. Chromosoma 108, 426–435 (1999).

64. C. S. Osborne, L. Chakalova, J. A. Mitchell, A. Horton, A. L. Wood, D. J. Bolland, A. E. Corcoran, P. Fraser, Myc dynamically and preferentially relocates to a transcription factory occupied by Igh. PLoS Biol 5, e192 (2007).

65. S. Wingett, P. Ewels, M. Furlan-Magaril, T. Nagano, S. Schoenfelder, P. Fraser, S. Andrews, HiCUP: pipeline for mapping and processing Hi-C data. F100Res 4, 1310 (2015).

66. B. Langmead, S. L. Salzberg, Fast gapped-read alignment with Bowtie 2. Nature Methods 9, 357–359 (2012).

67. R. Bracewell, K. Chatla, M. J. Nalley, B. Bachtrog, Dynamic turnover of centromeres drives karyotype evolution in Drosophila. eLife Sep 16;8:e49002 (2019).

68. S. Mahajan, K. H.-C. Wie, M. J. Nalley, L. Gibilisco, D. Bachtrog, De novo assembly of a young Drosophila Y chromosome using single-molecule sequencing and chromatin conformation capture. PLoS Biology 16, e2006348 (2018).

69. S. Heinz, L. Taxari, M. G. B. Hayes, M. Urbanowski, M. W. Chang, N. Givarkes, A. Rialdi, K. M. White, R. A. Albrecht, L. Pache, I. Marazzi, A. Garcia-Sastre, M. L. Shaw, C. Benner, Transcription Elongation Can Affect Genome 3D Structure. Cell 174, 1522–1536 (2018).

70. N. C. Durand, M. S. Shamim, I. Machol, S. S. P. Rao, M. H. Huntley, E. S. Lander, E. L. Aiden, Juicer provides a one-click system for analyzing loop-resolution Hi-C experiments. Cell Systems 3 (2016).

71. J. M. Coughlan, A. J. Dagilis, A. Serrato-Capuchina, H. Elias, D. Peede, K. Isbell, D. M. Castillo, B. S. Cooper, D. R. Matute, Patterns of Population Structure and Introgression Among Recently Differentiated Drosophila melanogaster Populations. Molecular Biology and Evolution 39, msac223 (2022).

72. S. Chen, Y. Zhou, Y. Chen, J. Gu, fastp: an ultra-fast all-in-one FASTQ preprocessor. Bioinformatics 34, i884– i890 (2018).

73. H. Li, R. Durbin, Fast and accurate short read alignment with Burrows–Wheeler transform. Bioinformatics 25, 1754–1760 (2009).

74. P. Cingolani, A. Platts, L. L. Wang, M. Coon, T. Nguyen, L. Wang, S. J. Land, X. Lu, D. M. Ruden, A program for annotating and predicting the effects of single nucleotide polymorphisms, SnpEff. Fly 6, 80–92 (2012).

75. D. J. Begun, A. K. Holloway, K. Stevens, L. W. Hillier, Y.-P. Poh, M. W. Hahn, P. M. Nista, C. D. Jones, A. D. Kern, C. N. Dewey, L. Pachter, E. Myers, C. H. Langley, Population Genomics: Whole-Genome Analysis of Polymorphism and Divergence in Drosophila simulans. PLOS Biology 5, e310 (2007).

76. S. Purcell, B. Neale, K. Todd-Brown, L. Thomas, M. A. R. Ferreira, D. Bender, J. Maller, P. Sklar, P. I. W. de Bakker, M. J. Daly, P. C. Sham, PLINK: A Tool Set for Whole-Genome Association and Population-Based Linkage Analyses. The American Journal of Human Genetics 81, 559–575 (2007).

77. D. Kuznetsov, F. Tegenfeldt, M. Manni, M. Seppey, M. Berkeley, E. V. Kriventseva, E. M. Zdobnov, OrthoDB v11: annotation of orthologs in the widest sampling of organismal diversity. Nucleic Acids Res 51, D445–D451 (2023).

78. F. Abascal, R. Zardoya, M. J. Telford, TranslatorX: multiple alignment of nucleotide sequences guided by amino acid translations. Nucleic Acids Res 38, W7–13 (2010).

79. K. Katoh, D. M. Standley, MAFFT Multiple Sequence Alignment Software Version 7: Improvements in Performance and Usability. Molecular Biology and Evolution 30, 772–780 (2013).

80. Z. Yang, PAML 4: phylogenetic analysis by maximum likelihood. Mol Biol Evol 24, 1586–1591 (2007).

81. M. Kimura, The Neutral Theory of Molecular Evolution (Cambridge University Press, Cambridge, 1983; https://www.cambridge.org/core/books/neutral-theory-of-molecular-evolution/0FF60E9F47915B17FFA2620C49400632).

82. D. L. Hartl, A. G. Clark, Principles of Population Genetics (Oxford University Press, Oxford, New York, Fourth Edition, Fourth Edition., 2006).

83. D. A. Briscoe, J. M. Malpica, A. Robertson, G. J. Smith, R. Frankham, R. G. Banks, J. S. F. Barker (1992). Rapid loss of genetic variation in large captive populations of Drosophila flies: Implications for genetic management of captive populations. Conservation Biology 6, 416–425 (1992).

84. R. Frankham, Effective population size/adult population size ratios in wildlife: a review. Genetics Research 66, 95–107 (1995).

85. A. J. Tarashansky, J. M. Musser, M. Khariton, P. Li, D. Arendt, S. R. Quake, B. Wang, Mapping single-cell atlases throughout Metazoa unravels cell type evolution. eLife 10, e66747 (2021).

86. E. W. Green, G. Fedele, F. Giorgini, C. P. Kyriacou, A Drosophila RNAi collection is subject to dominant phenotypic effects. Nat Methods 11, 222–223 (2014).

## References

1. S. Ohno, Evolution by Gene Duplication (Springer, New York, 1970).

2. H. J. Muller, Bar Duplication. Science. 83, 528–530 (1936).

3. A. Force, M. F. Lynch, B. Pickett, A. Amores, Y. Yan, J. Postlethwait, Preservation of Duplicate Genes by Complementary, Degenerative Mutations. Genetics. 151, 1531–1545 (1999).

4. C. T. Hittinger, S. B. Carroll, Gene duplication and the adaptive evolution of a classic genetic switch. Nature. 449, 677–681 (2007).

5. U. Bergthorsson, D. I. Andersson, J. R. Roth, Ohno’s dilemma: Evolution of new genes under continuous selection. Proc. Natl. Acad. Sci. USA. 104, 17004–17009 (2007).

6. J. Nasvall, L. Sun, J. R. Roth, D. I. Andersson, Real-Time Evolution of New Genes by Innovation, Amplification, and Divergence. Science. 338, 384–387 (2012).

