## SUPPLEMENTARY INFORMATION for "The 3-Dimensional Genome Drives the Evolution of Asymmetric Gene Duplicates via Enhancer Capture-Divergence"

6.	J. Nasvall, L. Sun, J. R. Roth, D. I. Andersson, Real-Time Evolution of New Genes by Innovation, Amplification, and Divergence. *Science*. **338**, 384–387 (2012).

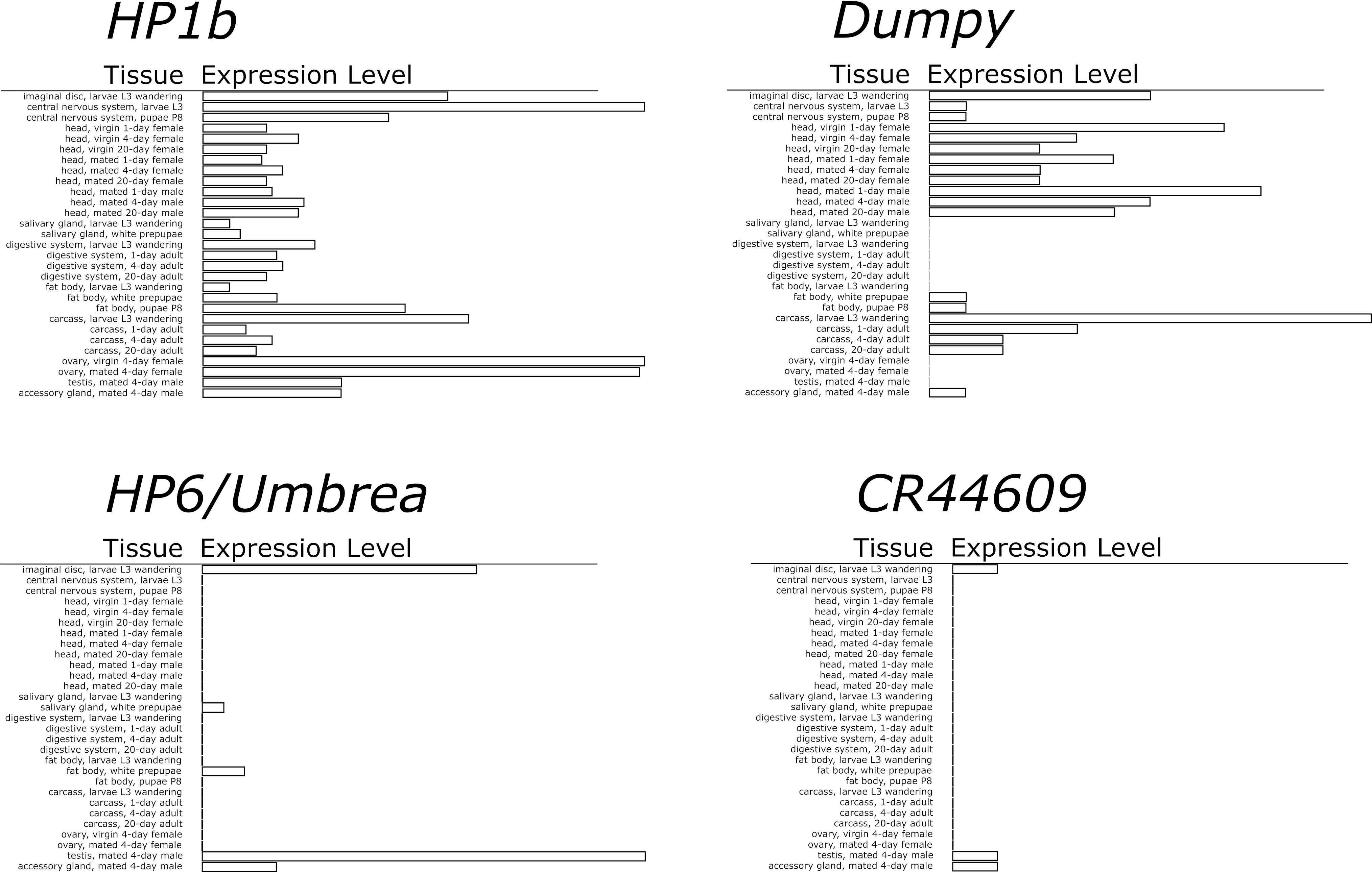

**Figure S1. Expression patterns of *HP6/Umbrea* and other genes.** Unlike the broad expression pattern of parental gene *HP1b*, the tissue expression pattern of *HP6/Umbrea* is stereotypical of new gene expression patterns, with high tissue specificity, restricted in this case to primarily the imaginal discs, larval salivary glands, and male reproductive organs. While *HP6/Umbrea* was inserted into an intronic region of the larger gene *dumpy*, *HP6/Umbrea*’s expression pattern is shared with *HP6/Umbrea*’s neighboring gene *CR44609*.

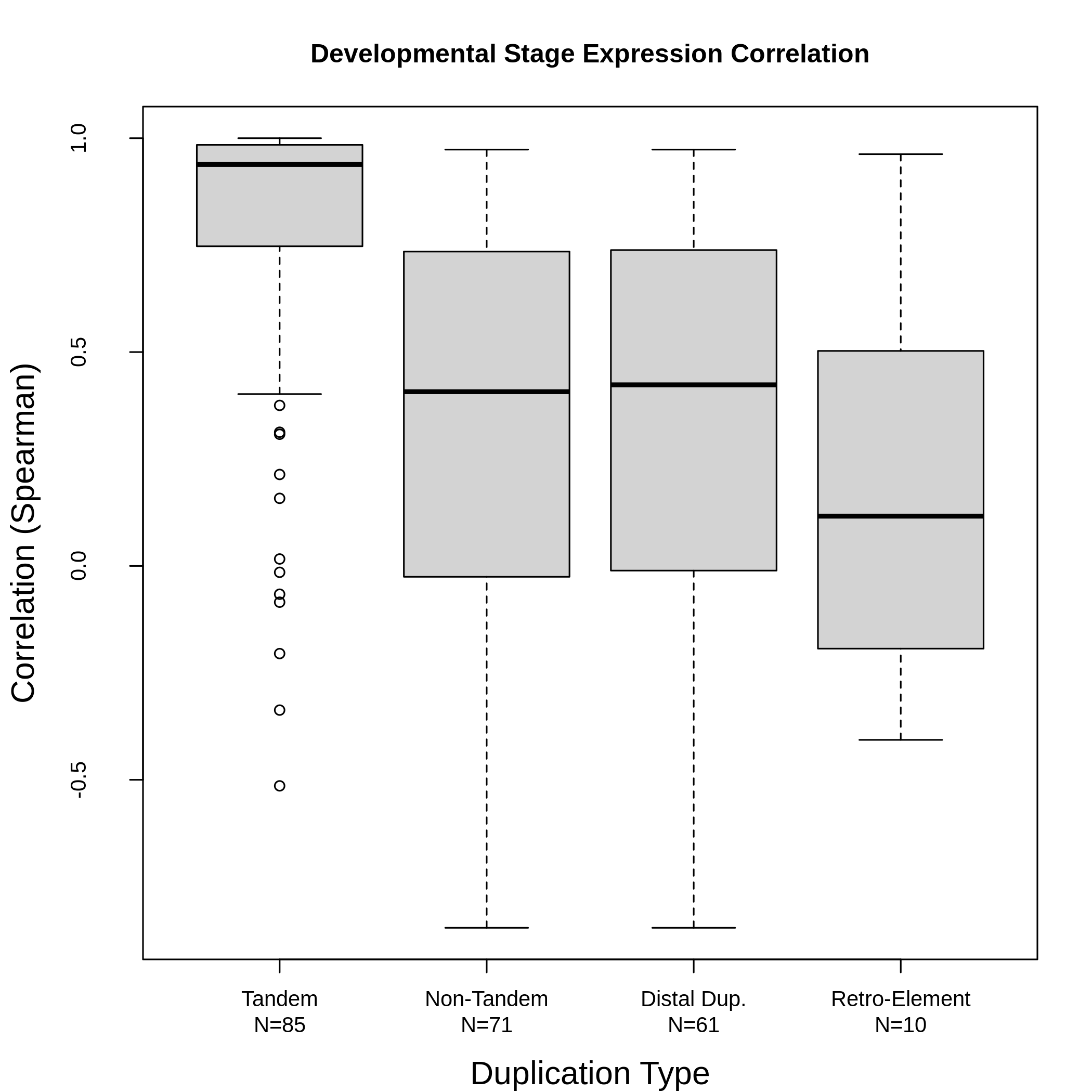

**Figure S2. Enhancer Capture-Divergence drives regulatory neo-functionalization of new duplicate genes.** Parental and new gene co-expression for new duplicate genes arising by tandem, distal duplicates, retro-transposons, and non-tandem (distal + retro-transposons) duplicates were calculated using gene expression data for 30 developmental stages in *D. melanogaster* (“developmental co-expression”). The development co-expression of non-tandem duplicates was significantly lower than the developmental co-expression of tandem duplicates (p=3.45 x 10^-10^) as well as distal duplicates and retro-transposons alone (distal: p=8.99 x 10^-9^, retro-transposition: p=5.41 x 10^-3^). These combined results demonstrate how Enhancer Capture-Divergence is a significant driver of regulatory neo-functionalization in new duplicate genes, which cannot be explained by symmetric models of new duplicate gene evolution.

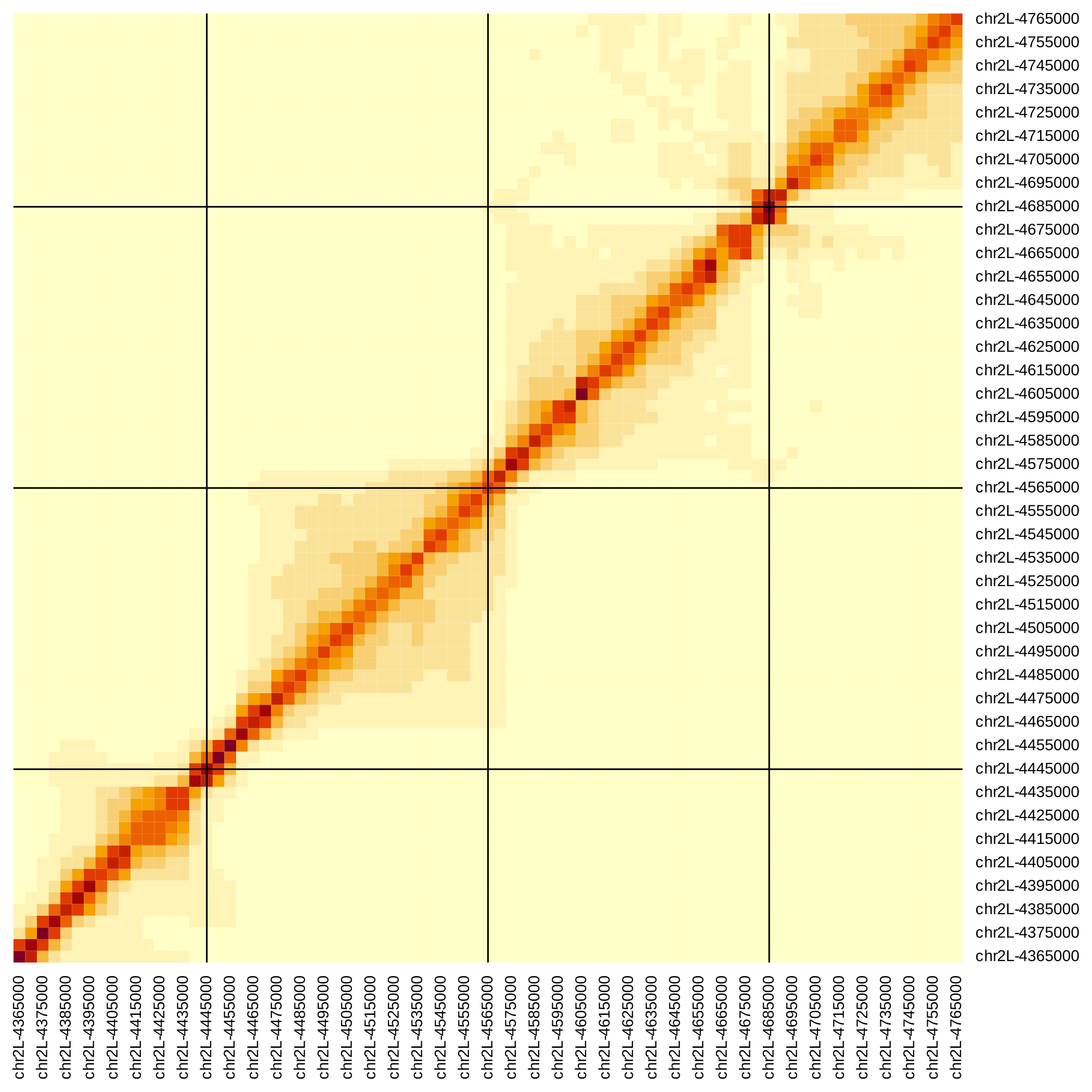

**Figure S3. Local Hi-C heatmap for *D. melanogaster.*** Shown above is the local chromosomal configuration of chromosome 2L in the vicinity of *HP6/Umbrea* (chr2L:4570000, center).

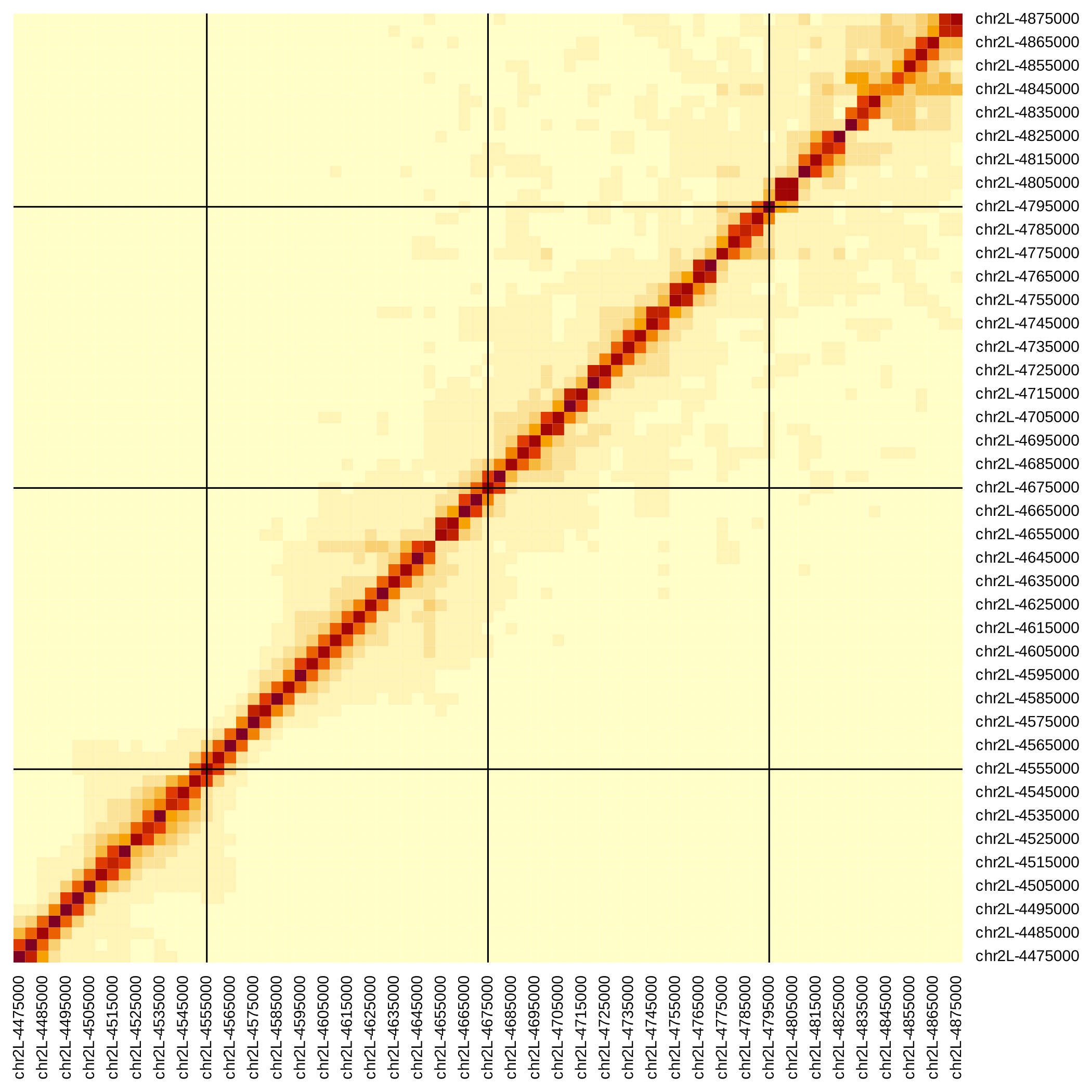

**Figure S4. Local Hi-C heatmap for *D. yakuba****.* Shown above is the local chromosomal configuration of chromosome 2L in the vicinity of *HP6/Umbrea* (chr2L:4680000, center).

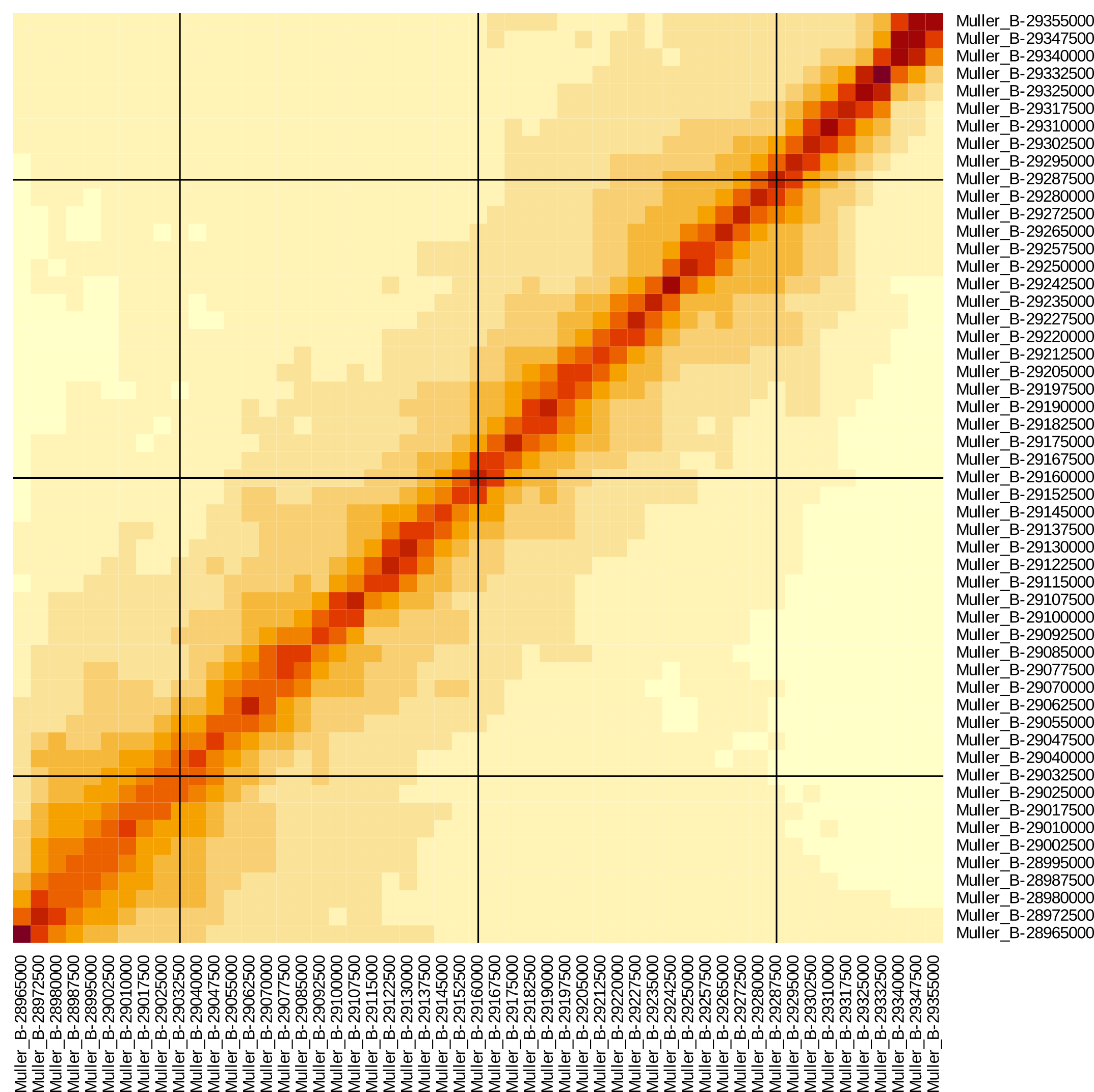

**Figure S5. Local Hi-C heatmap for *D. pseudoobscura*.** Shown above is the local chromosomal configuration of Muller Element B in the vicinity of *HP6/Umbrea*’s future insertion site (Muller B:29165000, center). Note a large-scale chromosomal inversion event occurred between *D.* *melanogaster* and *D.* *pseudoobscura* (not shown).

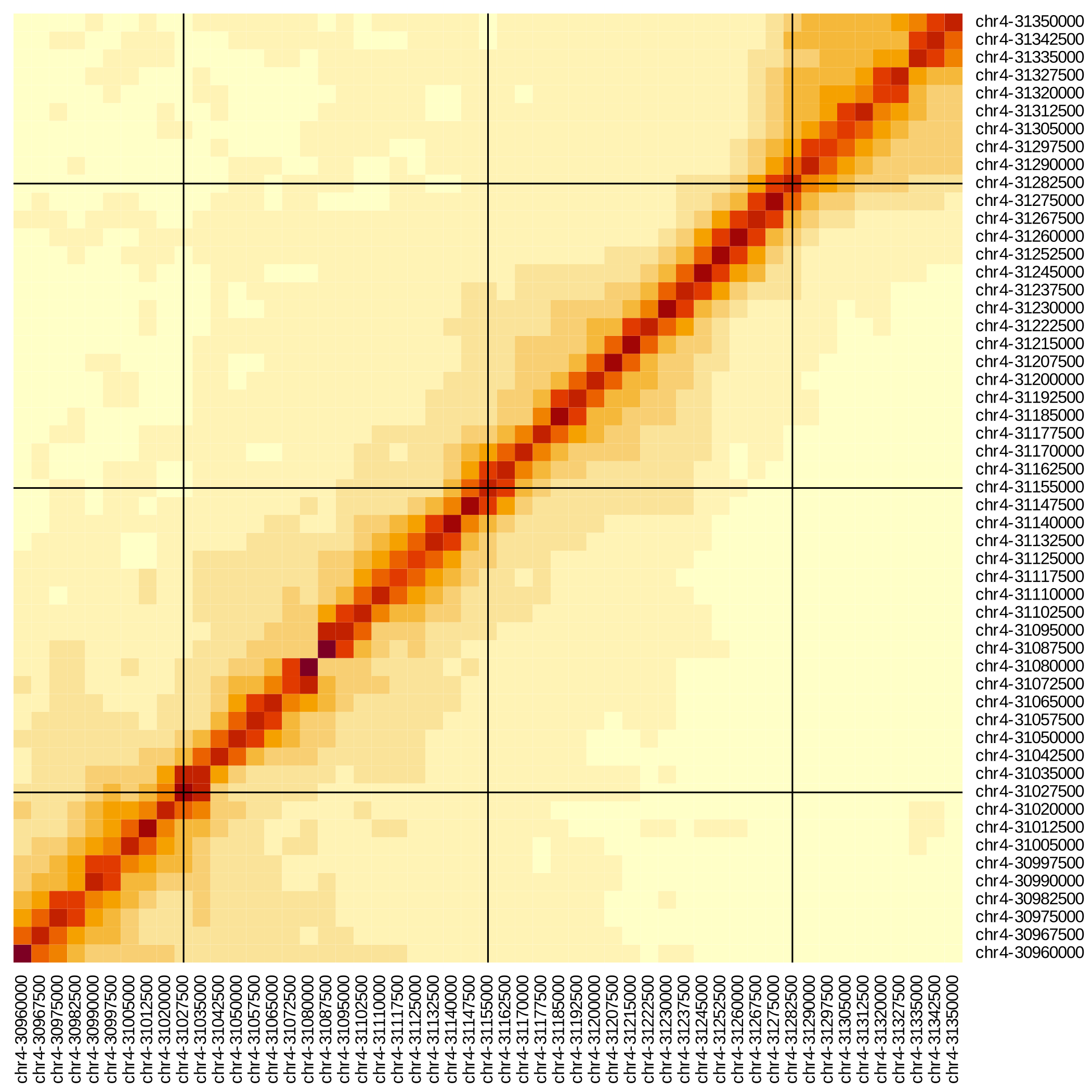

**Figure S6. Local Hi-C heatmap for *D. miranda*.** Shown above is the local chromosomal configuration of chromosome 4 in the vicinity of *HP6/Umbrea*’s future insertion site (chr4:31160000, center).

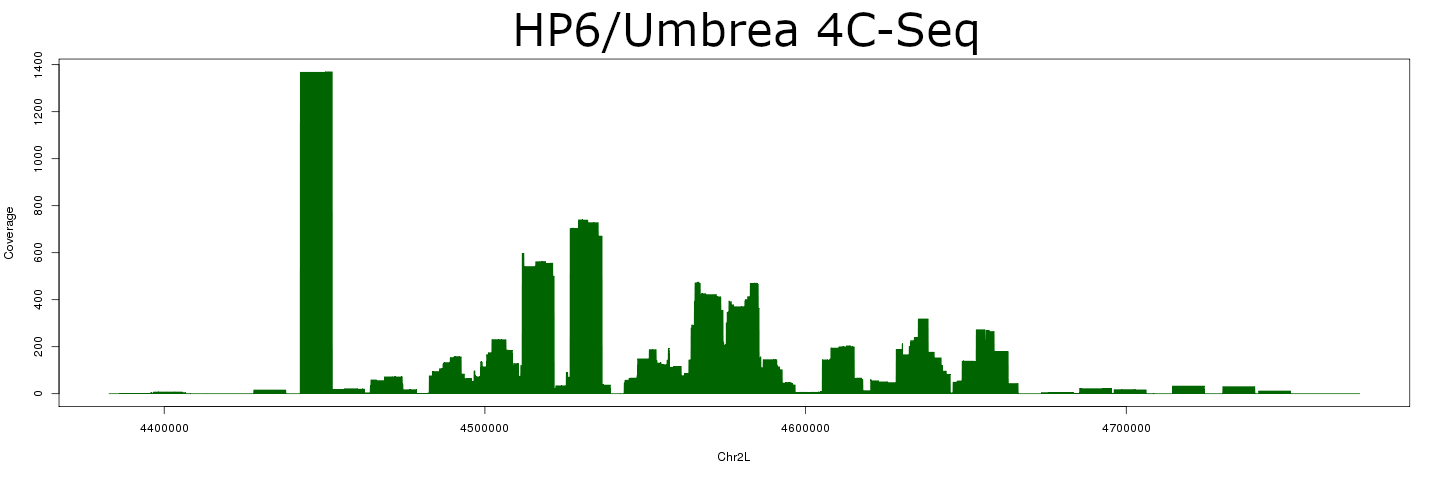

**Figure S7. 4C-Seq in *D. melanogaster*.** Shown above are raw read coverage results from 4C-Seq derived from *D. melanogaster* larval tissue with self-self interactions removed, centered on *HP6/Umbrea*. The strong peak on the left shows the location of FLEE1.

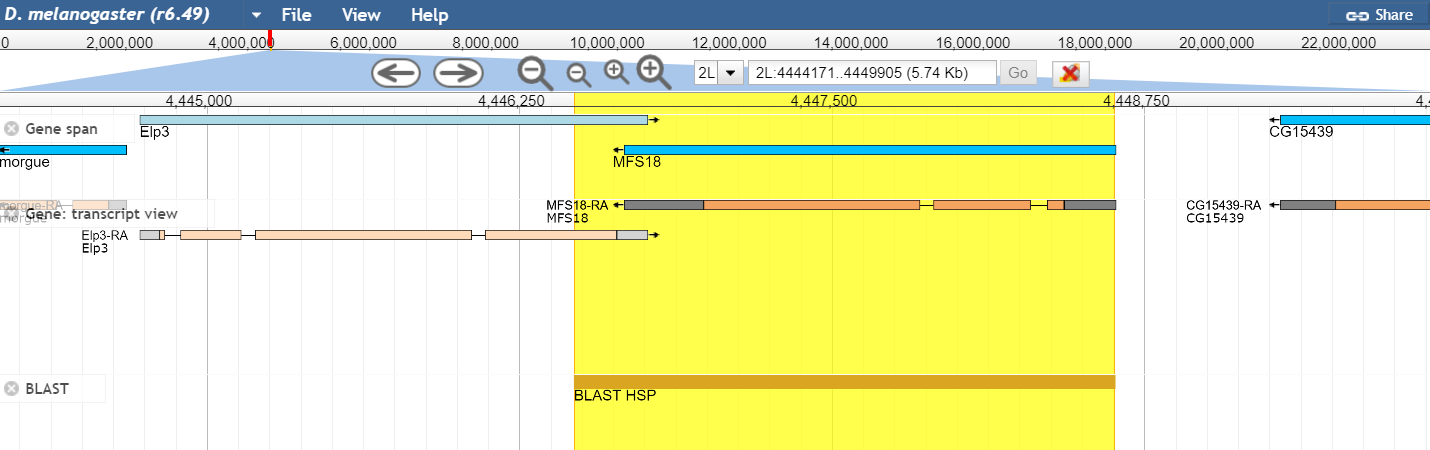

**Figure S8. FLEE1 is located within exonic sequence.** The FlyBase gene track and transcript view for FLEE1 shows that it is contained nearly entirely within the coding sequences of *MFS18* and *Elp3* on chromosome 2L.

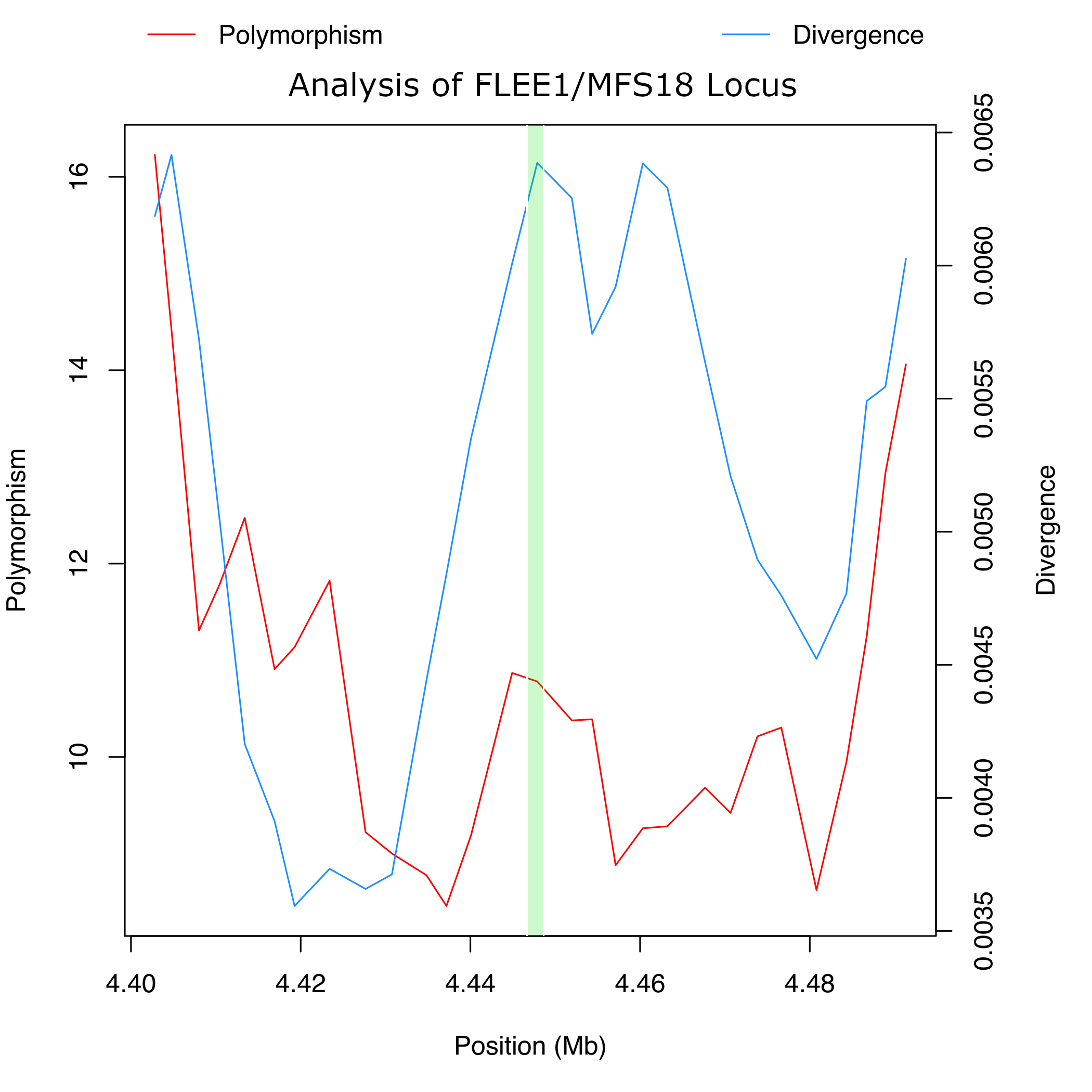
**Figure S9. Polymorphism and divergence in FLEE1/*MFS18* locus.** Polymorphism and divergence calculations on chromosome 2L with a 2.5kb window, with the *MFS18* locus highlighted in green, show how the relative number of polymorphisms is low when compared to sequence divergence.

**File S1. Sequence for FLEE1 enhancer.** Sequence for the cloned FLEE1 element in FASTA format.

>FLEE1

ACCAGGGCTTCGGTATGTTGCTGATGGAGGAGGCGGAGCGAATTGCTCGAGAGGAGCACGGCAGCACAAAACTGGCGGTCATATCGGGAGTGGGCACCAGAAACTACTATCGCAAAATGGGATACCAACTTGACGGACCCTACATGTCAAAGAGCATAGAAGAAAATAACTAGGTATAGCGTTAAATGACTGTCTTGGTGGTATGTTGGAGGATTAAATATTTGTATTTTATCACGCTAGGAGCAGTAAGATTTCGCTACTTAAAACTACTCTCTTAAATATATACATTAATATATAGAATGAATCGATTTATTGGCTAAAACTCACAGGGTCCTTTAAAGTATCAATGATACCACATATTTTTTGGCTTTAACATCTCACAAGAACAACTAAATGATCGTCAATCATAAACGTGTATACTAAATAATATAAGCAGCATGAACATAAATCGATCCACTCCAATATACCCCACACATAAATAAATAGGTTAGTTTTTCTGAGGAAAGTGTGCAAGAGAATGTTAAACGATGGCTTCCGCCGAACCAAAGACTATAAATATGATCCAGCCAACCAAATTGATGCCAGCAGCGGCGCTGAACACCATCGGCCAGCTTTGTGTGAGCTCCAGAATGTGTCCGGCCAAGTATACTCCGAGAAAGCCAGGAATCGCGCCCACTGTGTTCATCAGGCCAAAGACGCTGCCCGAATGCAGAGGTGCCAGGTCTTGGGGATTCACTGTTACCGCGTTGTTGTGGAAGCCCGTGCCGCCAATGATAATGGTCATGCAGATGAGCGCCGTATGGAAGTCCGAGGTGCGGCTCATCACAAACAGGGCCAGATTCTGAGCGGCAAAGCAGCAACTTTGGATGACCTTGCGCACCGTCGTCGTGTGCCATTCGCGAGCGAGTAATCTGGTGGTCAAGTACTTGGCGAATAGCGTGCACGGTGGCAGGGCAAGCCACGGGATCATGTTCACTACCCAACCCTTGGCGTGTGGAAAGCCGTCGTGGAAGTATGTAGGCAGCCAGGAGAGTAGCACGAAGAAGCAGTTCATCTCGCAGGCGTGAGTCAGCACACAGGCCCAGAAGGACAGCCTACGAAAGTATCGCAACCAAGGCACGGCTGACGTCTCTGCCGGACTCTTGTTCGCGCACAGTCGGGATGGCGTGGCAATATTAATGATTCGGTTTCGCTCGCCGGCCATTGCATAGTAGCGCAGCACCAGCGCCCATGCGATGCCCATCAGTCCTATCACCCGGAATACATACGACCAGCCGAAGTAGTCCAGCAGAAAAGATCCCATAATCCCAGTCAGAAGAGTACCTAGAGCCGATCCCGCTGTGAGCAGCCCAAAGAAGCTGCTTCTCTCATTGGGGCACAAATTCTGCAAACGATTAAGTTATAGTTTATGTGTAAATTTATAAAATTAGCTAAGCACCTGACTGGTTAGACTAATCATGCTAGGAAAGTGCACGCCCTGAAGAGCGCCGTTCAGGATTCGAATGGCAACAATGAAGGGAATAGCGTAACTCTTGATGGAGCCCGCCGTCCAGATGATAGTGGGCATTAGGAATGTGATAAGCGACCAGCCGATTGCGGCAAACAGAATGACTCGCTGGCCTCCAAAGCGGTCGCTGAAGTAGCCGCCCACAACCTGCGTGAGTGTGTAGCCCCAGAAGAAGGAGCTGAGCACAGTGCCCGAGTCGGTTTTGCTCCACTTTTGGGCGGATGCCACGGCCGGCACAAGAAGTGGCATAGTGGTGCGGGTGGAGTACAGCATACAGGTGCCCGTAATAAGGGTGATGAACCAGACACGCTTCTCATGCCTGCAATGATCCGACAAAGGAGTTGTACTTGGGGAGGTTTAGTGAGCTGTATGCTGTGAGACCCACCTGGTCCAAATGCTCTGCGTGTCCACCAGTTCCCCGCGCAGCAGAGAATATTTTAGCTTCTCGTCCATGGTCACAAACTGGGTCCGGAACTATTGCCTTTCCTTCACGTCACATATCAACTCCAACTGCTTCGTTGCTTGCCGCTGTGGCATATTTTACTGCCCTTTGTTTACTTTCATTCACGTTGGCGACTAGACACGCCAAGTATTTGCGCCTGTTAAAATTATGTTTTTACGTGGCCGTTTTTCCAACAGCCGCTGGACTAGAGCATAG

**Table S1. Co-expression and essentiality data for newly evolved distal duplicates.** Data for each new gene/parent gene pair with parental coexpression, maximal neighboring coexpression, and essentiality data reported.

| new_gene | par_gene | neigh_cor | par_cor | essentiality |
| --- | --- | --- | --- | --- |
| CG31875 | CG33525 | 0.544016 | 0.418772 | essential |
| CG33458 | CG30090 | 0.510544 | 0.061535 | essential |
| CG33459 | CG30090 | 0.510544 | 0.170643 | essential |
| CG5372 | CG8095 | 0.533216 | 0.100566 | essential |
| CG3347 | CG17440 | 0.690522 | 0.917109 | essential |
| CG31313 | CG8066 | 0.262423 | 0.262423 | essential |
| CG17802 | CG17806 | 0.754936 | 0.754936 | essential |
| CG30395 | CG15040 | 0.564165 | 0.503847 | essential |
| CG8664 | CG5107 | 0.739614 | -0.02909 | essential |
| CG4477 | CG18223 | 0.991723 | 0.87925 | essential |
| CG4907 | CG13978 | 0.473646 | 0.90807 | essential |
| CG31509 | CG31508 | 0.568805 | 0.351047 | essential |
| CG33109 | CG16826 | 0.863108 | 0.855139 | essential |
| CG5609 | CG31508 | 0.568805 | 0.025656 | essential |
| CG4259 | CG3117 | 0.310205 | 0.370675 | essential |
| CG6289 | CG6663 | 0.896926 | 0.896926 | essential |
| CG1736 | CG9327 | 0.476144 | 0.31898 | essential |
| CG2826 | CG13686 | 0.611623 | 0.840596 | essential |
| CG10090 | CG14666 | 0.581515 | 0.72728 | essential |
| CG11597 | CG32505 | 0.294694 | 0.434793 | essential |
| CG9720 | CG31524 | 1 | 0.749838 | essential |
| CG9873 | CG9091 | 0.73774 | -0.00342 | essential |
| CG15358 | CG15818 | 0.285515 | -0.17483 | essential |
| CG11466 | CG4486 | 0.755441 | 0.854113 | essential |
| CG8358 | CG5527 | 0.195031 | 0.546997 | essential |
| CG31791 | CG31801 | 0.858564 | 0.726142 | essential |
| CG32588 | CG33252 | 0.770768 | 0.809965 | essential |
| CG15527 | CG2998 | 0.546119 | 0.011347 | essential |
| CG13686 | CG2839 | 0.727607 | 0.652654 | essential |
| CG17011 | CG17799 | 0.473299 | 0.954925 | essential |
| CG6687 | CG18525 | -0.11768 | -0.11768 | essential |
| CG17012 | CG30031 | 0.999027 | -0.12837 | essential |
| CG30036 | CG33145 | 0.995132 | 0.12326 | essential |
| CG30037 | CG33145 | 0.995132 | 0.083991 | essential |
| CG18125 | CG15040 | -0.12326 | -0.30413 | non_essential |
| CG32368 | CG18754 | 0.049468 | -0.02339 | non_essential |
| CG16992 | CG16761 | -0.05143 | -0.22744 | non_essential |
| CG12224 | CG6392 | 0.876585 | -0.40969 | non_essential |
| CG4580 | CG7052 | 0.23011 | 0.210476 | non_essential |
| CG17650 | CG17945 | 0.937531 | 0.744143 | non_essential |
| CG17637 | CG7754 | 0.769495 | 0.589164 | non_essential |
| CG13463 | CG33489 | 0.63252 | 0.798583 | non_essential |
| CG4712 | CG17734 | 0.852126 | -0.14781 | non_essential |
| CG7815 | CG6663 | 0.795447 | 0.326526 | non_essential |
| CG33490 | CG8622 | 0.774678 | 0.288208 | non_essential |
| CG3640 | CG33101 | 0.622135 | 0.3315 | non_essential |
| CG10799 | CG5784 | 0.639334 | -0.58441 | non_essential |
| CG5509 | CG11958 | 0.546399 | 0.268802 | non_essential |
| CG33920 | CG9091 | 0.512647 | 0.116871 | non_essential |
| CG8626 | CG8095 | 0.958655 | 0.224197 | non_essential |
| CG17673 | CG33104 | 0.947211 | 0.191063 | non_essential |
| CG10232 | CG14213 | 0.122907 | 0.160009 | non_essential |
| CG18748 | CG17799 | 0.914127 | 0.711878 | non_essential |
| CG6036 | CG5724 | 0.943401 | 0.162411 | non_essential |
| CG3217 | CG4842 | 0.753369 | -0.22816 | non_essential |
| CG12842 | CG18249 | 0.722673 | -0.17571 | non_essential |
| CG31769 | CG4486 | 0.906835 | 0.57304 | non_essential |
| CG5265 | CG15503 | 0.486472 | 0.194038 | non_essential |
| CG31918 | CG7599 | 0.726024 | -0.07309 | non_essential |
| CG31932 | CG6912 | 0.705203 | -0.30682 | non_essential |
| CG13091 | CG17843 | 0.17007 | 0.685303 | non_essential |
| CG31370 | CG17843 | 0.772457 | -0.24621 | non_essential |
| CG7594 | CG32107 | 0.999657 | -0.00293 | non_essential |
| CG13656 | CG15293 | 0.64774 | 0.569462 | non_essential |
| CG6639 | CG32284 | 0.826954 | -0.04802 | non_essential |
| CG17268 | CG1844 | 0.544297 | 0.05546 | non_essential |
| CG33235 | CG11598 | 0.349626 | 0.556675 | non_essential |
| CG10700 | CG10863 | 0.560563 | 0.022261 | non_essential |
| CG6208 | CG33252 | 0.250033 | 0.129214 | non_essential |
| CG14610 | CG33145 | 0.998236 | 0.043089 | non_essential |
| CG31508 | CG30083 | 0.531385 | 0.346498 | non_essential |
| CG30450 | CG1101 | 0.585754 | -0.32546 | non_essential |
| CG31524 | CG33525 | 1 | 0.198622 | non_essential |
| CG1840 | CG5983 | 0.447979 | 0.301494 | non_essential |
| CG13977 | CG30362 | 0.680651 | 0.590968 | non_essential |
| CG6690 | CG1906 | 0.523537 | 0.023827 | non_essential |
| CG30473 | CG1718 | 0.760268 | 0.361607 | non_essential |
| CG8856 | CG8036 | 0.652693 | 0.330431 | non_essential |
| CG17174 | CG3422 | 0.909234 | 0.14695 | non_essential |
| CG17176 | CG4199 | 0.923376 | 0.114519 | non_essential |
| CG12493 | CG31508 | 0.642318 | -0.1908 | non_essential |
| CG11833 | CG31013 | 0.057125 | 0.015086 | non_essential |
| CG7931 | CG12359 | 0.960604 | -0.19223 | non_essential |
| CG30494 | CG8588 | 0.818025 | 0.676018 | non_essential |
| CG15461 | CG5302 | 0.045446 | 0.361934 | non_essential |
| CG33462 | CG1942 | 0.762024 | -0.09881 | non_essential |
| CG15636 | CG7041 | 0.818182 | 0.087641 | non_essential |

**Movie S1. Live GFP expression.** Video of live larvae with FLEE1 under the control of an enhancer-reporter vector. Expression is seen restricted to the larval salivary glands. (MovieS1.mp4)
